# Design approaches to expand the toolkit for building cotranscriptionally encoded RNA strand displacement circuits

**DOI:** 10.1101/2023.02.01.526534

**Authors:** Samuel W. Schaffter, Molly E. Wintenberg, Terence M. Murphy, Elizabeth A. Strychalski

**Affiliations:** National Institute of Standards of Technology, Gaithersburg, MD 20899, USA

## Abstract

Cotranscriptionally encoded RNA strand displacement (ctRSD) circuits are an emerging tool for programmable molecular computation with potential applications spanning *in vitro* diagnostics to continuous computation inside living cells. In ctRSD circuits, RNA strand displacement components are continuously produced together *via* transcription. These RNA components can be rationally programmed through base pairing interactions to execute logic and signaling cascades. However, the small number of ctRSD components characterized to date limits circuit size and capabilities. Here, we characterize 220 ctRSD gate sequences, exploring different input, output, and toehold sequences and changes to other design parameters, including domain lengths, ribozyme sequences, and the order in which gate strands are transcribed. This characterization provides a library of sequence domains for engineering ctRSD components, *i.e*., a toolkit, enabling circuits with up to four-fold more inputs than previously possible. We also identify specific failure modes and systematically develop design approaches that reduce the likelihood of failure across different gate sequences. Lastly, we show ctRSD gate design is robust to changes in transcriptional encoding, opening a broad design space for applications in more complex environments. Together, these results deliver an expanded toolkit and design approaches for building ctRSD circuits that will dramatically extend capabilities and potential applications.

## Introduction

Nucleic acid-based circuits, *i.e*., networks of interacting nucleic acids programmed to process molecular information, are an increasingly useful tool for molecular computation with applications spanning *in vitro* diagnostics^1–5^ and biosensing^6,7^ to synthetic cells^8,9^ and cellular computation^10–13^. Nucleic acids are an ideal substrate for building molecular circuits because predictable base pairing rules facilitate the rational design of programmable interactions. Compared to transcription factor-based cascades^14^, nucleic acid circuits can also operate with faster response times and lower energetic costs^15,16^. Many nucleic acid circuits operate *via* toehold-mediated strand displacement (TMSD), in which a single-stranded toehold domain of a nucleic acid duplex or hairpin recruits a sequence complementary input strand to initiate strand displacement and expose a new domain that enacts a downstream response^17^. As a testament to the programmability, modularity, and scalability of TMSD reactions, the field of DNA computing has demonstrated *in vitro* TMSD circuits composed of tens to hundreds of components programmed to execute computational and information processing tasks, such as digital calculations^18^ and pattern recognition^19^.

Cotranscriptionally encoded RNA strand displacement (ctRSD) circuits^20^ are an emerging technology, modeled after state-of-the-art, DNA-based TMSD circuits^18,19^, with the potential for implementation across many synthetic biology platforms. In ctRSD circuits, DNA templates are transcribed to produce single-stranded RNA (ssRNA) inputs and double-stranded RNA (dsRNA) gates. Gates are initially transcribed as ssRNA hairpins that cleave *via* an internal self-cleaving ribozyme after gate folding to produce a dsRNA complex (Figure 1A,B). This allows multiple gates with complementary domains to be produced together without prominent cross-reaction.

**Figure 1:**
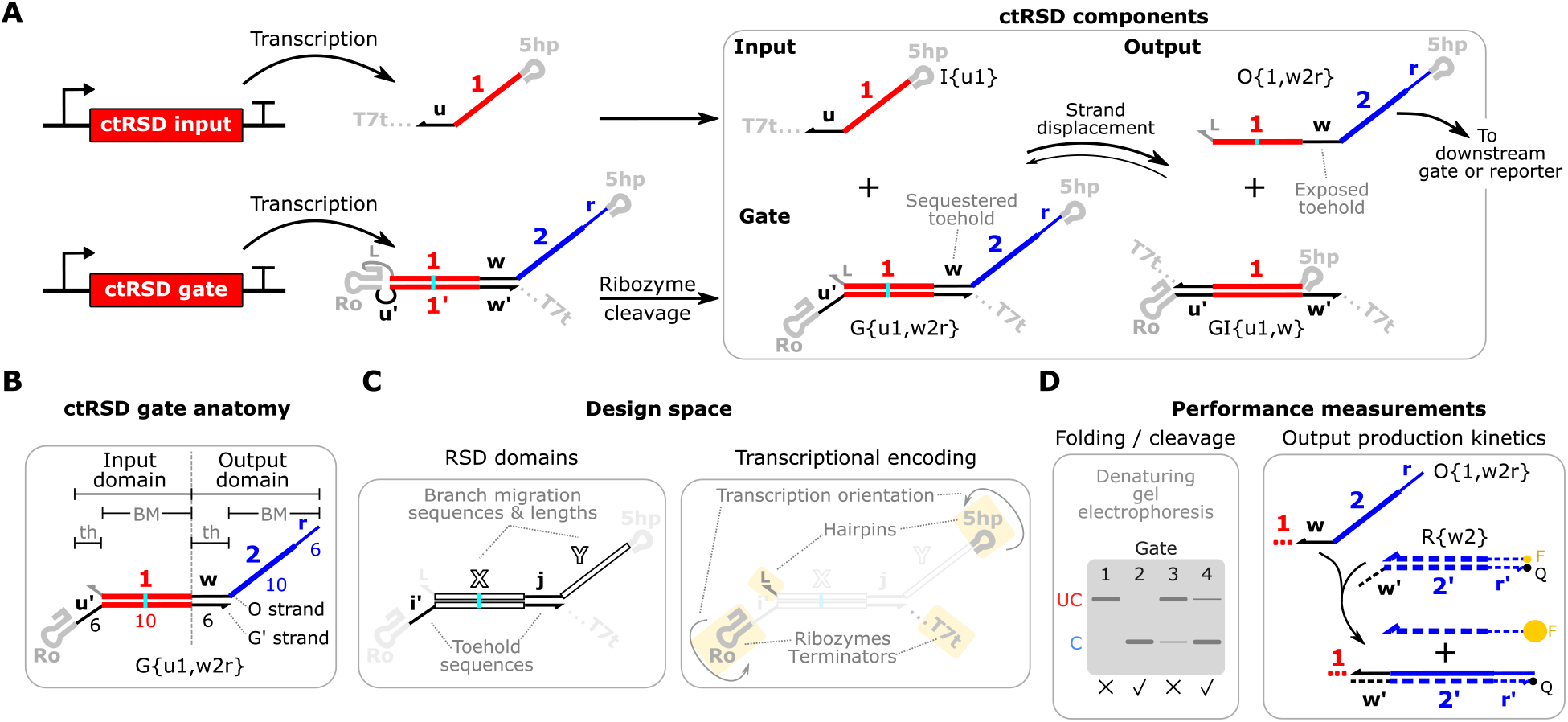
Cotranscriptionally encoded RNA strand displacement (ctRSD) circuit components, nomenclature, and measurements. (**A**) ctRSD components and reactions. Inputs and gates are transcribed from dsDNA templates. An internal self-cleaving ribozyme (Ro) cleaves ctRSD gates to produce dsRNA components that can undergo strand displacement with ssRNA inputs to release ssRNA outputs. Gates are designed with an internal G-U wobble base pair (cyan), which is replaced with a G-C base pair by the input in the gate-input complex (GI) to drive the forward reaction. All gates and inputs have a 5′ hairpin and a 3′ terminator hairpin. These domains are omitted for brevity in subsequent figures. (**B**) ctRSD gates are composed of an output strand (O strand) and a strand complementary to a portion of the output strand (G′ strand). Gates are defined by their input and output domains, each composed of a toehold (th) subdomain and a branch migration (BM) subdomain. Bold numbers and letters above the gate represent domain sequence identity, and nonbold numbers below the gate represent domain lengths in bases. Gates are designated as G{input domain, output domain}. Inputs and outputs follow a similar nomenclature, designated by I{} and O{}, respectively. (**C**) The designs explored in this study. Changes were made to domains relevant to either RNA strand displacement (RSD) or to gate transcriptional encoding (highlighted in yellow). (**D**) Gate performance was characterized with two assays, one that measures gate cleavage (denaturing gel electrophoresis) and another that measures output production (fluorescence-based DNA reporter assay). Note the *r* domain present in the output domain of gates connects to the reporter to facilitate an irreversible reporting reaction. See Supporting Note 3 for additional gate anatomy and sequence schematics.

After ribozyme cleavage, gates react with sequence complementary inputs *via* TMSD to produce ssRNA outputs. Strand displacement exposes a toehold domain on the outputs that can initiate downstream strand displacement reactions with other ctRSD gates or reporters (Figure 1A). Based on these principles, ctRSD circuits have been programmed to execute logic operations and multilayer signaling cascades with predictable kinetics in *in vitro* transcription reactions^20^. Because all ctRSD circuit components are produced together *via* transcription, these circuits could be genetically encoded for continuous operation in living cells, cell lysates, or biological samples, where nucleases would eventually deplete TMSD components added at fixed concentrations^13,21,22^. In these environments, ctRSD circuits could be programmed to sense changing patterns of native nucleic acid sequences in real-time and, in response, regulate downstream gene expression^11,13,23^.

Although ctRSD circuits have potential relevance to broad applications in synthetic biology, the design space for ctRSD components has not been extensively explored. Only five input-output sequences have been tested to date; multi-input information processing for biologically relevant patterns will require at least two- to three-fold more sequences^1,2^. Further, changes to the gate transcriptional encoding parameters have not been investigated, *e.g*., the ribozyme, terminator, and hairpin sequences added to gates for effective transcription. Modulation of transcriptional encoding domains could tune RNA stability in environments with nucleases^24–26^ and could improve the performance of certain gates^27^. However, it is difficult to predict how changes in sequence will influence performance because ctRSD gate formation is governed by out-of-equilibrium cotranscriptional folding. Even minor sequence changes that do not appreciably change the minimum free energy structure can induce misfolds during transcription that disrupt gate functionality^28,29^. Creating a library of functional sequence domains with which to construct many ctRSD components, *i.e*., a toolkit, as well as identifying design approaches that yield components that perform as expected across many combinations of these sequence domains, would greatly extend the capabilities of this technology.

Here, we expand the toolkit for building ctRSD circuits by characterizing the performance of 220 gates *in vitro*, exploring changes to domains relevant to both strand displacement and transcriptional encoding (Figure 1C). We find the initial ctRSD gate design is readily scalable, and we develop a set of 17 unique input-output sequences. The design is amenable to other alterations, such as changes in branch migration length, transcriptional encoding domains, and RNA folding path, which could improve performance in complex environments. However, not all gates function as designed and those that do not work exhibit diverse failure modes. So, we systematically develop design approaches to reduce the likelihood of failure across gate sequence contexts and recover the performance of failed gates. These design approaches should enable straightforward expansion of the toolkit for building ctRSD circuits with a high chance that new gate sequences perform as expected. Together, our results provide an extensive library of ctRSD sequence domains with which multicomponent circuits can be quickly and reliably designed, implemented, and adopted to different applications.

## Results and Discussion

### Overview of gate performance measurements and metrics

To assess the performance of each ctRSD gate, we conducted measurements of ribozyme cleavage, output production from RNA strand displacement between the gate and its designed input, and output production from the gate in the absence of its designed input, *i.e*., leak. We characterized ribozyme cleavage using denaturing gel electrophoresis, assuming gates that cleaved poorly were not folded correctly. We characterized output production from RNA strand displacement and leak with a fluorescence-based DNA reporter assay (Figure 1D). For each gate we evaluated, we conducted the DNA reporter assay with both the designed input to characterize output production from RNA strand displacement and a scrambled input (Io) to measure leak.

To classify a specific gate sequence as having either expected or poor performance based on our measurements, we developed the following metrics. For ribozyme cleavage, we deemed gates that cleaved ≤50% under our assay conditions as having poor performance. Gates with expected performance typically cleaved >90%. To determine a metric for expected RNA strand displacement between a gate and its input, we compared experimental results to a kinetic model of ctRSD reactions (Supporting Note 2). Based on previous literature^30,31^, we expected the RNA strand displacement rate constant (k_rsd_), the rate constant that governs the reaction between a gate and its input, to differ up to an order of magnitude across different input sequences. Therefore, we used kinetic simulations to assess whether the reporter kinetics of a gate transcribed with its input were consistent with a k_rsd_ value up to 5-fold higher or lower than the expected value measured previously^20^, *i.e*., k_rsd_ spanning an order of magnitude from the fastest to slowest sequences. Sequence-specific differences in other reaction rate constants, such as transcription and ribozyme cleavage, may also influence reporter kinetics, but k_rsd_ captures the salient effects of these differences in a single parameter (Supporting Note 2). Regarding leak, *i.e*., output production in the absence of input, our previous results suggest this arises from truncated or misfolded gate transcripts that react directly with downstream components and are produced at (3 to 6) % the rate of correctly folded gate transcripts^20^. Because excessive leak interferes with building multilayer cascades, we considered leak of ≥10% of the gate’s transcription rate as the cutoff for poor performance; this corresponds to >50% reacted reporter in three hours for a gate.

*Throughout this study, the above metrics for ribozyme cleavage, RNA strand displacement, and leak served as criteria for characterizing gates as having either expected or poor performance*. Components with performance outside these expected ranges likely have a large population of misfolded transcripts (Figure S1), making them unreliable for engineering circuits.

### Characterizing more sequence contexts of the initial ctRSD gate design reveals failure modes

The initial ctRSD gate design described previously^20^ included the domain lengths specified in Figure 1B and the following four design choices. First, the gates used a minimal version of the anti-genomic HDV ribozyme sequence^32^ (Ro) (Supporting Note 3). Second, the single-stranded domains of input and output strands were restricted to C, A, or U bases, to reduce undesired secondary structure and cross hybridization of single-stranded components. Third, based on the sequence constraints above, the output strand of the gate was transcribed first to mitigate misfolding in the nascent transcript. Fourth, the input branch migration domain of the gates contained a G-U wobble base pair (cyan in Figure 1) that becomes a G-C base pair when the input binds to favor the forward strand displacement reaction. Here, we first investigated how this initial gate design performed with additional sequences and sequence contexts.

We began by evaluating the performance of gates with 16 unique input sequences and a previously-characterized output domain, domain *2*^20^. We refer to these gates as G{uX,w2r}, where *X* corresponds to different input branch migration domains. Note, domain *2* was exclusively an output domain in this study (Figure 2A). The 16 input sequences were derived from a previous DNA computing study^18^ with slight modifications to follow the initial ctRSD gate design. tested each gate in the DNA reporter assay with its designed input I{uX} and a scrambled input Io. As expected, reporter kinetics differed slightly across input sequences but were consistent with rate constants for RNA strand displacement spanning a factor of eight for all but G{u11,w2r}, *i.e*. all but one gate had reporter kinetics within the expected range from simulations (Figure 2A). We confirmed relative differences in reporter kinetics across gates with different input domains and the same output domain were reproducible in both technical and biological replicates (Methods, Supporting Note 4). Other than G{u11,w2r}, the gates cleaved well (Figure 2B, top). Sequences upstream of the ribozyme can interfere with folding and cleavage^27^, so the specific sequence of G{u11,w2r} may have caused the ribozyme to misfold during transcription (Figure S1).

**Figure 2:**
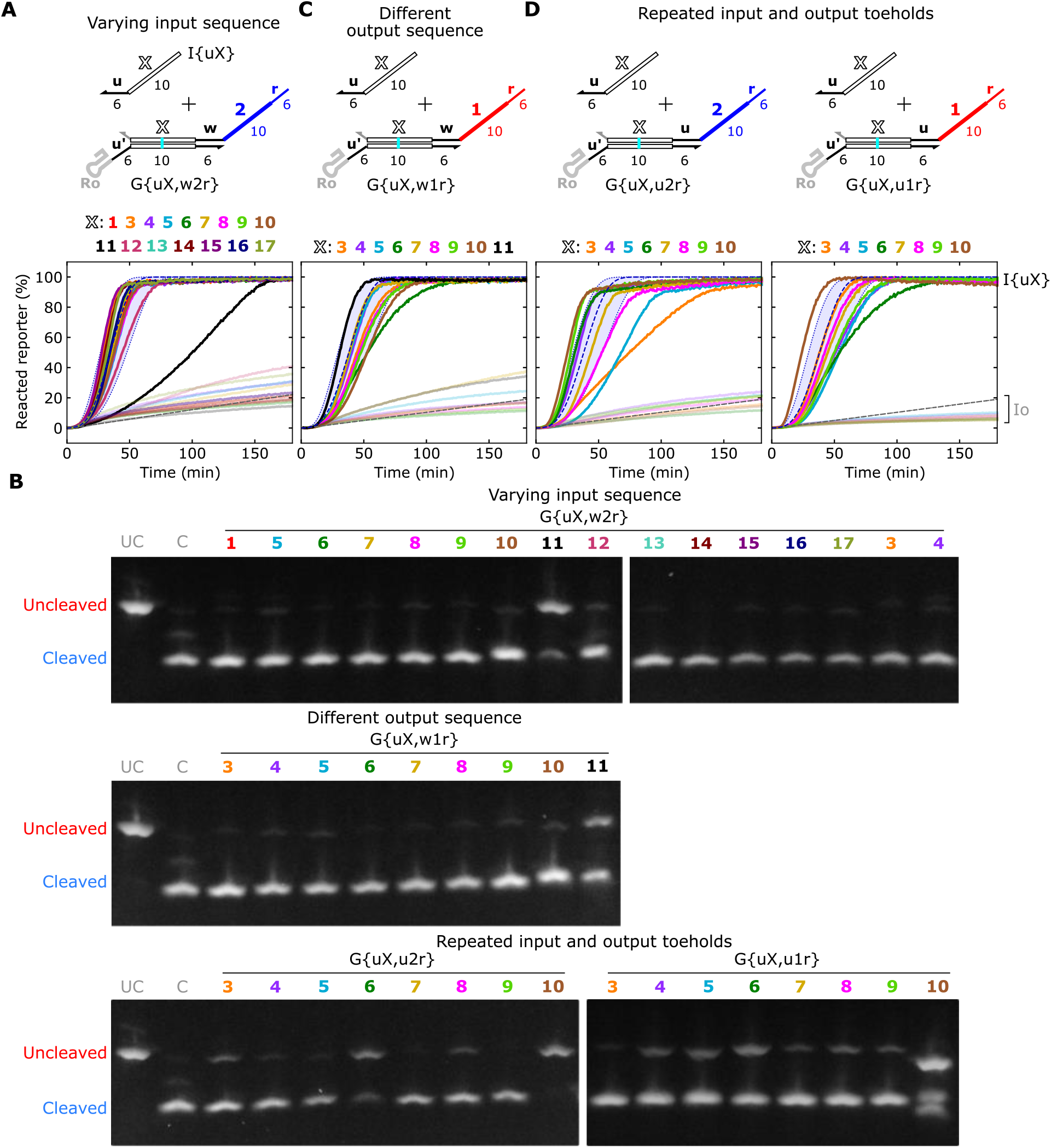
Characterization of ctRSD gates with different input and output sequences. (**A,C,D**) Gates with plots of their reporter kinetics. The white *X* in the schematics denotes the varied domain of the gate. Line colors in the plots correspond to the domain number colors above the plots. In plots, full color lines indicate gates transcribed with their designed input (I{uX}), and semi-transparent lines indicate gates transcribed with a scrambled input (Io). Dashed blue lines represent simulation results with k_rsd_ = 10^3^ L·mol^-1^s^-1^. The blue shaded region between dotted lines represents simulations spanning 4k_rsd_ to k_rsd_/4. Experiments for each plot used a different DNA reporter: R{w2} (A), R{w1} (B), R{u2} (D, left), R{u1} (D, right). See Supporting Note 3 for reporter sequence schematics. Gate and input templates were present at 25 nmol/L and 50 nmol/L, respectively. Reporters were present at 500 nmol/L. (**B**) Denaturing gel electrophoresis results of the gates indicated above the gels. UC is an uncleaved gate size marker, which is G{u1,w2r} with a single base mutation in Ro that prevents cleavage. C is a cleaved gate size marker, which is the G′ strand of G{u1,w2r}, the largest product of a cleaved gate. See Supporting Note 6 for individual kinetic plots.

To explore a different sequence context, we changed the output branch migration domain and evaluated performance of G{uX,w1r} with a subset of input domains tested in Figure 2A, (Figure 2C). All nine gates tested had reporter kinetics that fell within the expected range from simulations (Figure 2C). Other than G{u11,w1r}, the gates cleaved well (Figure 2B, middle). Interestingly, both G{u11,w2r} and G{u11,w1r} cleaved poorly, but G{u11,w2r} had reporter kinetics that were much slower than expected, while G{u11,w1r} was within the expected range. These results suggest the two gates adopted different misfolded structures (Figure S1).

To investigate the influence of input branch migration length on gate performance, we tested gates with several extended input branch migration lengths. Previously, 16 base branch migration lengths were used^20^ but this relatively short duplex could dehybridize in environments with low salt concentrations or elevated temperatures. Here, we tested G{u1,w2r} with (18, 20, and 22) base input branch migration lengths and found all gates had the expected performance. We also observed similar performance for gates with extended branch migration domains in a two-layer cascade (Figure S2).

We next evaluated the performance of gates with the *u* toehold in both the input and output domains. Repeating the same input and output toehold sequence is common in DNA strand displacement circuits to promote uniform kinetics and increase circuit composability^18,19,30^, so we investigated whether this design would also work for ctRSD components. We tested G{uX,u2r} and G{uX,u1r} with eight different input domains (Figure 2D). Of the 16 gates tested, six with the *u* toehold in both the input and output domain either cleaved poorly (Figure 2B, bottom) or had reporter kinetics well outside the expected range from simulations (Figure 2D). In contrast, gates with the same input domains and the *w* toehold in the output domain performed as expected (Figure 2A, B).

*These results present an expanded set of input-output sequences for building ctRSD circuits (Figure 7A)*. However, sequence-specific failure modes warrant further investigation, such as why many gates with a *u* output toehold and input domain *u11* have poor performance.

### Differing input and output toehold sequences within gates reduces failure across sequences

Our results in Figure 2 suggest two possible failure mechanisms for gates with the *u* toehold in both the input and output domains. The *u* toehold in the output domain may itself cause poor performance in that context. Alternatively, repeating the same toehold sequence in the input and output domains may compromise performance. We next sought to distinguish between these two possible mechanisms.

We hypothesized that repeating input-output toehold sequences within gates can facilitate gate misfolding during transcription. For example, G{uX,u2r} in Figure 3A has two *u′* domains in the gate sequence; the first one produced during transcription is the input toehold, which could serve as an alternative binding site for the *u* domain of the output strand. This alternative binding could nucleate a larger misfold in the nascent transcript, causing the gate to become kinetically trapped in a structure with diminished cleavage or slower strand displacement. Guided by this potential mechanism, we devised possible misfolded structures, in which the alternative binding of the *u* output toehold induces sequence-specific disruption of ribozyme folding or prevents input binding (Figure 3A, I and II, respectively). Consistent with these possible misfolded structures, disruption of the HDV ribozyme helix adjacent to the input toehold has been shown to reduce cleavage activity^27^.

**Figure 3:**
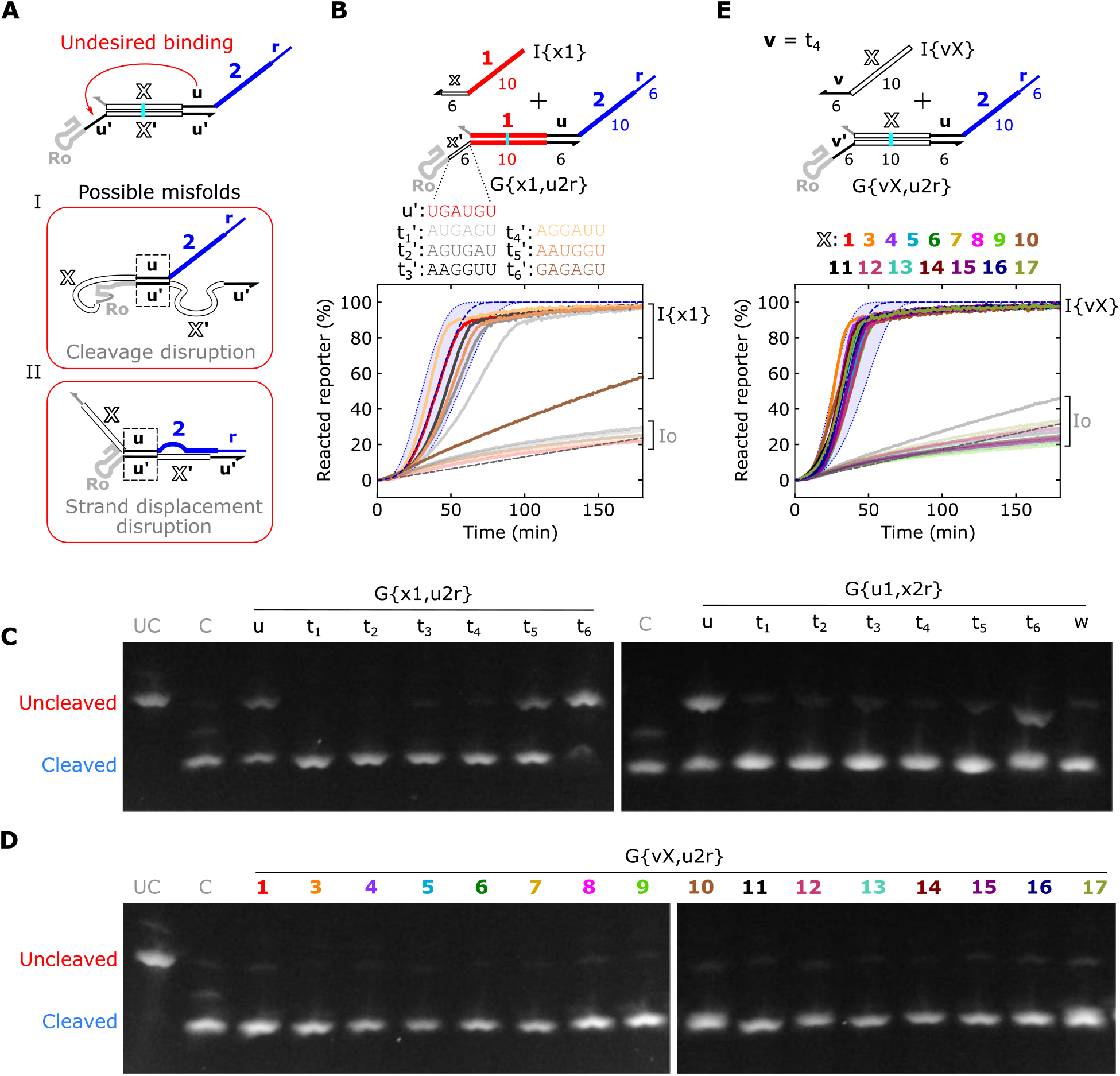
Characterization of ctRSD gates with different toehold sequences. (**A**) Possible misfolds of gates with repeated input and output toeholds. Dashed boxes show the toehold of the output strand binding to the input toehold of the gate instead of the intended output toehold. Depending on the adjacent sequences, this alternative binding could facilitate: (I) disruption of the ribozyme, while leaving enough *X′* unhybridized to allow strand displacement with the input, (II) alternative hybridization with the *X′* domain that precludes strand displacement with the input, or a combination of I and II (Figure S1). (**B,E**) Reporter kinetics of the gates indicated above the plots. The white *X* in the schematics denotes the varied domain of the gate. In (B), line colors in the plot correspond to the toehold sequence colors above the plot. In (E), line colors in the plots correspond to the domain number colors above the plot. Blue dashed lines represent simulation results with the shaded region spanning 4k_rsd_ to k_rsd_/4. Gate and input templates were present at 25 nmol/L. R{u2} was present at 500 nmol/L. (**C,D**) Denaturing gel electrophoresis results of the gates indicated above the gels. UC and C are size markers described in Figure 2. See Supporting Note 6 for individual kinetic plots.

These possible misfolded structures (Figure 3A) are also consistent with our measurements of gates with repeated input-output toeholds (Figure 2D). For example, none of the gates with poor performance had abnormally high leak, indicating the *u* toehold in the output domain was sequestered. G{u6,u2r}, G{u10,u2r}, and G{u10,u1r} had poor cleavage but the expected reporter kinetics, consistent with a disrupted ribozyme and an exposed *X*′ domain that could serve as a site for an input to bind and expose the output toehold *via* strand displacement (Figure 3A, I, Figure S1C). Additionally, G{u3,u2r}, G{u5,u2r}, and G{u6,u1r} cleaved well but had reporter kinetics that were slower than expected, consistent with the possible structure in Figure 3A, II.

If our hypothesis regarding repeating input-output toehold sequences within gates is correct, rather than a *u* output toehold itself compromising performance, we expected changing the input toehold sequence to recover the performance of gates that failed with repeated *u* toeholds. Testing this hypothesis first required an input toehold sequence orthogonal to the *u* sequence and with similar performance. So, we designed and tested gates with six different input toehold sequences (*t_1_* to *t_6_*), each with similar GC content to the *u* toehold to obtain comparable strand displacement kinetics (Figure 3B). The gate with a *t_6_* input toehold had very poor performance (Figure 3B, C), possibly because this toehold had substantial sequence overlap with an adjacent helix of Ro^27^ (Supporting Note 5). The reporter kinetics of the gate with a *t_1_* input toehold were slightly slower than expected from simulations, but the rest of the gates performed as expected (Figure 3B, C). Similar results were observed for select toehold sequences with G{x3,u2r}, G{x6,u2r}, and G{x10,u2r} (Figures S3, S4). The *t_4_* toehold had the best performance across all the gates tested, so we selected it to use alongside the *u* toehold. For convenience, we refer to the *t_4_* toehold sequence as *v* for the remainder of this study.

To test our hypothesis that repeating input-output toehold sequences within gates can cause gates to fail, we measured the performance of gates with a *v* input toehold and a *u* output toehold across 16 different input domains, G{vX,u2r}. All 16 gates cleaved well (Figure 3D) and had reporter kinetics within the expected range from simulations (Figure 3E), indicating that the *u* toehold is not disruptive by itself in the output domain. We also confirmed gates with a different output branch migration domain had the expected performance (G{vX,u1r}, Figure S5). Additionally, the *v* and *w* toeholds were compatible, as G{vX,w2r} and G{vX,w1r} had expected performance across nine different input domains (Figure S6). To further confirm the detrimental effects of repeating input-output toeholds within ctRSD gates, we validated that two gates that had poor cleavage with repeated *u* toeholds also had poor cleavage with repeated *v* toeholds (Figure S7).

*These results indicate that repeating input-output toeholds within ctRSD gates should be avoided. We developed a set of compatible toehold sequences (*u, v, w*) to facilitate the design of gates with alternating toeholds (Figure 7A)*.

### Changing ribozyme sequences can recover the performance of failed gates

Although designing gates with different input and output toehold sequences reduced the chances of poor gate performance across sequence contexts, a few gates with this constraint still performed poorly. For example, G{u11,w2r} and G{u11,w1r} (Figure 2) and G{t61,u2r} had poor ribozyme cleavage (Figure 3). We also found three gates with a *u* input toehold and a *v* output toehold (G{u4,v1}, G{u5,v1}, G{u16,v1}) had poor cleavage and reporter kinetics that were slower than expected (Figure S8).

Because these gates all cleaved poorly, we hypothesized using a different ribozyme sequence may improve performance. Closer analysis of the Ro ribozyme sequence revealed 14 contiguous G, A, U bases spanning the input toehold of the gate and an adjacent helix of Ro, termed the P2 helix (Figure 4A). The output strand of the gate is composed entirely of C, A, U bases and could bind cotranscriptionally to the stretch of G, A, U bases in and around the P2 helix of Ro to disrupt ribozyme folding and cleavage^27^. To reduce this possibility, we designed three ribozyme sequences with mutations in the P2 helix to interrupt the contiguous G, A, U bases (R2, R3, R4 in Figure 4B). We also tested the genomic HDV ribozyme (Rg)^33^ and a ribozyme with an HDV-like fold identified in humans (Rh)^34^, as these have different P2 sequences compared to Ro.

**Figure 4:**
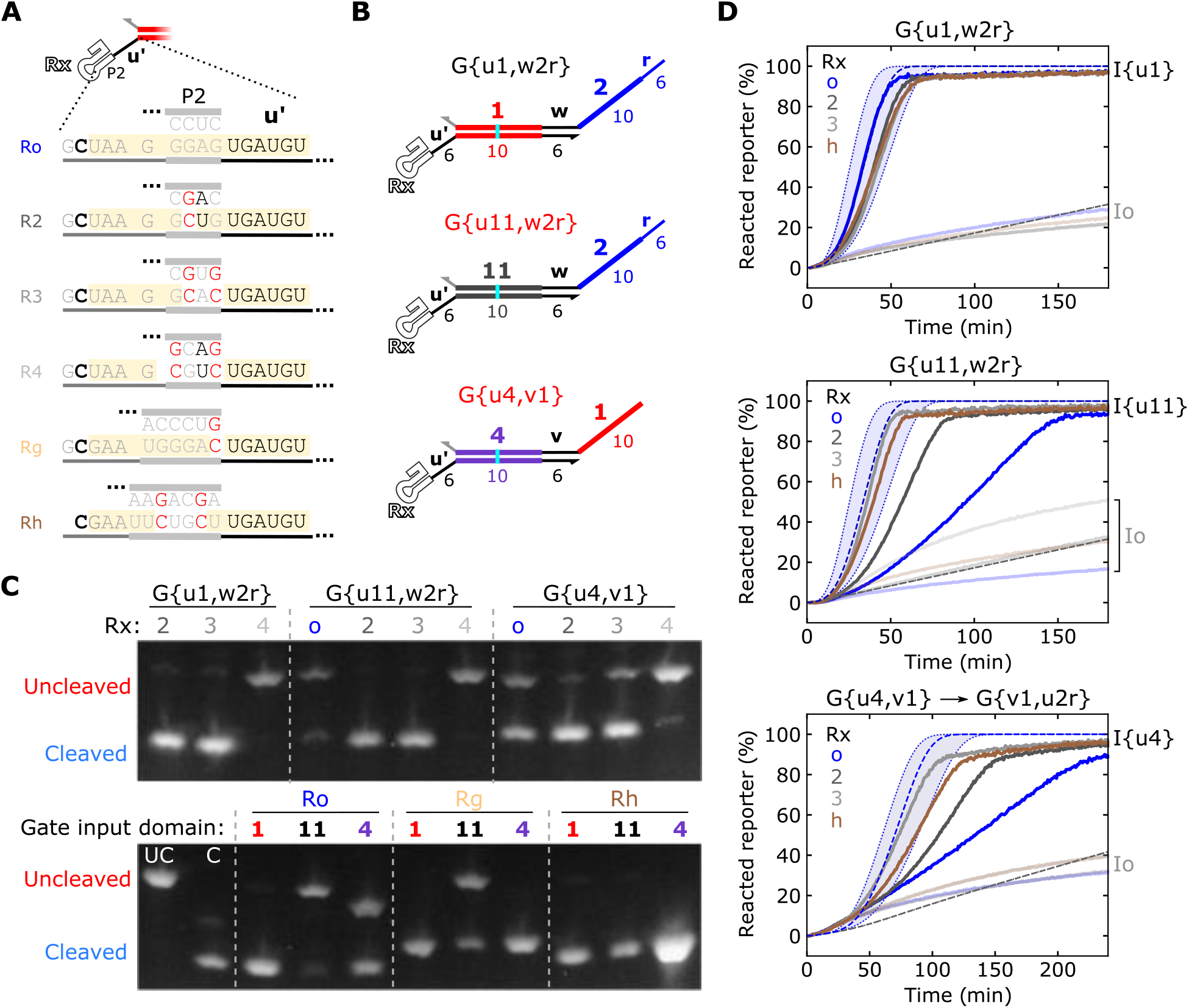
Characterization of ctRSD gates with different ribozyme sequences. (**A**) The P2 helix sequence of ribozymes with flanking sequences. The bold **C** indicates a base necessary for cleavage. The yellow shading highlights regions of contiguous G, A, U bases. Red bases indicate base changes that disrupt contiguous G, A, U bases. (**B**) The gates tested with different ribozyme sequences (white Rx). G{u11,w2r} and G{u4,v1} had poor performance in previous experiments (Figure 2A and Figure S8, respectively). (**C**) Denaturing gel electrophoresis results of the gates indicated above the gels. UC and C are size markers described in Figure 2. (**D**) Reporter kinetics of the gates indicated above the plots. The line colors in the plot correspond to the colors of the Rx labels in the plot. The blue dashed lines represent simulation results with the shaded region spanning 4k_rsd_ to k_rsd_/4 in the top and middle plots and 5k_rsd_ to k_rsd_/5 for G{u4,v1} in the bottom plot. Gate and input templates were present at 25 nmol/L. Reporters were present at 500 nmol/L.

To test the hypothesis that alternative ribozyme sequences can recover gate performance, we evaluated their effect on the performance of three gate sequences (Figure 4B). We tested G{u1,w2r} as a positive cleavage control, given its good performance with Ro in many sequence contexts. We also tested G{u11,w2r} and G{u4,v1}, as these gates performed poorly with Ro. R2 and R3 recovered cleavage activity for G{u11,w2r}, and G{u4,v1}. R4 had poor cleavage for all three gates tested (Figure 4C, top), possibly because this sequence fails to fold correctly in any sequence context. Rg recovered cleavage activity for G{u4,v1} but not G{u11,w2r}, while Rh recovered cleavage activity for both gates (Figure 4C, bottom). Based on these results, we chose R2, R3, and Rh to test further in the DNA reporter assay. For G{u1,w2r}, all these ribozyme sequences had reporter kinetics within the expected range from simulations (Figure 4D). For G{u11,w2r} and G{u4,v1}, all three ribozyme sequences increased reporter kinetics compared to the same gates with Ro, but the rates for R2 were still slower than expected from simulations (Figure 4D). We confirmed the generalizability of these results by validating R3 and Rh recovered the performance of three additional gates that cleaved poorly with Ro: G{u5,v1}, G{u16,v1}, and G{t61,u2r} (Figures S9, S10).

To further expand the library of ribozyme sequences to choose from, we tested G{u1,w2r} with three additional ribozymes with HDV-like folds^35–37^. Two of these ribozyme sequences had the expected performance (Figure S11). We also found that Ro with an extended P2 helix recovered the cleavage activity of G{u11,w2r} (Figure S12).

*These results indicate alternative ribozyme sequences can recover the performance of gates that cleave poorly with Ro. We identified a set of ribozyme sequences to screen in entirely new gate sequence contexts (Figure 7B)*.

### The design approaches for building ctRSD circuits yields expected performance in diverse cascades

To investigate how well the sequence domains and design approaches identified in this study performed across many sequence contexts, we tested 37 new gates within multilayer cascades. We designed the gates using only the *u* and *v* toeholds, alternating those sequences between the input and output domain of gates across circuit layers. We also designed all new gates in the cascades with the R3 ribozyme sequence. We first tested a two-layer cascade (G{uX,v1} to G{v1,u2r}), where *X* corresponded to sequence domains *3* through *17* (Figure 5A). All 15 gates cleaved well (Figure 5B) with reporter kinetics within the expected range from simulations (Figure 5C). To explore different sequence contexts, we next designed two cascades using gates with different output branch migration domains (6 and 7) in the second layer (Figure 5D). The gates in the second layer were also designed with two base extensions (*e*) in their input domains. We tested each of these second layer gates with ten different input branch migration domains and found the expected performance for all 20 cascades (Figure 5E,F). We also observed the expected performance for three- and four-layer cascades with gates that alternated the *u* and *v* output toeholds across layers (Figure S13). *These results demonstrate that alternating toeholds within gates and using the R3 ribozyme sequence are reliable design approaches that can facilitate the successful construction of more sophisticated ctRSD circuits (Figure 7A)*.

**Figure 5:**
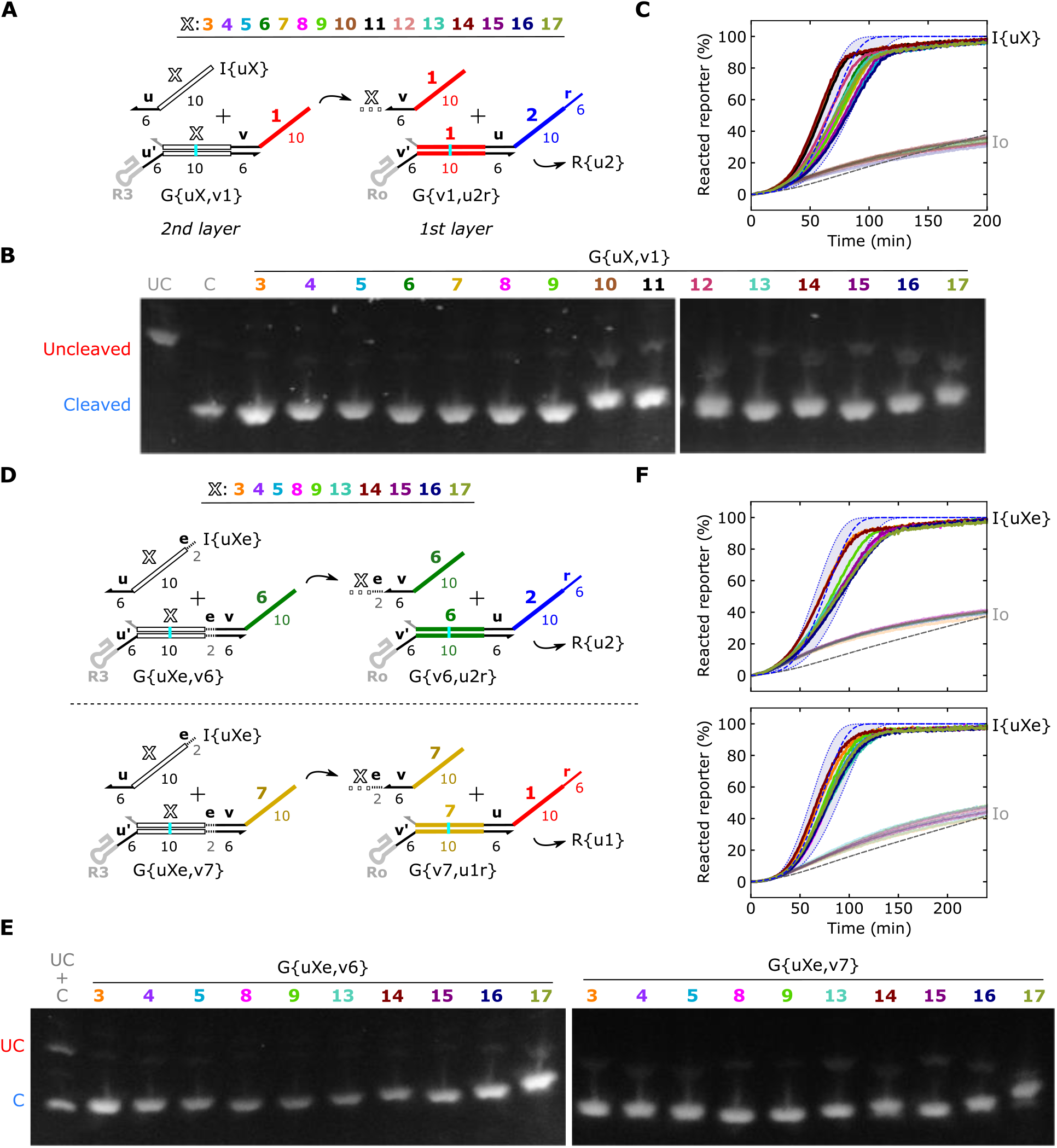
Characterization of two-layer ctRSD cascades. (**A,D**) Schematics of the two-layer cascades tested. The white *X* in the schematics denotes the domain of the gate that was varied. Gates in the 2^nd^ layer do not have the *r* domain because they do not connect to a reporter. (**B,E**) Denaturing gel electrophoresis results of the gates indicated above the gels. UC and C are size markers described in Figure 2. In (E), the UC and C templates were mixed at equal concentrations in a single well. (**C,F**) Reporter kinetics of the gates indicated above the plots. The line colors in the plots correspond to the domain number colors in the schematics in (A) and (D). The blue dashed lines represent simulation results with the shaded region spanning 5k_rsd_ to k_rsd_/5 for gates in the 2^nd^ layer of the cascades. Gate and input templates were present at 25 nmol/L. Reporters were present at 500 nmol/L. See Supporting Note 6 for individual kinetic plots.

### Short spacer sequences adjacent to the input toehold increase the rate of output production

Even though the cascades performed as expected, it could be desirable to tune the kinetics of individual gates in a circuit without redesigning upstream components, *i.e*., without changing the length or sequence of the input toehold on a gate and upstream inputs or outputs. Previously, we found adding a single-stranded region, termed a spacer (*s*), between the ribozyme and the input toehold on the gates (Supporting Note 3) increased the rate of output production. Compared to gates without a spacer, reporter kinetics for gates with two-base or four-base spacers were consistent with ~10-fold and ~100-fold higher RNA strand displacement rate constants, respectively^20^. In agreement with these results, we further found inserting a four-base spacer increased the putative RNA strand displacement rate constant ~100-fold for gates with either a *v* or *u* input toehold (Figures S14, S15). *These results present spacers adjacent to the input toehold of a gate as a design approach to increase output kinetics without changing upstream circuit components (Figure 7B)*.

### ctRSD gates perform well with changes in transcriptional encoding

Other than evaluating different ribozyme sequences, changes in transcriptional encoding of ctRSD gates have not been investigated (Figure 1C). All gates were transcribed in same orientation, starting at the 5′ end of the output strand (O → G′, Figure 6A). Further, the 5′ hairpin, 3’ terminator, and 3’ linker (*L*) sequences were the same for all gates. We next investigated whether the ctRSD gate design tolerated changes in transcriptional encoding without compromising performance.

**Figure 6:**
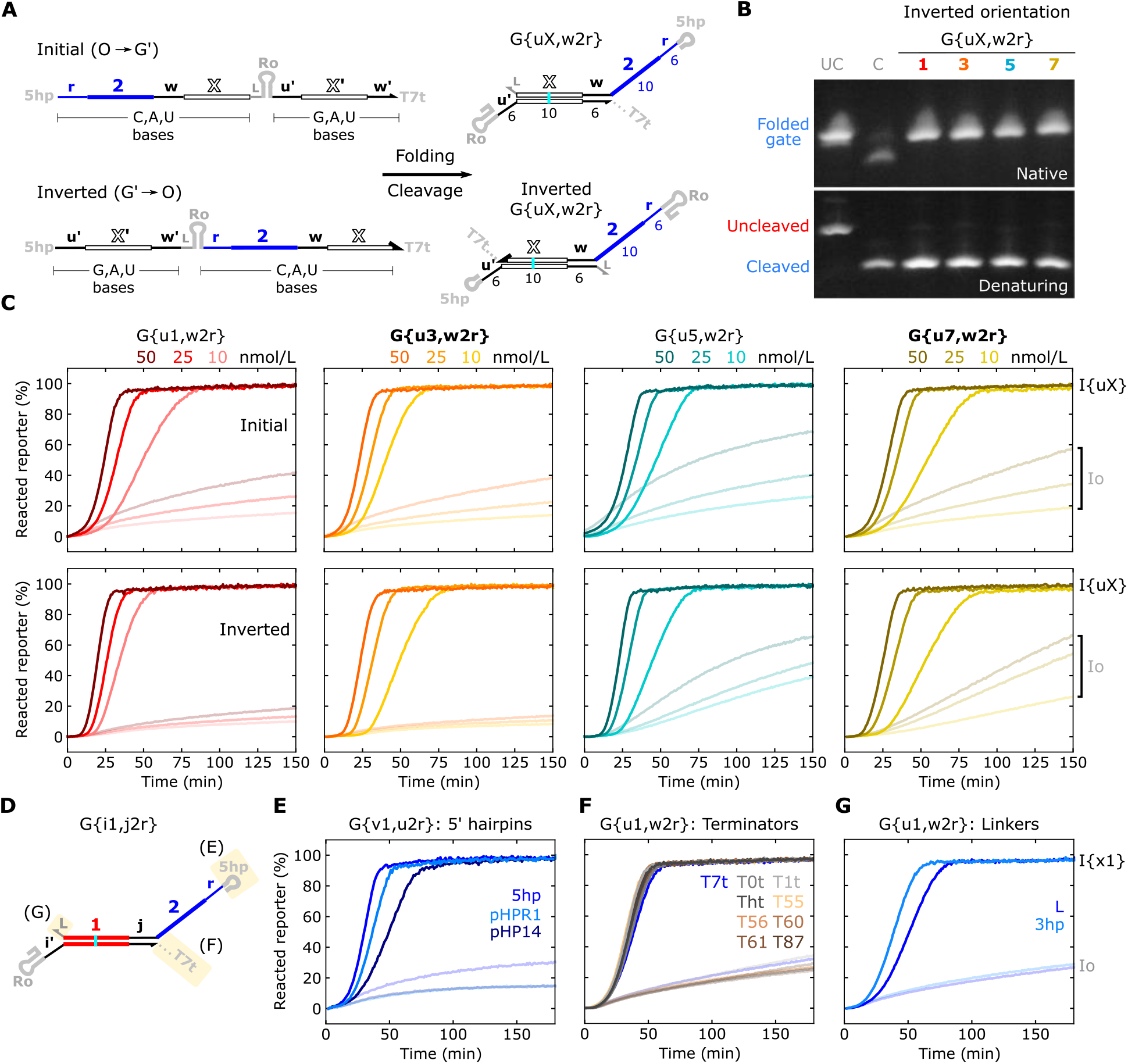
Characterization of ctRSD gates with changes in transcriptional encoding. (**A**) ctRSD gate transcription orientations. The initial design starts transcription at the 5′ end of the output (O) strand and the inverted design starts at the 5′ end of the input toehold domain on the G′ strand. The 5′ hairpin (5hp) and terminator (T7t) domains, which are present on all gates, are shown here for comparison. (**B**) Gel electrophoresis results for gates transcribed in the inverted orientation. UC is an uncleaved gate size marker, which is G{u1,w2r} with a single base mutation in Ro that prevents cleavage. C is a cleaved gate size marker, which is the O strand of G{u1,w2r} transcribed in the inverted orientation, the largest product of a cleaved gate with inverted transcription orientation. (**C**) Reporter kinetics for gates with initial (top panel) or inverted (bottom panel) transcription orientations. Bolded gate names above plots indicate gates with G′ strands predicted to have secondary structure (Figure S17). Full color lines indicate gates transcribed with their designed input (I{uX}), and semi-transparent lines indicate gates transcribed with a scrambled input (Io). The three gate concentrations above the plots were tested. I{uX} templates were present at 50 nmol/L. Io was added in appropriate amounts to maintain 100 nmol/L total template concentration across samples. (**D**) A ctRSD gate highlighting the transcriptional encoding domains changed in subsequent panels. *i*, *j* represent toeholds *u, v*, or *w*. (**E**,**F**,**G**) Reporter kinetics for the gates above the plots. The legends in the plots denote different sequences for the transcriptional encoding domain specified above the plots. Gate and input templates were present at 25 nmol/L. Reporters were present at 500 nmol/L. See Figures S18 to S20 for gel electrophoresis results.

We first explored transcription orientation, because inverting the order in which gate strands are synthesized (G′ → O, Figure 6A) may reduce leak, *i.e*., the undesired production of output from a gate in the absence of the gate’s designed input. As described previously^20^, the leak mechanism is consistent with either truncated or misfolded gate transcripts that have an exposed output domain that reacts downstream (Figure S16). For gates transcribed in the initial orientation (O → G’, Figure 6A) many truncated transcripts would have a completely exposed output domain (Figure S16). If truncated transcripts were the primary source of leak, we reasoned that inverting the transcription orientation (G′ → O, Figure 6A) could reduce leak by shifting synthesis of the output domain to the 3′ end of the transcript (Figure S16). A potential drawback of the inverted transcription orientation is that undesired secondary structure in the G′ strand could be introduced by G-U wobble base pairing prior to synthesis of the output strand, compromising performance^20^.

Based on these considerations, we selected four gates to test using the inverted transcription orientation, including two with (G{u3,w2r}, G{u7,w2r}) and two without (G{u1,w2r}, G{u5,w2r}) G′ strands predicted to form secondary structure (Figure S17). All four gates with inverted transcription orientation cleaved well and produced the expected band in native gel electrophoresis experiments (Figure 6B), providing evidence these gate transcripts folded correctly. Gates in both transcription orientations had similar reporter kinetics when transcribed with the designed inputs. G{u1,w2r} and G{u3,w2r} transcribed in the inverted orientation had less leak than when transcribed in the initial orientation, consistent with a leak mechanism based on early truncation (Figure S16A). However, G{u5,w2r} and G{u7,w2r} had similar leak in both transcription orientations (Figure 6C), which may be more consistent with a leak mechanism based on misfolded gates. For example, in both transcription orientations most of the transcript is produced but the G′ strand misfolds to expose the output domain (Figure S16B). There was no obvious pattern between leak and gates with predicted secondary structure in the G′ strand. Further testing is required to fully understand the leak mechanisms and validate the performance of gates transcribed in the inverted orientation across more sequence contexts, especially for gates with input domains longer than the initial gate design.

To further explore the changes in transcriptional encoding of ctRSD gates, we tested gates with different 5′ hairpin (*5hp*), terminator (*T7t*), and linker (*L*) sequences (Figure 6D). We found a single ctRSD reaction performed well when the 5′ hairpin sequence of both the gate and the input were changed to either of two sequences reported previously to increase RNA stability^25,26^ (Figures 6E, S18). We next evaluated the performance of gates with one of eight terminator sequences. In addition to measuring reporter kinetics, we also used gel electrophoresis to measure the fraction of gate transcripts that terminated for each terminator, *i.e*., termination efficiency. The primary terminator used in this study (*T7t*) had the highest termination efficiency at ~75% in our assay conditions, while the next best terminators were ~50% efficient (Figure S19). Despite differences in termination efficiency, gates with different terminator sequences had similar reporter kinetics (Figure 6F, S19), suggesting the sequence of the terminator and the sequence downstream of the terminator did not influence performance in this context. Lastly, we designed a gate with a short, non-terminating hairpin (*3hp*) inserted in the linker (*L*) domain between the 3′ end of the output strand and ribozyme. A hairpin at this location could stabilize the output strand in environments with RNases^24^. This gate cleaved well and had similar reporter kinetics as the same gate without the hairpin linker (Figures 6G, S20).

*These results, together with the results using different ribozymes sequences, indicate ctRSD gates are amenable to changes in transcriptional encoding. Transcriptional encoding alterations offer opportunities to tune ctRSD gate performance, for example in applications with RNases present*. For example, 5′ and 3′ hairpins alter RNA stability in a sequence-specific manner^24–26,38^. In addition to the potential to reduce leak for some sequences, gates with the inverted transcription orientation produce outputs with stabilizing secondary structure at both their 5′ and 3′ ends in the ribozyme and terminator domains^24,38^, respectively.

## Conclusion

Here, we expanded the toolkit for building ctRSD circuits, compiling a diverse set of sequence domains for assembling circuit components with expected performance. Compared to previous reports^20^, the toolkit contains four-fold more input sequences, each of which was demonstrated in a broad range of sequence contexts. We developed and extensively tested a set of recommended design approaches that yield gates with expected performance across diverse sequence contexts (Figure 7A). Even if a specific gate has poor performance with the recommended design approaches, performance can likely be recovered with minor design alterations (Figure 7B). A key feature of ctRSD circuits is the potential for continuous component production and computation in complex environments, such as cell lysates or living cells. Our expanded toolkit also introduces sequence domains to explore and tune performance for applications in these settings (Figure 7C).

**Figure 7:**
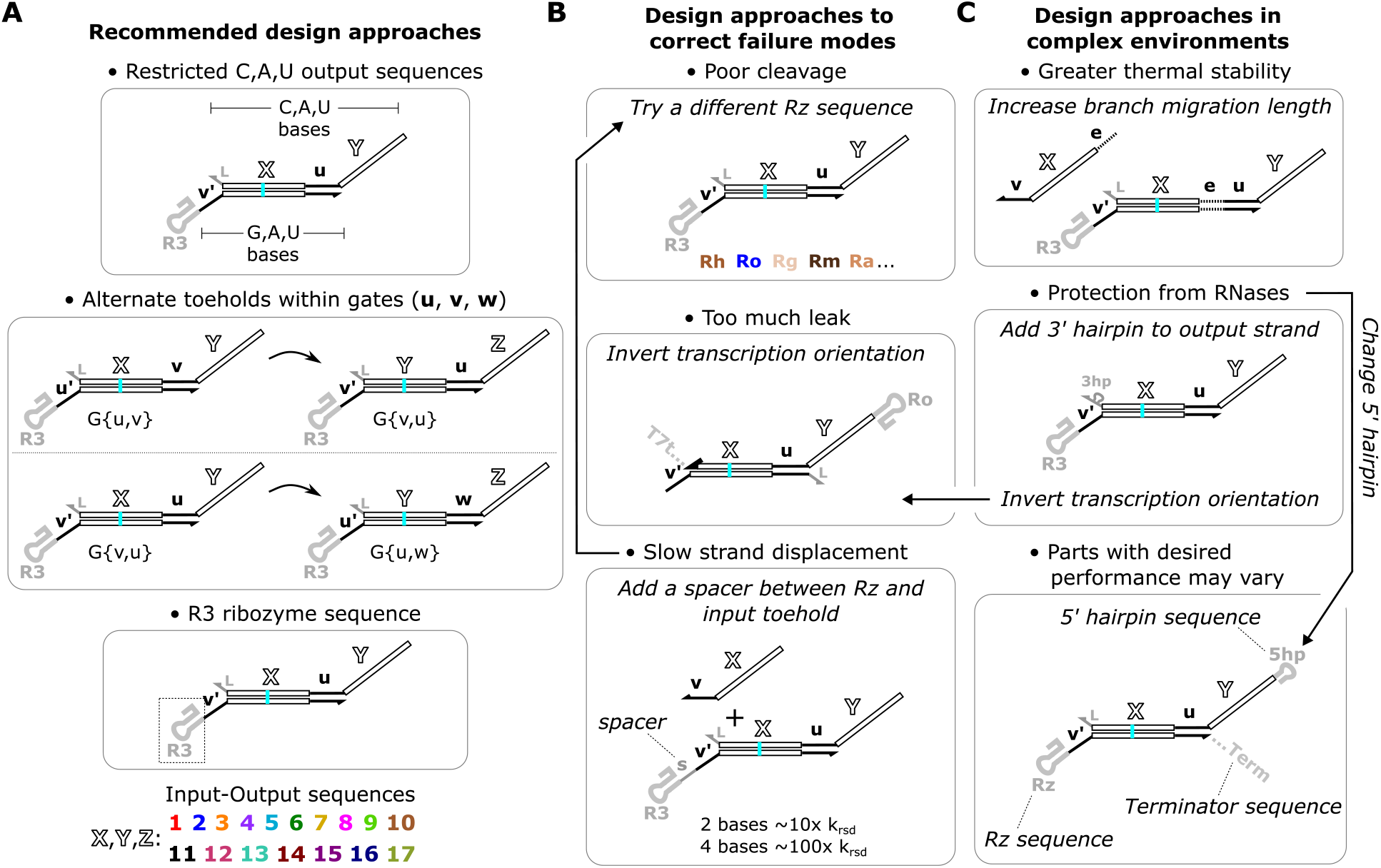
Summary of the design approaches for building ctRSD circuits. (**A**) Recommended design approaches to start with when building ctRSD circuits. (**B**) Design approaches to recover the performance of gates based on failure mode. If a gate has poor cleavage, try changing its ribozyme sequence in the order prescribed: Rh, Ro, Rg, Rm, Ra. If a specific gate has too much downstream leak, try inverting its transcription orientation. This change reduced leak in two out of four gates tested. If a gate has slow strand displacement kinetics, try adding a single stranded spacer sequence between the ribozyme and the input toehold. A two-base and four-base spacer can increase the putative RNA strand displacement rate constant nearly one to two orders of magnitude, respectively^20^. (**C**) Design approaches that could tune gate performance in complex environments, such as cell lysates or inside living cells. Top: The input branch migration domain can be extended to increase thermal stability of the gate complex. Middle: A hairpin can be added to the 3′ end of the output strand for protection from RNase degradation. Inverting gate transcription orientation could also protect the output from RNase degradation by introducing secondary structure at both ends of the output strand. Bottom: The optimal 5′ hairpin, ribozyme, and terminator sequences may differ across environments.

The toolkit, along with the design approaches we developed to further expand the toolkit, should now enable the construction of ctRSD circuits capable of much more sophisticated tasks^10,18,19^ than demonstrated so far^20^. To facilitate the design of such circuits, we developed a Python package for simulating the kinetics of ctRSD circuits and compiling component sequences^39^. This package enables rapid *in silico* prototyping of circuits with different architectures, initial conditions, and rate constants. The sequence compiler allows any combination of toolkit sequence domains to be stitched together.

While our design approaches typically yield gates with the expected performance, ctRSD components are susceptible to misfolding during transcription, making it difficult to predict *a priori* if a particular gate sequence will perform as expected. The measurements in this study provide insight into the salient features of the misfolded structures adopted by ctRSD gates (Figure S1, Supporting Note 5), but more detailed measurements are required to fully understand the failure mechanisms that lead to poor performance. For example, RNA sequencing techniques could be used to understand the secondary structures of misfolded gates^40–42^ and leak mechanisms^43,44^. These measurements would guide further improvements to ctRSD gate design and could enable predictive models of the ctRSD gate sequence-structure-function relationship that optimize gate sequences *in silico* for a given circuit.

Our results also highlight the importance of measuring both RNA structure and function to properly characterize ctRSD circuits, and this is likely applicable for other RNA-based circuits^11,12,23,45,46^. For example, a few gates we tested had poor ribozyme cleavage (incorrect structure) yet had reporter kinetics within the range expected from simulations (correct function) (Figure 2), suggesting reactions other than the designed mechanisms can unexpectedly produce the anticipated outputs (Figure S1C). As ctRSD circuits are moved to more complex environments, such as cells, it will be essential to adopt or develop measurements of both RNA structure and circuit function^47–50^ to ensure ctRSD reactions proceed *via* the designed mechanisms. The expanded toolkit, design approaches, and software tools developed here lay the foundation to begin exploring a wide range of applications for ctRSD circuits across environments.

## MATERIALS AND METHODS

### DNA and materials

DNA transcription templates were ordered in 96-well PCR plates as eBlock gene fragments eluted to 10 ng/μL in Buffer IDTE, pH 8.0 from Integrated DNA Technologies (IDT). eBlocks were amplified *via* polymerase chain reaction (PCR) with Phusion High-Fidelity PCR Master Mix (cat. no. F531L) from ThermoFisher Scientific and purified using Qiagen PCR clean-up kits (cat. no. 28104). To meet the length requirements for ordering eBlock DNA, flanking sequences were appended adjacent to the sequence of the ctRSD component amplified with PCR. Further, a G-U wobble base pair was introduced in the branch migration domain of the gates to reduce synthesis errors (see Supporting Note 1). DNA primers for PCR were ordered from IDT without purification (standard desalting). The fluorophore-modified strands of the DNA reporters were ordered from IDT without purification (standard desalting), and the quencher modified strands of the DNA reporters were ordered from IDT with HPLC purification. DNA reporter complexes were prepared by annealing 20 μmol/L of each strand in transcription buffer (heat to 90 °C for 5 min and then cool to 20 °C at 1 °C per min). For *in vitro* transcription experiments, T7 RNA polymerase (RNAP) (200 U/μL) and ribonucleotide triphosphates (NTPs) were ordered from ThermoFisher Scientific (cat. nos. EP0113 and R0481, respectively). 5x transcription buffer was prepared in house (200 mmol/L tris-HCl (pH 7.9), 30 mmol/L MgCl2, 50 mmol/L dithiothreitol, 50 mmol/L NaCl, and 10 mmol/L spermidine). Deoxyribonuclease (DNase) I (cat. no. M0303S) was purchased from New England Biolabs. Four percent agarose EX E-gels were purchased from Invitrogen (cat. no. G401004). All other chemicals were purchased from Sigma-Aldrich.

All DNA sequences from this study are available in Supporting File S1. Additionally, the ctRSD simulator 2.0 package (https://ctrsd-simulator.readthedocs.io/en/latest/SeqCompiler.html) contains a sequence compiling function to stitch together full gate sequences for any combination of sequence domains explored in this study.

### Transcription template preparation

Transcription templates were prepared by PCR of eBlock DNA (0.2 ng/μL) with Phusion High-Fidelity PCR Master Mix and forward and reverse primers (0.5 μM). PCR was conducted for 30 cycles with a 30 s denaturing step at 98 °C, a 30 s primer annealing step at 60 °C, and a 30 s extension step at 72 °C. A 3 min extension step at 72 °C was executed at the end of the program. Following PCR amplification, the samples were purified with a Qiagen PCR clean-up kit (cat. no. 28104) and eluted in Qiagen Buffer EB (10 mmol/L tris-HCl, pH 8.5). After PCR clean-up, DNA concentrations were measured with A260 on a DeNovix D-11 Series Spectrophotometer. We note that using (10 to 25) fold less eBlock DNA (0.02 ng/μL to 0.008 ng/μL) for PCR produced similar yields with reduced side products compared to 0.2 ng/μL (see Supporting Note 1).

### Gel electrophoresis and imaging

4 % agarose EX E-gels were used for all RNA gel electrophoresis experiments. These gels are prestained with SYBR Gold for fluorescence imaging. Electrophoresis was conducted on an E-gel powerbase, and all E-gels were imaged using the E-gel power snap camera (Invitrogen, cat. no. G8200). To prepare RNA for gel electrophoresis, 25 nmol/L of DNA templates were transcribed at 37 °C for 30 min in transcription conditions (see the next section) with 1 U/μL of T7 RNAP. Transcription was stopped by adding both CaCl2 to a final concentration of 4.17 mmol/L and DNase I to a final concentration of 0.17 U/μL to degrade DNA templates. After addition of the DNase I, the samples were left at 37 °C for 30 min and subsequently characterized with gel electrophoresis. For denaturing gels, a solution of 100% formamide and 36 mmol/L EDTA was mixed 1:1 by volume with the samples before electrophoresis, and the samples were heated to 85 °C for 5 min. The samples were then immediately loaded on gels for electrophoresis and run for (20 to 30) min before imaging. For native gels, E-gels that had been cooled to 4 °C were sandwiched between frozen cold packs to keep the gels below 37 °C during electrophoresis and were run for (45 to 60) min before imaging. Gel images were not postprocessed; any brightness and contrast adjustments were applied uniformly across each image during acquisition to aid visualization and facilitate qualitative comparison between different gel images. Unless otherwise stated in the figure captions, white spaces between gel images represent images taken from different gels. Each gel had its own size markers, the uncleaved (UC) and cleaved (C) controls, and these controls were used to align gel images for comparison.

### Fluorescent DNA reporter kinetic assays

The *in vitro* transcription reactions with DNA reporter complexes were conducted at 37 °C in a BioTek Synergy Neo2 plate reader (see “Fluorescence data acquisition and normalization”). The reactions took place in transcription buffer prepared in house (see “DNA and materials”) supplemented with 2 mmol/L final concentration of each NTP type (adenosine triphosphate, uridine triphosphate, cytidine triphosphate, and guanosine triphosphate). Reactions were typically conducted in 70 μL volumes in Greiner μClear Black 96-well plates (cat. no. 655096) read from the bottom. Gate and input templates were added individually to each well. A master mix containing water, transcription buffer, NTPs, DNA reporter, and T7 RNAP (added last) was then prepared and pipette into each well containing the DNA templates. Unless otherwise stated, 500 nmol/L DNA reporter was used. As described previously^20^, to compare the response of a given ctRSD circuit to different input template concentrations or a different number of input templates, the same total template concentration was used across samples in a given experiment to ensure the same transcriptional load. A template (Io) that produces a scrambled input, *i.e*., an RNA the length of an input that does not interact with the gates, was added to maintain a constant total template concentration across samples. See Supporting File S2 for transcription template and T7 RNAP concentrations for individual experiments.

### Fluorescence data acquisition and normalization

Fluorescence measurements were conducted in BioTek Synergy Neo2 plate readers. Reactions were conducted in 70 μL volumes in Greiner μClear Black 96-well plates (cat. no. 655096) read from the bottom. The DNA reporter complexes were fluorescently labeled as follows: R{w2} and R{u2} were modified with a HEX dye, which was measured with excitation: 524 nm (20 nm bandwidth), emission: 565 nm (20 nm bandwidth), and a gain of 80 to 100 to ensure the fluorescence values were within the linear range of detection. R{w1} and R{u1} were modified with a FAM dye, which was measured with excitation: 487 nm (20 nm bandwidth), emission: 528 nm (20 nm bandwidth), and a gain of 60 to 80 to ensure the fluorescence values were within the linear range of detection. Fluorescence readings were taken every 46 s. Fluorescence data was then normalized as:

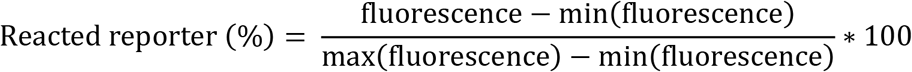

The initial measurements in each well at the beginning of an experiment served as the minimum fluorescence value for normalization. A control well in which the ctRSD reaction had resulted in 100% reacted reporter served as a maximum value for normalization.

### Transcription rate calibration and sample variability

As described previously^20^, the transcription rate in the experiments presented here depends on the concentration of T7 RNAP and the total concentration of DNA templates. Variability of T7 RNAP activity across manufacturer lots is an expected source of variation^51^. So, in each of our experiments, we calibrated the transcription rate for the T7 RNAP stock and concentration and total template concentration for that given experiment. This sample was used to calibrate the first order transcription rate constant used to simulate the experiment, thus accounting for transcription rate differences when assessing the agreement between experimental results and model predictions. In this study, the sample used to calibrate transcription rate was typically a gate and input pair with previously well-characterized kinetics, for example, G{u1,w2r} and I{u1} (see Supporting Note 6).

Given that a primary goal of this study was to assess how different ctRSD gate sequence combinations influenced gate performance, we confirmed that measured differences in performance across gate sequences were reproducible and distinguishable from experimental noise (Supporting Note 4). Briefly, we selected two to three gates for both one- and two-layer cascades and performed the DNA reporter assay in triplicate for each gate. These experiments recapitulated the relative DNA reporter kinetics initially measured for these gates. The technical replicates exhibited a standard deviation of < 6 % from the mean. We also prepared samples from a second PCR for each of the selected gates and repeated the above experiments with similar results (Supporting Note 4). For the small circuits studied here, we do not expect this level of variability to influence our conclusions. Unless otherwise stated, DNA reporter experiments were conducted with a single experimental replicate.

### Kinetic simulations

The simulations in this study are based on a deterministic model composed of systems of coupled ordinary differential equations derived from mass action kinetics. Systems of ordinary differential equations were integrated numerically using the *solve_ivp()* function in Python’s SciPy package (version 1.9.2, Python 3.8.3) using the Spyder integrated development environment (version 4.1.4). The specific chemical reactions and rate constants relevant to this study are described in detail in Supporting Note 2. See Supporting File S2 and figure captions for additional details on specific simulations.

To facilitate this and future studies, a user-friendly ctRSD kinetic simulation package, implemented in Python, was developed for conducting ctRSD simulations: https://ctrsd-simulator.readthedocs.io/en/latest/index.html. This software allows users to input the ctRSD components and concentrations that describe a system they want to simulate. The software automatically assembles the set of differential equations describing the system and simulates the system based on the provided initial conditions. Beyond the simple cascades described in this study, the software contains many additional ctRSD chemical reactions and rate constants are customizable for individual species.

## Supporting information

Simulation Scripts

## Acknowledgments

The authors thank S. Baral, T. Gorochowski, S. Servetas, and M. Zwolak for insightful comments on the manuscript and N. Alperovich, E. Romantseva, D. Ross, G. Taghon, O. Vasilyeva for helpful discussions throughout this work.

## Author contributions

SWS conceived, designed, and performed most of the research. MEW conducted experiments in Supporting Note 4. SWS conducted data analysis and executed simulations. SWS and TMM developed the simulation software. EAS supervised the research. SWS, EAS, and MEW wrote the manuscript.

## Data and materials availability

All data needed to evaluate the conclusions in the paper are present in the paper and/or the Supporting Information. Fluorescence data and simulation code are available as Supporting Files. The latest versions of the simulation code will be maintained on GitHub at https://github.com/usnistgov/ctRSD-simulator and the full documentation of the simulation software package is available at https://ctrsd-simulator.readthedocs.io/en/latest/.

## Disclaimer

Certain commercial entities, equipment, or materials may be identified in this document to describe an experimental procedure or concept adequately. Such identification is not intended to imply recommendation or endorsement by the National Institute of Standards and Technology, nor is it intended to imply that the entities, materials, or equipment are necessarily the best available for the purpose. Official contribution of the National Institute of Standards and Technology; not subject to copyright in the United States.

