## Supporting Information for "Design approaches to expand the toolkit for building cotranscriptionally encoded RNA strand displacement circuits"

**Description of Supporting Information Files**

`Supporting File S1.xlsx`: This file contains the eBlock DNA sequences for the ctRSD components tested in this study.

> The column 'seq_compile' specifies the custom parameters of the `ctRSD_seq_compiler()` function needed to generate the sequence.
> Sequence compiler documentation:
> https://ctrsd-simulator.readthedocs.io/en/latest/SeqCompiler.html
>
> An Excel file containing the sequences of individual ctRSD domains can be found at:
> https://github.com/usnistgov/ctRSD-simulator/blob/main/ctRSD-simulator-2.0/Sequence Compiler/ctRSD_domains_list.xls

`Supporting File S2.xlsx`: This file contains the concentrations of components used in each DNA reporter assay in this study.

> This includes the concentrations of DNA templates, DNA reporters, T7 RNAP, and the parameters used to simulate the results of the experiments (transcription and effective RNA strand displacement rate constants).

`Supporting File S3.xlsx`: This file contains the raw and normalized fluorescence data for each DNA reporter assay in this study.

### Figures S1 – S20

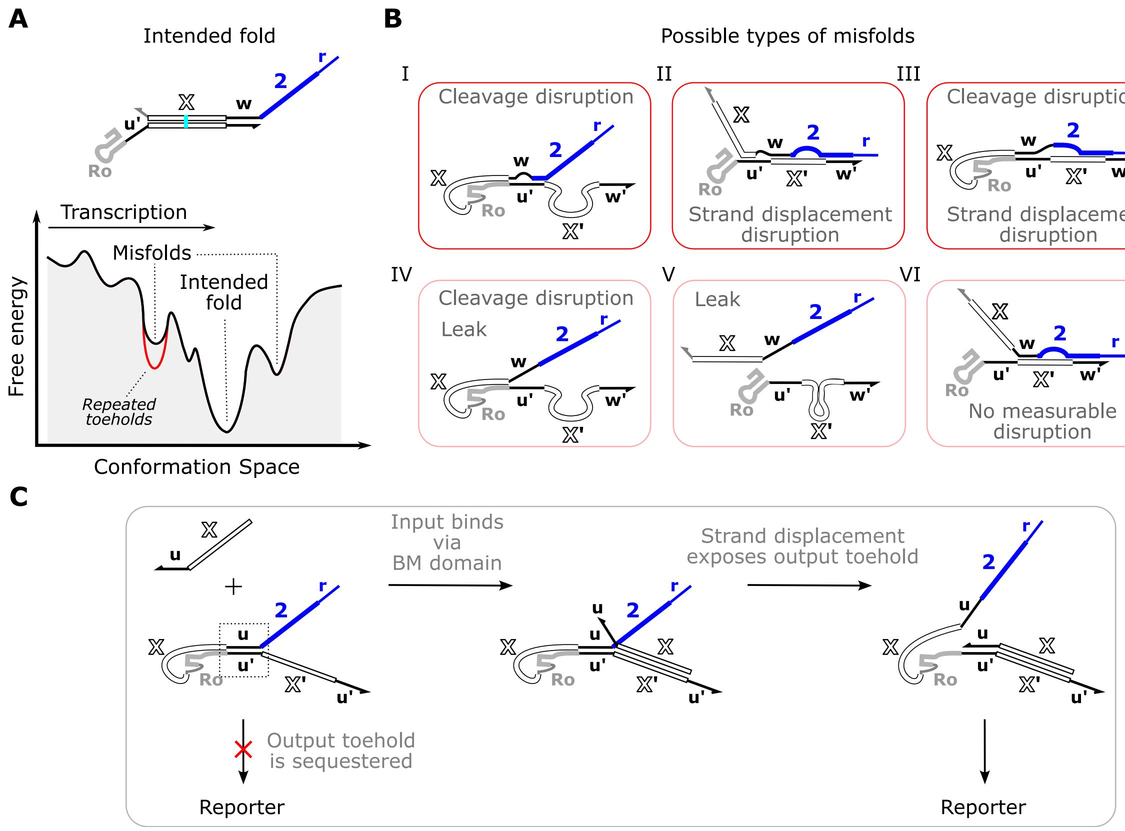

**Figure S1:** Possible ctRSD gate misfolding pathways and the potential impact of these misfolds on performance measurements. Gates with poor performance likely have a large population of transcripts that have adopted undesired misfolded structures. Our measurements of ribozyme cleavage, reporter kinetics with the correct input, and leak output production provide insight into the salient features of these misfolded structures. (**A**) Schematic of a hypothetical ctRSD gate energy landscape during transcription. Because gates fold out-of-equilibrium during transcription[1,2], it is possible that partially transcribed gates fold into local free energy minimum structures that are slow to rearrange after the full length transcript is produced. Presumably, repeating toehold sequences within gates can stabilize misfolds, because there are two binding sites for the output toehold, as indicated by the deep free energy well in red. Alternatively, misfolded gates could represent structures expected in the ensemble for a given sequence at equilibrium. (**B**) Possible misfolds of ctRSD gates and the inferred influence these misfolds could have on performance measurements. The first three possible misfolds are consistent with experimental measurements of specific gates in this study. For example, G{u11,w1r}, G{u10,u2r}, G{u10,u1r}, and G{u6,u2r} cleaved poorly but had expected reporter kinetics, suggesting misfolds in which the ribozyme is disrupted but an input can still release an output with the expected kinetics (I). G{u3,u2r}, G{u5,u2r}, and G{u6,u1r} cleaved well but had reporter kinetics that were slower than expected, suggesting misfolds that do not disrupt the ribozyme but interfere with strand displacement (II). G{u11,w2r} cleaved poorly and had reporter kinetics that were slower than expected, suggesting misfolds that do not disrupt the ribozyme and strand displacement (III). Because no drastic increases in leak were observed for any of the gates tested in this study, the structures in IV and V do not appear to be prominent misfolds. The structure in VI is possible but would be difficult to distinguish from a correctly folded gate with the measurements used in this study, *i.e.*, this structure would cleave and undergo strand displacement with its input *via* the input toehold. (**C**) A possible reaction pathway that could explain how some gates have poor cleavage and low leak but relatively fast RNA strand displacement. It is possible that the strand displacement reaction could also induce the ribozyme to refold and cleave.

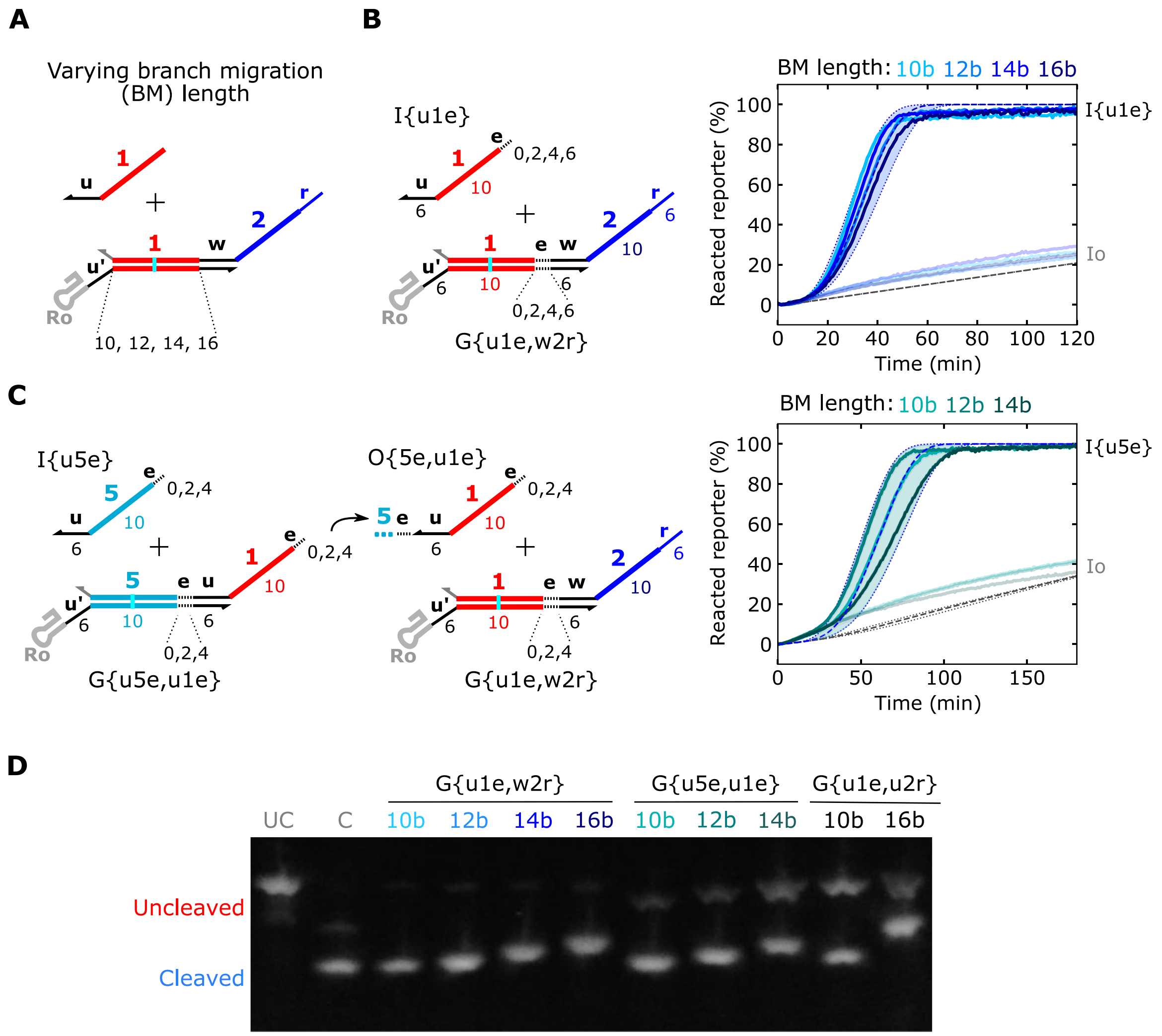

**Figure S2:** Extending input branch migration (BM) domains in one-layer and two-layer ctRSD cascades. (**A**) Schematic of the varying lengths of input branch migration lengths for a single gate. (**B**) Reporter kinetics of the gates shown to the left of the plot. The *e* domain was either (0, 2, 4, or 6) bases to produce input BM domains of (10, 12, 14, or 16) bases. Note both the input domain of the gate and the input to the gate are extended by the *e* domain. Gate and input templates were present at 25 nmol/L nmol/L. R{w2} was present at 500 nmol/L. (**C**) Reporter kinetics of the cascades shown to the left of the plot. Gate and input templates were present at 25 nmol/L and 50 nmol/L, respectively. R{w2} was present at 500 nmol/L. For (B) and (C), the dashed lines represent simulation results with the shaded region spanning $2k_{rsd}$ to $k_{rsd}/2$ for each ctRSD gate. (**D**) Denaturing gel electrophoresis results of the gates indicated above the gels. UC and C are size markers described in Figure 2 of the main text. See Methods for additional experimental concentrations, simulation parameters, and gel image processing.

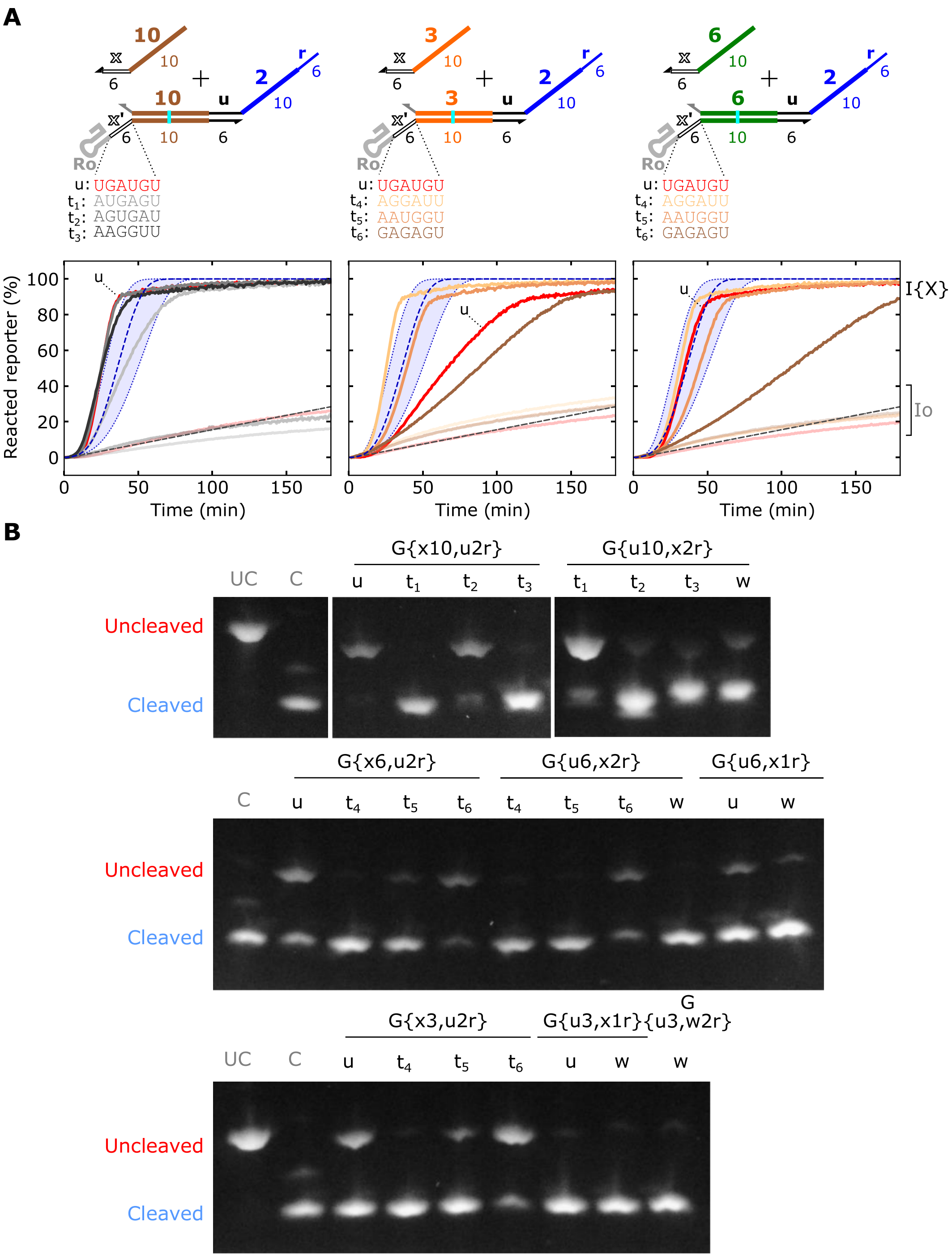

**Figure S3:** Varying input toehold sequences with input domains 10, 3, and 6. (**A**) Reporter kinetics of the gates indicated above the plots. The line colors in the plot correspond to the toehold sequence colors above the plot. Full color lines: designed input, semi-transparent lines: scrambled input (Io). The blue dashed lines represent simulation results with the shaded region spanning $4k_{rsd}$ to $k_{rsd}/4$. Gate and input templates were present at 25 nmol/L. R{u2} was present at 500 nmol/L. (**B**) Denaturing gel electrophoresis results of the gates indicated above the gels. UC and C are size markers described in Figure 2 of the main text. See Methods for additional experimental concentrations, simulation parameters, and gel image processing. Compared to the results for G{x1,u2r} and G{u1,x2r} in Figure 4 of the main text, toeholds $t_1$ and $t_2$ result in poor cleavage for gates with input domain *10* so these two toeholds were not considered.

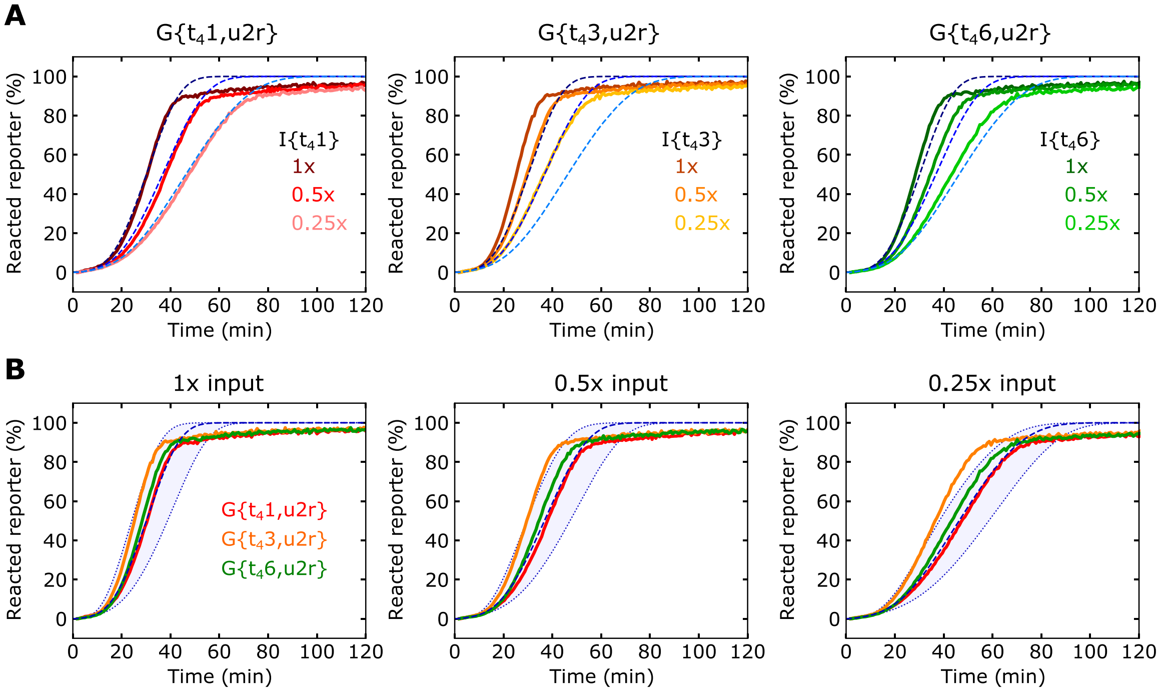

**Figure S4:** Comparison of $t_4$ ($v$) toehold sequence with different input domains. (**A**) Reporter kinetics of the gates indicated above the plots with different input template concentrations. Gate templates were present at 25 nmol/L and R{u2} was present at 500 nmol/L. Input templates were present at the concentrations specified in the plot legends with 1x referring to 25 nmol/L. (**B**) The same data from panel (A) replotted to compare gates in each plot instead of input concentrations. The blue dashed lines represent simulation results with the shaded region spanning $3k_{rsd}$ to $k_{rsd}/3$.

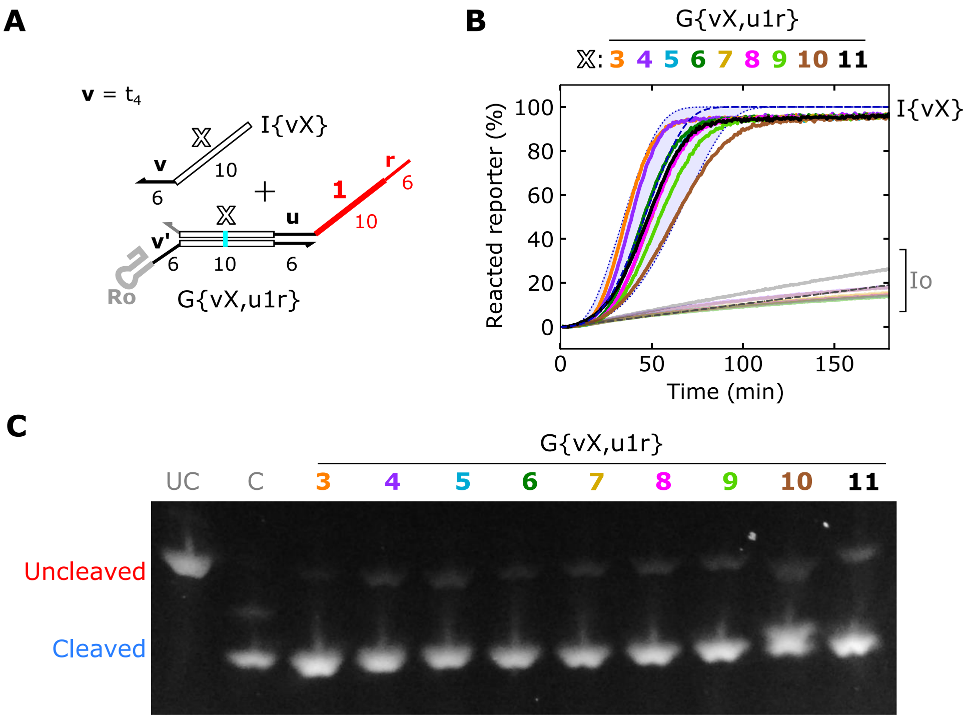

**Figure S5:** Characterization G{vX,u1r} with different input domains. (**A**) Schematic of G{vX,u1r}, where *X* denotes different input domain sequences. Note the *v* toehold is the same sequence as the $t_4$ toehold in Figure 3 of the main text. (**B**) Reporter kinetics of the gates indicated (A). The line colors in the plots correspond to the input domain number colors above the plot. The blue dashed lines represent simulation results with the shaded region spanning $4k_{rsd}$ to $k_{rsd}/4$. Gate and input templates were present at 25 nmol/L. R{u1} was present at 500 nmol/L. (**C**) Denaturing gel electrophoresis results of the gates indicated above the gels. UC and C are size markers described in Figure 2 of the main text. See Methods for additional experimental concentrations, simulation parameters, and gel image processing. See Supporting Note 6 for individual kinetic plots.

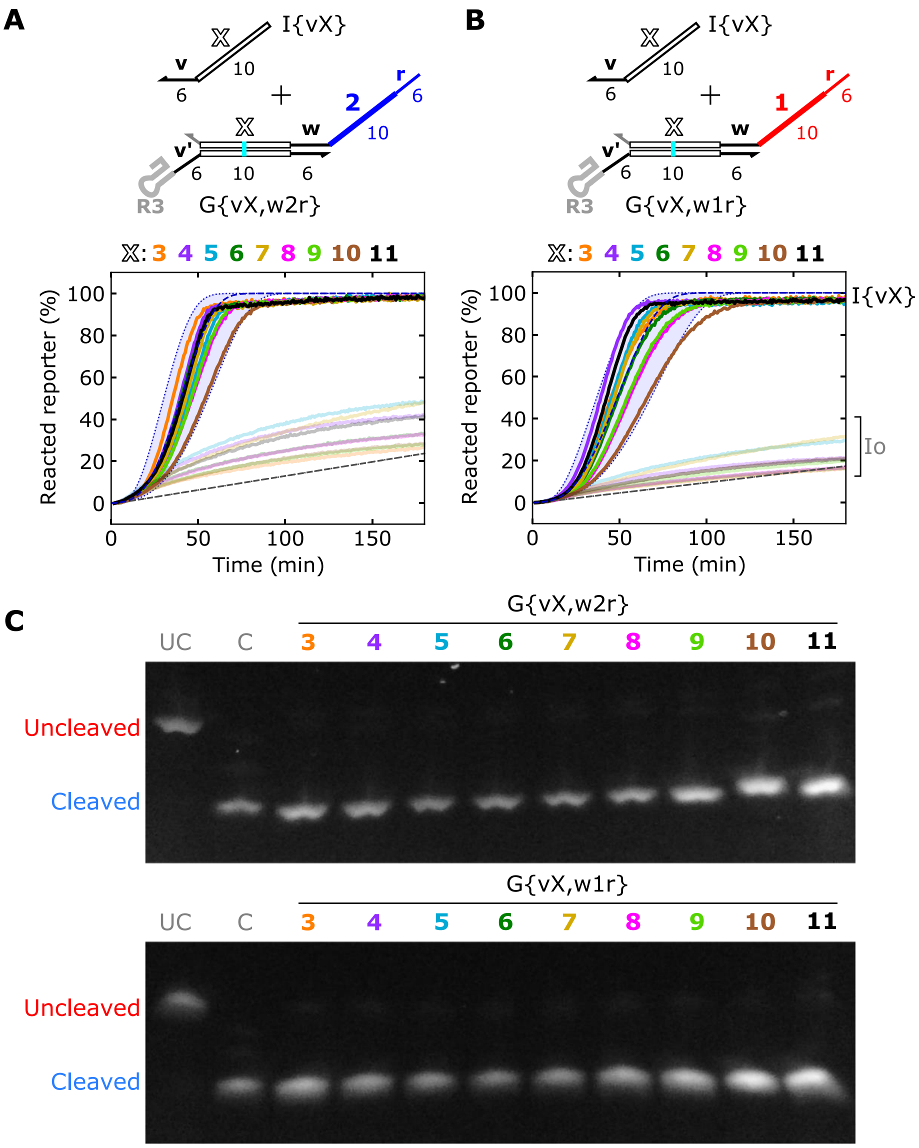

**Figure S6:** Characterization G{vX,w2r} and G{vX,w1r} with different input domains. (**A**,**B**) Reporter kinetics of the gates indicated above the plots. The line colors in the plot correspond to the domain colors above the plot. The blue dashed lines represent simulation results with the shaded region spanning $4k_{rsd}$ to $k_{rsd}/4$. Gate and input templates were present at 25 nmol/L. Reporters were present at 500 nmol/L. (**C**) Denaturing gel electrophoresis results of the gates indicated above the gels. UC and C are size markers described in Figure 2 of the main text See Methods for additional experimental concentrations, simulation parameters, and gel image processing. See Supporting Note 6 for individual kinetic plots.

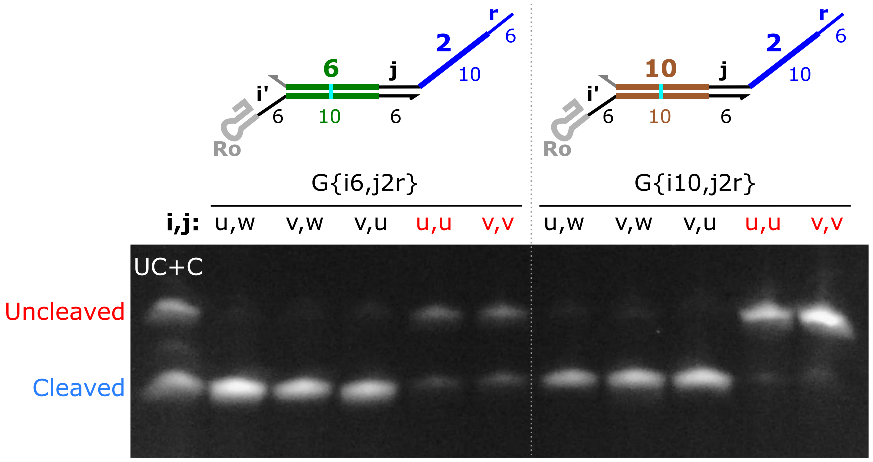

**Figure S7:** Characterization of gates with different combinations of input-output toeholds. Denaturing gel electrophoresis results of the gates indicated above the gels. UC and C are size markers described in Figure 2 of the main text. The UC and C templates were mixed at equal concentrations in a single well. Repeating either the *u* or *v* toeholds results in poor cleavage, while all combinations of non-repeated toeholds cleave well. See Methods for gel image processing.

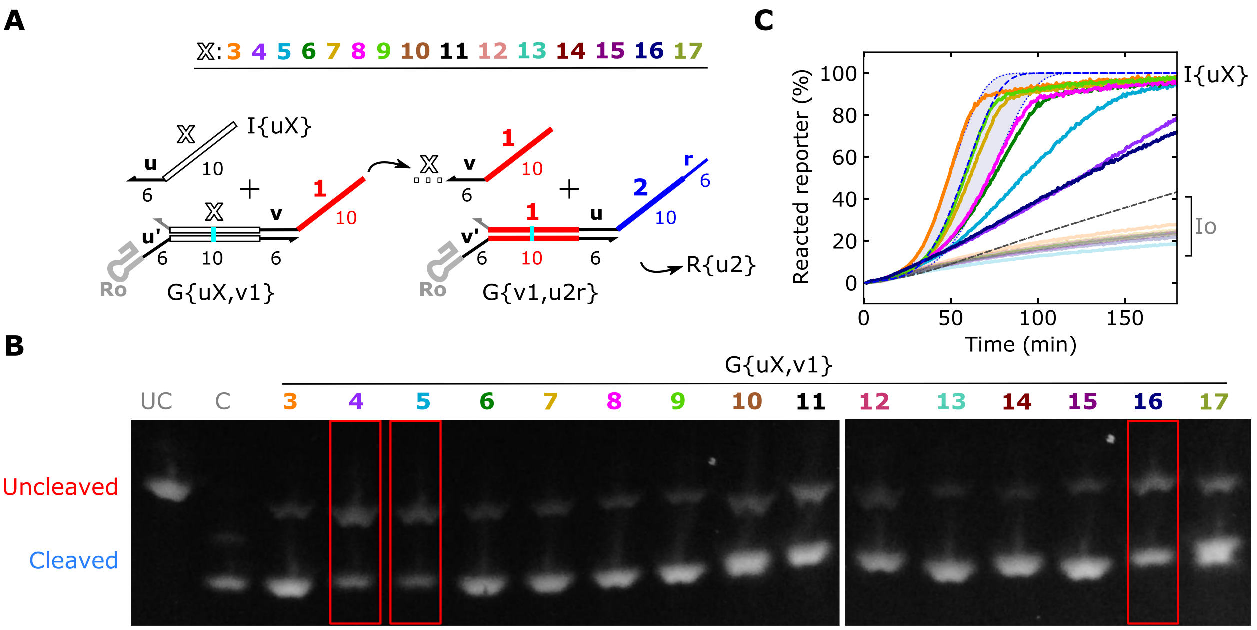

**Figure S8:** G{uX,v1} to G{v1,u2r} cascades with Ro for all gates. (**A**) Schematic of the two-layer cascades tested. The white *X* in the schematics denotes the domain of the gate that was varied. (**B**) Denaturing gel electrophoresis results of the gates indicated above the gels. UC and C are size markers described in Figure 2 of the main text. (**C**) Reporter kinetics of the gates indicated in (A). The line colors in the plots correspond to the domain number colors in the schematics in (A). Note only *X* = 3 through 9 and 16 were tested in this experiment. The blue dashed lines represent simulation results with the shaded region spanning $5k_{rsd}$ to $k_{rsd}/5$ for gates in the second layer of the cascades (G{uX,v1}). Gate and input templates were present at 25 nmol/L. R{u2} was present at 500 nmol/L. See Methods for additional experimental concentrations, simulation parameters, and gel image processing.

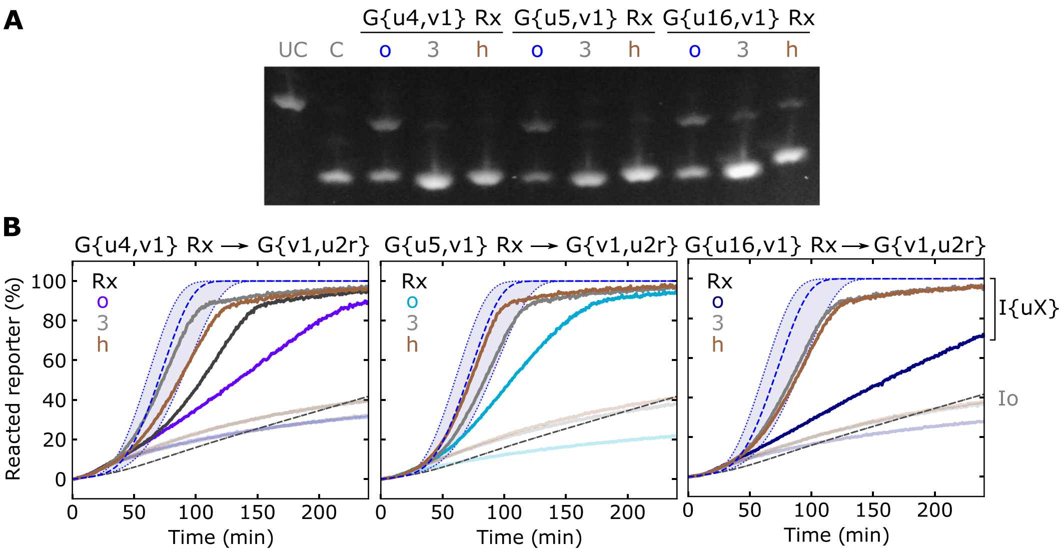

**Figure S9:** Characterization of G{v4,u1}, G{v5,u1}, and G{v16,u1} with different ribozymes. (**A**) Denaturing gel electrophoresis results of the gates indicated above the gels, where o, 3, and h refer to the ribozyme sequence (Rx). UC and C are size markers described in Figure 2 of the main text. (**B**) DNA reporter kinetics of the cascades indicated above the plots. The line colors correspond to the Rx colors that denote ribozyme sequence in the plots. The blue dashed lines represent simulation results with the shaded region spanning $5k_{rsd}$ to $k_{rsd}/5$ for gates in the second layer of the cascades. G{v1,u2r} was designed with the Ro ribozyme sequence in all experiments. Gate and input templates were present at 25 nmol/L. R{u2} was present at 500 nmol/L. The data in the leftmost plot of (B) is also presented in Figure 4 of the main text. See Methods for additional experimental concentrations, simulation parameters, and gel image processing.

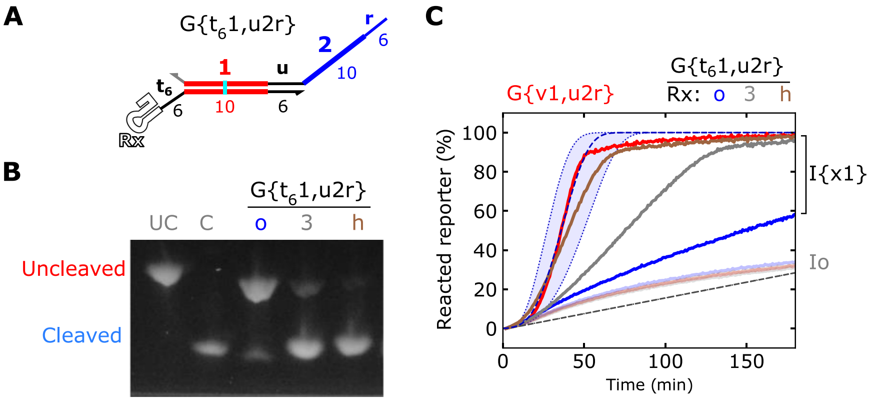

**Figure S10:** Characterization of G{$t_6$1,u2r} with different ribozymes. **A**) Schematic of G{$t_6$1,u2r}, where Rx denotes different ribozyme sequences, *i.e.*, Ro, R3, Rh. (**B**) Denaturing gel electrophoresis results of the gates indicated above the gels. UC and C are size markers described in Figure 2 of the main text. (**C**) Reporter kinetics of the gates indicated (A). The line colors in the plots correspond to the Rx colors that denote ribozyme sequence above the plots. G{v1,u2r} was designed with the Ro sequence. The blue dashed lines represent simulation results with the shaded region spanning $4k_{rsd}$ to $k_{rsd}/4$. Gate and input templates were present at 25 nmol/L. R{u2} was present at 500 nmol/L. See Methods for additional experimental concentrations, simulation parameters, and gel image processing.

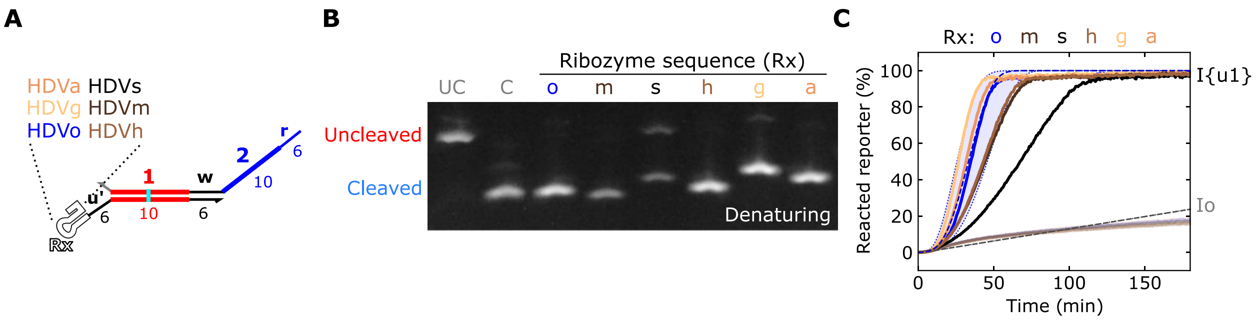

**Figure S11:** Characterization of G{u1,w2r} with different ribozymes. (**A**) Schematic of G{u1,w2r} with different ribozyme sequences. (**B**) Denaturing gel electrophoresis results of G{u1,w2r}with the ribozyme sequence indicated above the gel. UC and C are size markers described in Figure 2 of the main text. (**C**) Reporter kinetics for G{u1,w2r} with the ribozyme sequences indicated above the plot. Gate and input templates were present at 25 nmol/L and 50 nmol/L, respectively. R{w2} was present at 500 nmol/L. See Supporting Note 3 for sequence schematics of each ribozyme. See Methods for additional experimental concentrations, simulation parameters, and gel image processing.

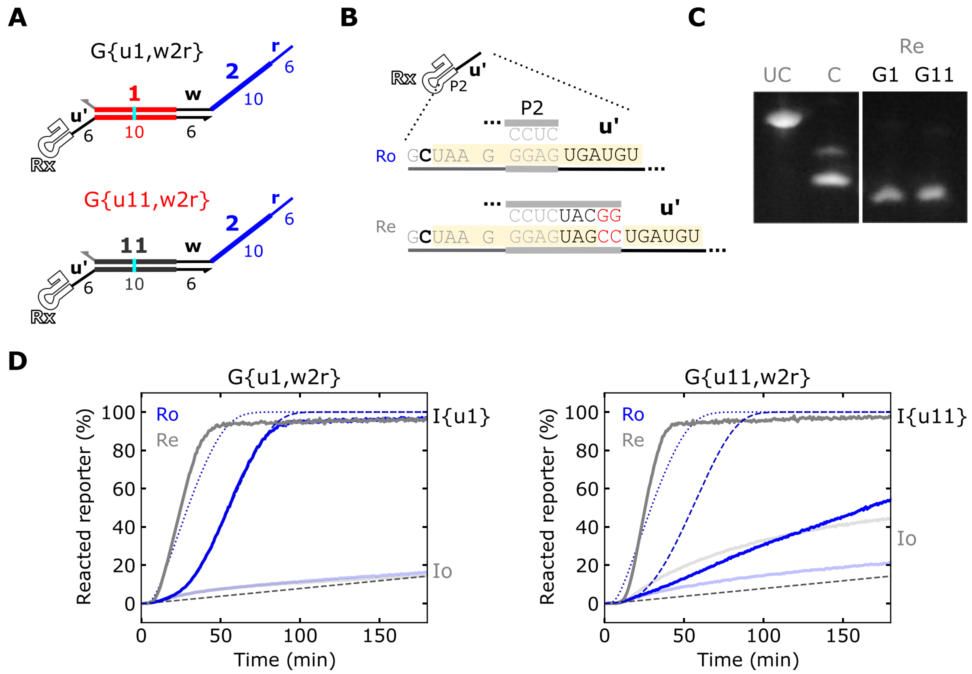

**Figure S12:** Characterization of gates with an extended P2 helix. (**A**) Schematics of gates tested. (**B**) Sequence schematics of Ro and the extended P2 helx (Re). (**C**) Denaturing gel electrophoresis results for G{u1,w2r} Re (G1) and G{u11,w2r} Re (G11). UC and C are size markers described in Figure 2 of the main text. The left and right image are from the same gel but from opposite sides. (**D**) Reporter kinetics for the gates indicated above the plots. The blue dashed lines are simulation with $k_{rsd} = 10^3$ L·mol$^{-1}$s$^{-1}$, and the gray dotted lines are simulations with $k_{rsd} = 10^5$ L·mol$^{-1}$s$^{-1}$. The extended P2 stem might act as a spacer (*s* domain in Supporting Note 3), which effectively increases strand displacement kinetics. Gate and input templates were present at 25 nmol/L. R{w2} was present at 500 nmol/L. See Methods for additional experimental concentrations, simulation parameters, and gel image processing.

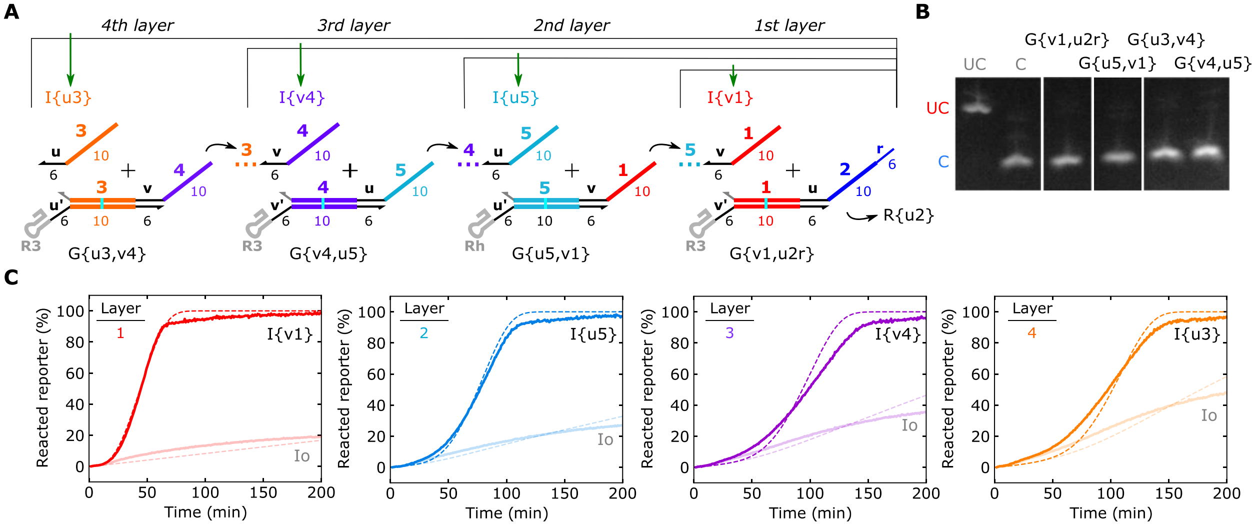

**Figure S13:** One- to four-layer cascades using gates with alternating *u*,*v* input-output toeholds. (**A**) Layout of one- to four-layer cascades. The Rh ribozyme was chosen for G{u5,v1} because this ribozyme was consistent with a $k_{rsd}$ closer to $1x10^3$ $L\text{-}mol^{-1}s^{-1}$ than R3 in our simulations (Figure S9). (**B**) Denaturing gel electrophoresis results for the gates in the cascades. The samples were run on the same gel. The white spacing between images denotes samples that were not in consecutive lanes. (**C**) Reporter kinetics of each cascade. Blue lines indicate simulation results using effective $k_{rsd}$ $1x10^3$ $L{\cdot}mol^{-1}s^{-1}$, $3x10^2$ $L{\cdot}mol^{-1}s\text{-}1$, $1x10^3$ $L{\cdot}mol^{-1}s^{-1}$, $5x10^3$ $L{\cdot}mol^{-1}s^{-1}$ for G{v1,u2r}, G{u5,v1}, G{v4,u5}, G{u3,v4}, respectively. These rate constant values are consistent with previous measurements of gates with these input domains in other contexts (Supporting Note 6). In each cascade, gates were present at 25 nmol/L and R{u2} was present at 500 nmol/L. The input that initiated each cascade was present at 50 nmol/L. See Methods for additional experimental concentrations, simulation parameters, and gel image processing.

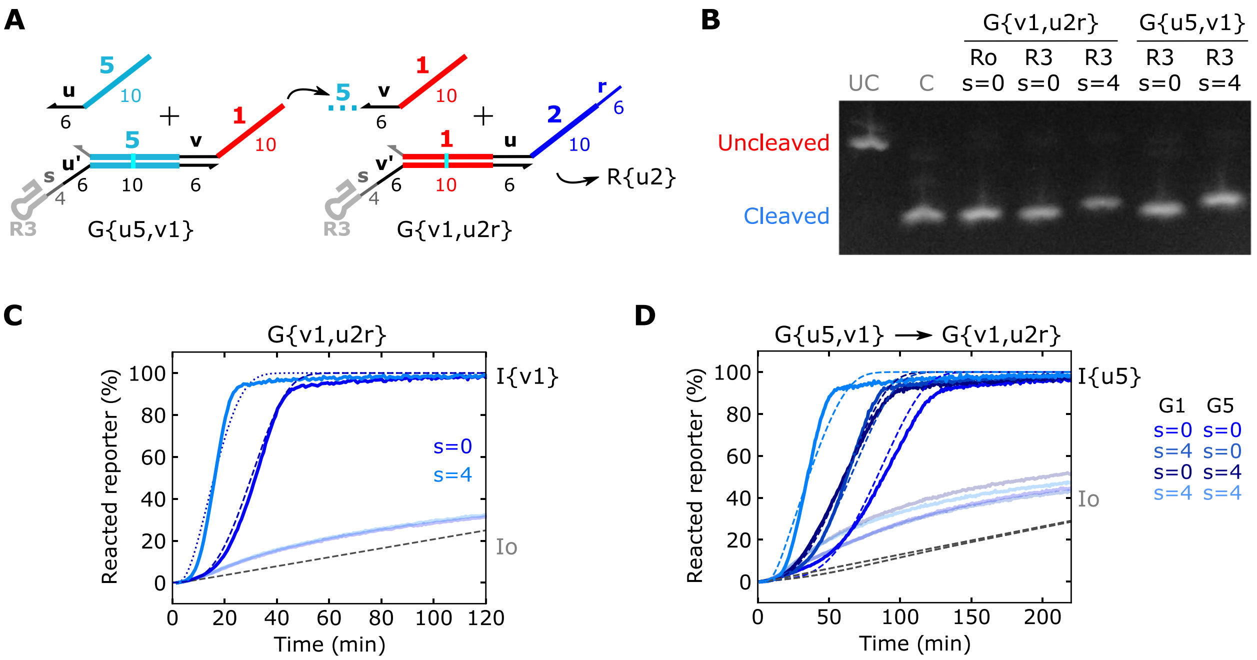

**Figure S14:** Characterization of gates with four-base spacers. (**A**) Schematics of gates with spacers (*s*) shown. (**B**) Denaturing gel electrophoresis results of the gates indicated above the gels. UC and C are size markers described in Figure 2 of the main text. (**C**,**D**) Reporter kinetics for one- and two-layer cascades, (C) and (D), respectively, composed of gates with and without spacers. Dashed lines are simulation results with gates with no spacer (s=0) using $k_{rsd} = 10^3$ $L{\cdot}mol^{-1}s^{-1}$, and gates with four-base spacers (s=4) using $k_{rsd} = 10^5$ $L{\cdot}mol^{-1}s^{-1}$. Gate and input templates were present at 25 nmol/L. R{u2} was present at 500 nmol/L. See Methods for additional experimental concentrations, simulation parameters, and gel image processing.

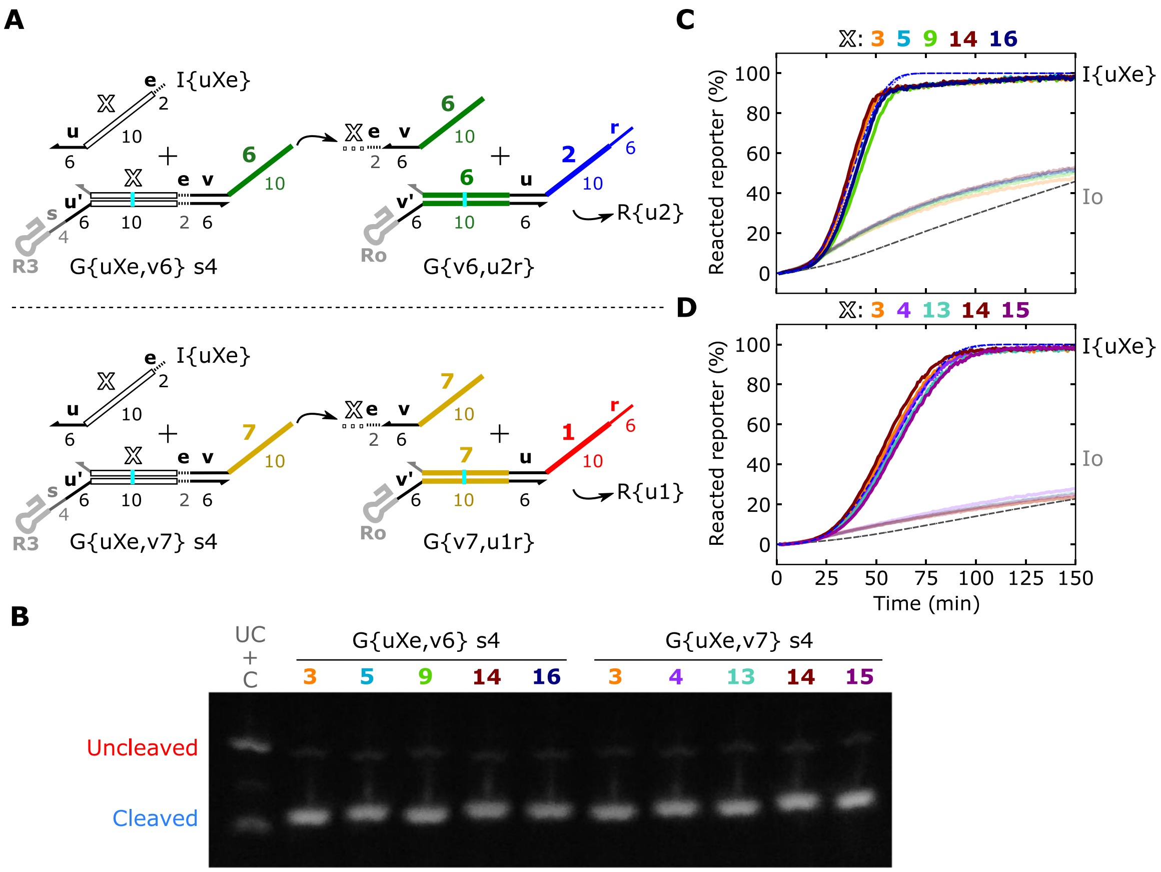

**Figure S15:** Two-layer cascades with four-base spacers (*s*). (**A**) Schematic of the two-layer cascades tested. The white *X* in the schematics denotes the domain of the gate that was varied. (**B**) Denaturing gel electrophoresis results of the gates indicated above the gels. UC and C are size markers described in Figure 2 of the main text. (**C**,**D**) Reporter kinetics of the gates indicated in (A). The line colors in the plots correspond to the domain number colors in the schematics in (A). The blue dashed lines represent simulation results, In the simulations, gates in the first layer (G{v6,u2r} and G{v7,u2r}) used $k_{rsd} = 10^3$ L·mol$^{-1}$s$^{-1}$, and gates in the second layer (G{uXe,v6} and G{uXe,v6}) used $k_{rsd} = 10^5$ L·mol$^{-1}$s$^{-1}$. The shaded region of the simulation results spans $5k_{rsd}$ to $k_{rsd}/5$ for gates in the second layer of the cascades. Note the higher strand displacement rate constant of the second layer makes the kinetics much tighter across sequences because RNA strand displacement is no longer the rate limiting step in the second layer. Gate and input templates were present at 25 nmol/L. R{u2} was present at 500 nmol/L. See Methods for additional experimental concentrations, simulation parameters, and gel image processing.

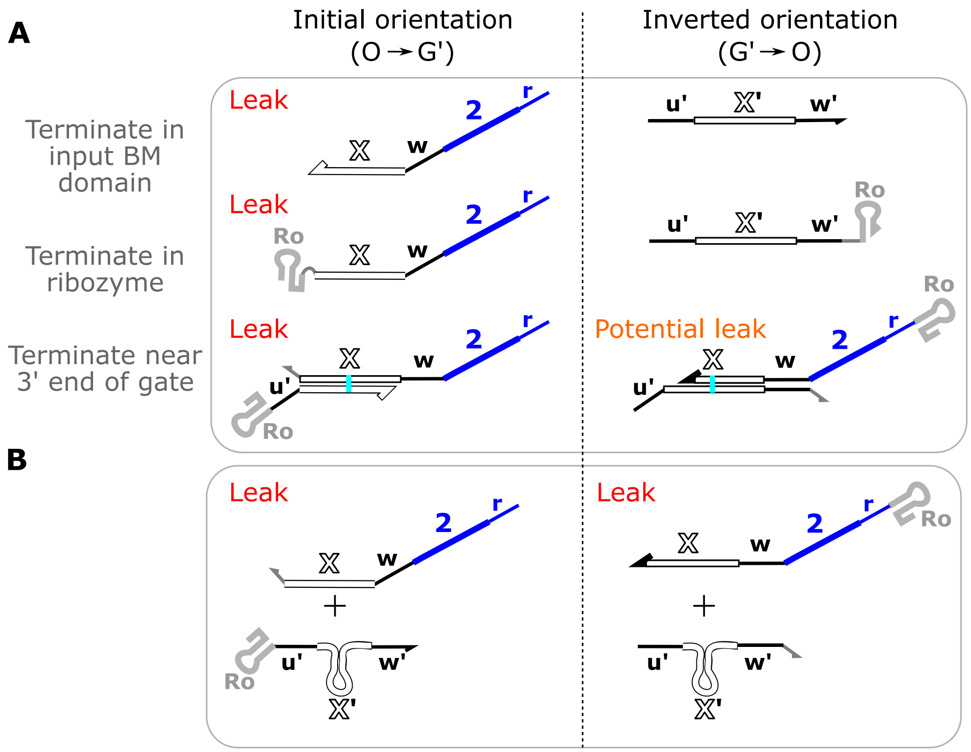

**Figure S16:** Possible leak products for gates transcribed in initial (O → G′) and inverted orientation (G′ → O). (**A**) Truncated products that result from premature transcription termination. With the initial transcription orientation, most truncated products leave the entire output domain exposed for downstream leak. With the inverted transcription orientation, truncated products should not result in downstream leak. The bottom truncated product for gates transcribed in the inverted orientation could result in leak if the truncated *X* domain is short enough to dehybridize, effectively releasing the output strand. (**B**) For both transcription orientations, a misfolded G′ strand could result in free output strands that introduce leak. Note any misfold that does not sequester the *w* toehold should behave similarly.

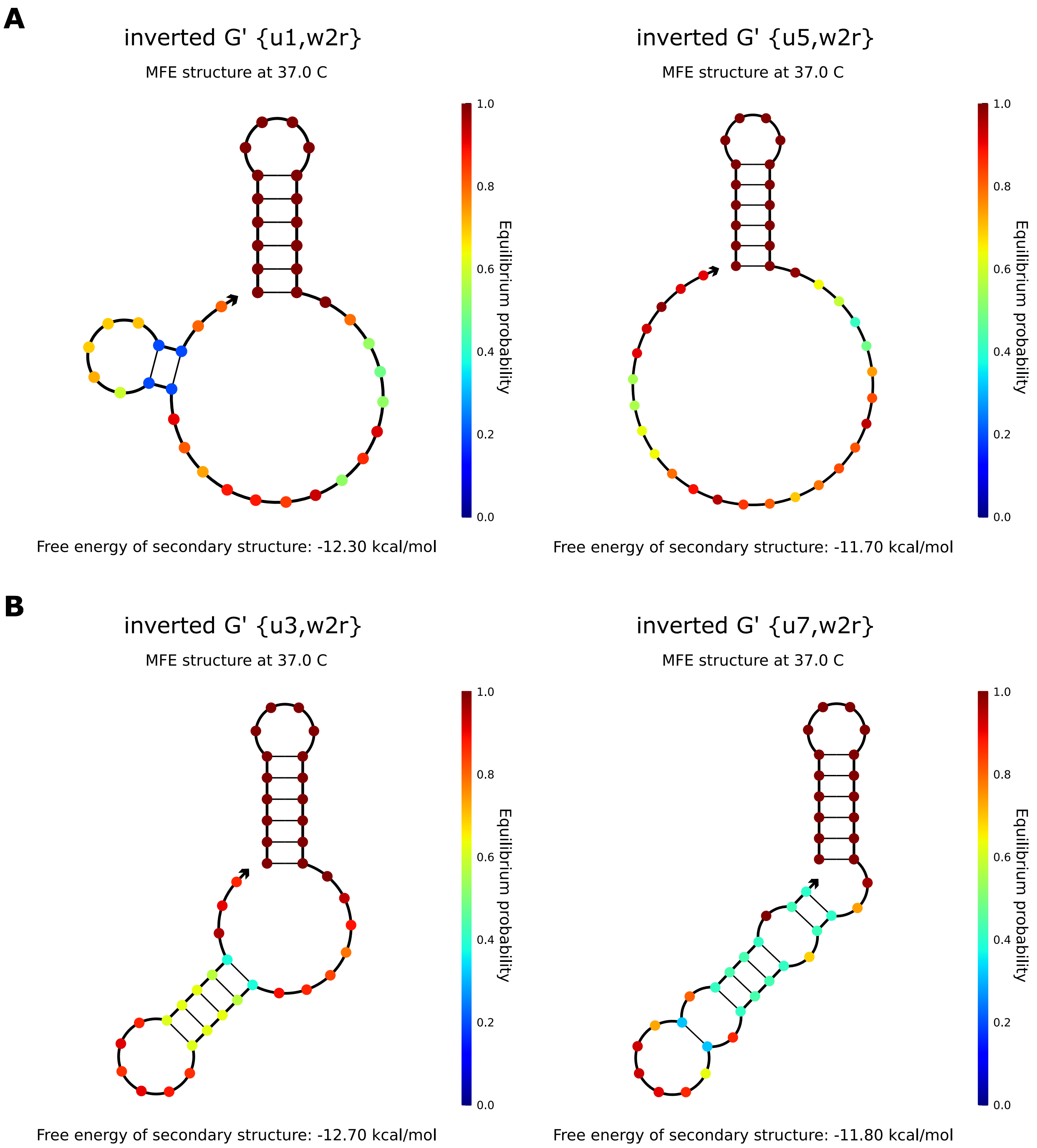

**Figure S17:** Predicted secondary structure of G′ strands of ctRSD gates with inverted transcription orientation. Predictions conducted in NUPACK 3.2.2[3] with default parameters at 37 ºC.

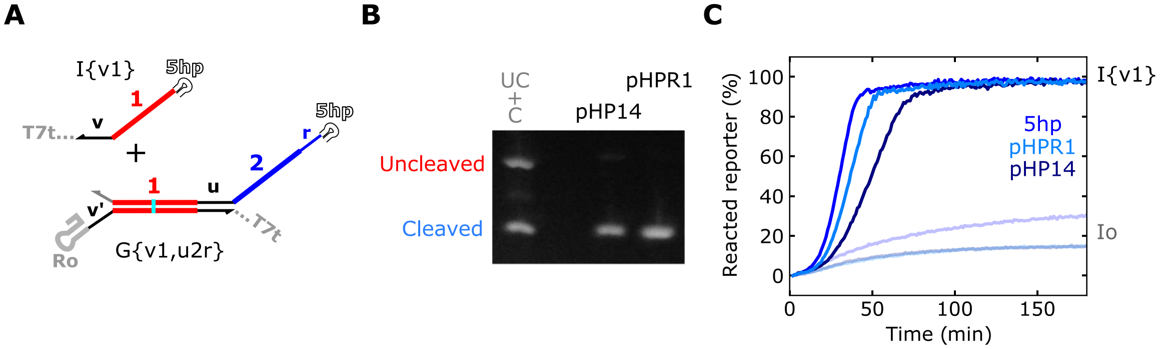

**Figure S18**: Characterization of G{v1,u2r} with different 5’ hairpin sequences. (**A**) Schematic of a gate with different 5’ hairpin sequences. pHP14 is from Ref[4] and pHPR1 is from Ref[5]. Three unpaired guanine bases were also present to the 5’ end of pHP14 and pHPR1 to promote efficient transcription by T7 RNAP. (**B**) Denaturing gel electrophoresis results. UC and C are size markers described in Figure 2 of the main text. (**C**) Reporter kinetics for gates with different 5’ hairpin sequences. The blue dashed lines represent simulation results with the shaded region spanning $5k_{rsd}$ to $k_{rsd}/5$. Gate and input templates were present at 25 nmol/L. R{u2} was present at 500 nmol/L. Differences in reporter kinetics could be related to differences in the transcription rates for these different hairpin sequences[6]. See Supporting Note 3 for sequence schematics of 5’ hairpins. See Methods for additional experimental concentrations, simulation parameters, and gel image processing.

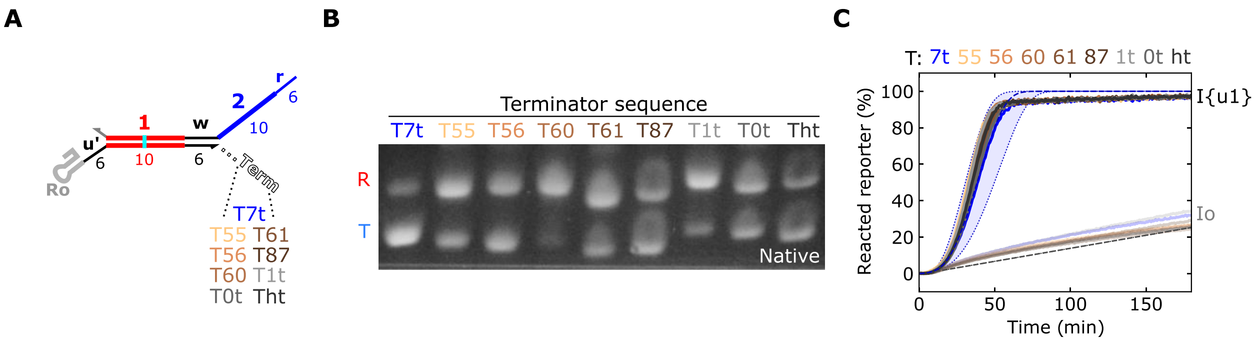

**Figure S19**: Characterization of G{u1,w2r} with different terminator sequences. (**A**) Schematic of G{u1,w2r} with different terminator sequences. (**B**) Native gel electrophoresis results of G{u1,w2r} with the terminator sequences indicated above the gel. In these experiments the DNA templates were PCR amplified such that 52 bases were present downstream of the terminator sequence (see Supporting File S2). This way readthrough transcripts (R) would have a much higher molecular weight than terminated (T) transcripts. (**C**) Reporter kinetics for G{u1,w2r} with the terminator sequences indicated above the plot. The blue dashed lines represent simulation results with the shaded region spanning $3k_{rsd}$ to $k_{rsd}/3$. Gate and input templates were present at 25 nmol/L. R{w2} was present at 500 nmol/L. See Supporting Note 3 for sequence schematics of each terminator. See Methods for additional experimental concentrations, simulation parameters, and gel image processing.

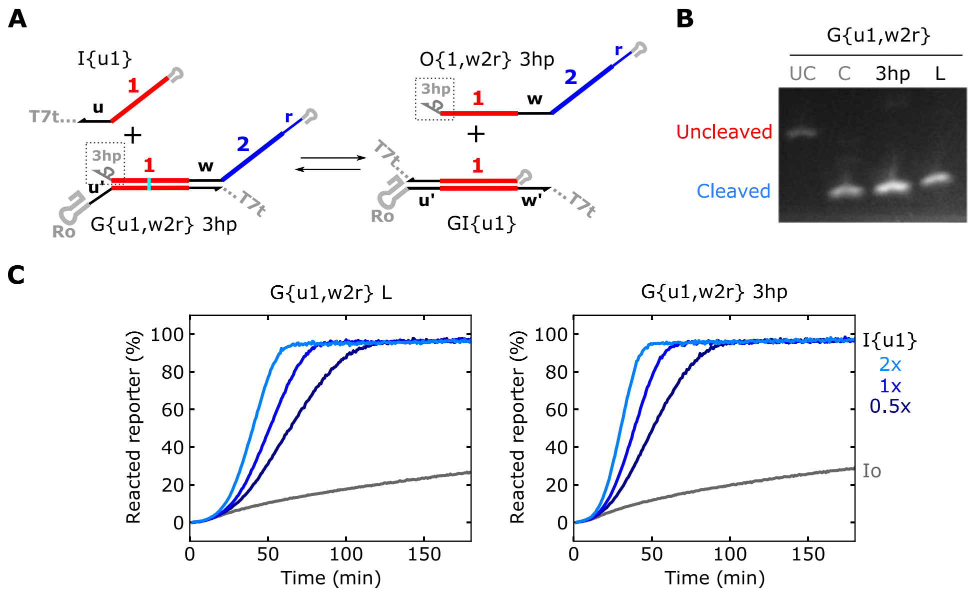

**Figure S20:** Characterization of G{u1,w2r} with a hairpin at the 3' end of the output strand. (**A**) Schematic of the modified gate investigated in this figure. The 3hp sequence was: 5'UCGUCGACUUUCGAGUUGACGUAC. (**B**) Denaturing gel electrophoresis results, where L indicates G{u1,w2r} with the 5'UUC linker sequence. UC and C are size markers described in Figure 2 of the main text. (**C**) Reporter kinetics for gates indicated above the plots. Gate templates were present at 25 nmol/L and the input template was present at either 50 nmol/L (2x), 25 nmol/L (1x), or 12.5 nmol/L (0.5x). R{w2} was present at 500 nmol/L. See Methods for additional experimental concentrations, simulation parameters, and gel image processing.

### Supporting Note 1: DNA template ordering and preparation

#### DNA template ordering

DNA templates encoding for ctRSD gates and inputs were ordered as eBlocks from Integrated DNA technologies, Inc (IDT). At the time of this study, IDT required eBlock sequences be at least 300 bases long. To meet this length requirement for ctRSD components, additional sequences were appended upstream and downstream of the region encoding for the ctRSD components (which we will term flanking sequences). Unless otherwise stated, only the region of the eBlock encoding for the ctRSD component was exponentially amplified during PCR. Additionally, to reduce the complexity of synthesis, a G-U wobble base pair was introduced into the double stranded branch migration domain of the gates to keep complementary sequences less than 12 contiguous bases.

Below are example eBlock sequences for a ctRSD input and gate. The uppercase bases represent the region of the sequence that encodes for the ctRSD component. The lowercase bases represent the flanking region sequences appended to meet the 300-base length requirement. The bases highlighted in cyan represent G-U wobble pairings.

The T7 RNA polymerase prompter sequence is shown in pink text.

The underlined sequences represent the primer binding sequences for PCR amplification of the region encoding the ctRSD component. The sequences of the PCR primers (T7fwd and T7rev) are also shown below.

```
I{u1}:
5’gaagtctaacgctgctctgggctaactgtcTTCTAATACGACTCACTATAGGGAGATTCGTCTCCCATCACTTCA
CAACATCACTATAACCCCTTGGGGCCTCTAAACGGGTCTTGAGGGGTTTTTTGctgaaacctcaggcatttgagaag
cacacgctgaaaggaggaactatatccggattggcgactgtgtacttgttataacatctgacagttaaagtcgggag
aataggagccgcaatacacaatttaccgcatctagacttaactgagatattaccatagatgactatatcta
```

```
G{u1,w2r}:
5’gaagtctaacgctgctctgggctaactgtcTTCTAATACGACTCACTATAGGGAGATTCGTCTCCCACTACATCC
ACATACTAATTAACTTACTTCACATTCGGGTCGGCATGGCATCTCCACCTCCTCGCGGTCCGACCTGGGCTACTTCG
GTAGGCTAAGGGAGTGATGTTGTGAAGTGAGTTAATCTATAACCCCTTGGGGCCTCTAAACGGGTCTTGAGGGGTTT
TTTGctgaaaggaggaactatatccggattggcgctgaaacctcaggcatttgagaagcacacgactgtgt
```

```
T7fwd: 5’TTCTAATACGACTCACTATAGGGAG
```

```
T7rev: 5’CAAAAAACCCCTCAAGACCCGTTTAG
```

In principle, the flanking region sequences added to meet the 300-base length requirement for the eBlocks could be any sequence that passes the IDT complexity tests. However, we appended sequences that could be of use for cloning these components into plasmids in the future. The blue lowercase bases are the 30 bases downstream of the T7 terminator sequence in the pETDuet™-1 plasmid. The orange lowercase bases are the 30 bases downstream of the T7 terminator sequence in the pCDFDuet™-1, pCOLADuet™-1, and pRSFDuet™-1 plasmids. The maroon lowercase bases are 30 bases taken from the plasmid backbone upstream of RNA switches in Ref[7]. These sequences could serve as homology domains for Gibson Assembly[8] of ctRSD components into the pDuet™-1 plasmids. The gray lowercase bases, termed excess sequences, have no intended purpose beyond extending the sequence to 300 bases. Other than the exceptions noted below the same flanking region sequences were appended to ctRSD input and gate templates.

Exceptions:

The following components use slightly different excess sequences – after the blue and orange sequences – than other components. This should not influence performance as these sequences should not be exponentially amplified during PCR.

- G{u6,w2r}
- I{u6}
- G{u1e,w2r} 12
- G{u5e,u1e} 12
- I{u1e} 12
- I{u1e} 14
- I{u3e} – I{u17e} with domain e02 (the last base)

Some templates use the sequence 5’ GCGC instead of 5’ TTC directly upstream of the T7 RNAP promoter sequence. Templates with either of these sequences were still PCR amplified using the same T7fwd primer.

The gates with different terminator sequences had a different sequence downstream of the blue pET sequence which was used as a primer binding site for PCR.

### DNA template preparation

DNA templates for ctRSD components are prepared by PCR amplifying eBlock DNA. As described above, the eBlock DNA possesses flanking sequences that should not be exponentially amplified during PCR.

For all the PCRs in this study the following protocol was followed:

1) Add 1.5 uL of 10 ng/µL eBlock DNA into a 200 µL PCR tube (USA Scientific, catalog no. 1402-4700)
2) Prepare the master mix below and then add to the PCR tube containing the eBlock DNA, pipette mix

| Component | Final volume or concentration |
|---|---|
| 2x Phusion MM | 37.5 µL |
| T7fwd primer | 0.5 µmol/L |
| T7rev primer | 0.5 µmol/L |
| Water | X µL to 73.5 µL total |

3) Add samples to a thermocycler and execute the following protocol with the lid temperature set to 105 °C

| Temp | Time | Repeat |
|---|---|---|
| 98 °C | 5 min | 1x |
| 98 °C | 30 s | 30x |
| 60 °C | 30 s | 30x |
| 72 °C | 30 s | 30x |
| 72 °C | 3 min | 1x |
| 4 °C | hold | |

4) Run a spin column PCR clean up with a QIAquick PCR Purification Kit (catalog no. 28104)
   a. Add 375 µL of Buffer PB to a spin column
   b. Add all 75 µL of the PCR sample to the spin column containing Buffer PB and pipette mix thoroughly (10 to 20 times)
   c. Spin column at 13,000 rpm for 1 min, discard flow through
   d. Add 750 µL of Buffer PE to the spin column
   e. Spin column at 13,000 rpm for 1 min, discard flow through
   f. Spin column at 13,000 rpm for 1 min to remove any last Buffer PE, discard flow through
   g. Place spin column insert into a fresh collection tube
   h. To elute the PCR product, add 50 µL of Buffer EB to the center of the spin column
   i. Let the column sit for 1 min to 2 min at room temperature
   j. Spin column at 13,000 rpm for 1 min, discard spin column
   k. Measure the concentration of the eluted PCR product

This protocol typically yields (65 to 80) ng/µL of gate template and (30 to 45) ng/µL of input template based on A260 measurements of undiluted samples taken on a DeNovix D-11 Series Spectrophotometer. For the lengths of the gate and input PCR products this equates to ~ (500 – 650) nmol/L.

The above protocol was developed previously without much optimization[9] and carried over into this study. Late in the study we found that our PCR conditions resulted in two distinct higher molecular weight products for both gate and input templates (Figure S21). We found decreasing the concentration of eBlock DNA used in the PCR reduced the fraction of higher molecular weight side products without a marked decrease in the concentration of purified PCR products for 10-, 25-, and 50-fold dilutions (Figure S21). Based on these results, **we suggest reducing the PCR eBlock concentration 10- to 25-fold to 20 pg/µL or 8 pg/µL to reduce side products.** If a higher concentration of final PCR product is desired, then the primer concentrations can be increased to 1 µmol/L.

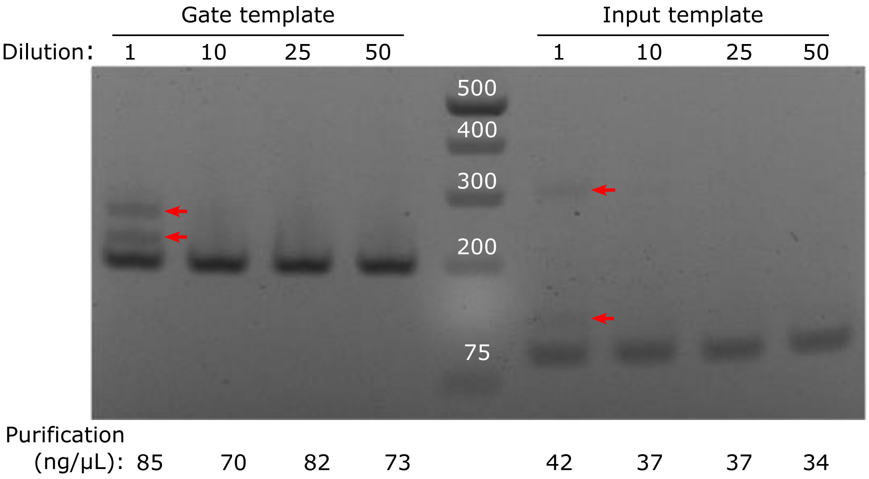

**Figure S21:** Native gel electrophoresis results of eBlock PCR products with decreasing DNA concentrations. Gate template DNA encoded for G{$t_5$1,u2r} and input template DNA encoded for I{$t_5$1}. Dilutions: 1) 0.2 ng/µL – 10 ng total, 10) 0.02 ng/µL – 1 ng total, 25) 0.008 ng/µL – 0.4 ng total, 50) 0.004 ng/µL – 0.2 ng total. The numbers below the gel image are the measured DNA concentrations of DNA products after PCR clean-up. The red arrows indicate undesired side products. Note all dilutions yield similar DNA concentrations. Samples were prepared by mixing 5 µL of unpurified PCR product with 15 µL of water. All 20 µL of diluted PCR product was run on a 2% Agarose E-gel pre-stained with ethidium bromide (Invitrogen: catalog no. G501802) and run for 30 min using an E-gel powerbase. The ladder was a GeneRuler 1 kb plus DNA ladder (Invitrogen: catalog no. SM1331) with the number of base pairs labeled in white. Any brightness or contrast adjustments to aid in visualization were applied uniformly to the entire gel image.

**Supporting Note 2: Description of kinetic simulations**

The chemical reactions of the ctRSD kinetic model relevant to this study are shown in Figure S22. As discussed in the Methods of the main text, a user-friendly ctRSD simulation package was developed in Python to simulate these reactions. This package includes many additional chemical reactions pertinent to more complicated ctRSD circuits than investigated in this study. The full documentation of the ctRSD simulation package can be found at: https://ctrsd-simulator.readthedocs.io/en/latest/index.html.

The model uses a first order approximation of enzyme kinetics for RNA transcription. From this approximation the transcription rate is modeled as linearly proportional to the template concentration, *e.g.*, $k_{txn}$*[G{i,j}$_{temp}$] where $k_{txn}$ is the apparent first order rate constant for production of RNA from the DNA template for G{i,j}. Additionally, leak transcription was modeled as previously described (Supplementary Section 5 of Ref[9]).

The expected rate constants values used in the simulations were determined previously (Supplementary Section 5 of Ref[9]) and are presented in Table S1. Note the nomenclature has changed slightly from the previous description. In this paper, the *u* and *w* toeholds correspond to the *a* and *b* toeholds, respectively, from the previous description[9].

In a typical simulation, we assumed the rate constant values in Table S1. But as we changed domain sequences within gates, we observed differences in reporter kinetics, suggesting the rate constant values can differ slightly across sequences. We verified these differences were reproducible across technical replicates and PCR preparations (Supporting Note 4). Nucleic acid strand displacement rates are known to vary up to an order of magnitude depending on the sequences[10], so we expected the reporter kinetics to vary within this range across gate sequences. To assess whether our experimental results were consistent with these previous observations, we conducted simulations in which $k_{rsd}$ was increased or decreased by a factor of 4 to 5 from the expected value and compared these simulation results to the spread in reporter kinetics observed across gates with different input sequences in experiments. We used this analysis as a heuristic to assess gate performance, deeming gates with reporter kinetics that fall within this range as having the expected performance and any gates that fall outside of this range as having poor performance. For one-layer ctRSD cascades the acceptable range was $4k_{rsd}$ to $k_{rsd}/4$ (Figure S23) and for two-layer ctRSD cascades the acceptable range was $5k_{rsd}$ to $k_{rsd}/5$ (Figure S23).

Considering Figure S22 there are rate constants other than $k_{rsd}$ that could influence the measured reporter kinetics if they varied from sequence to sequence. For example, sequence specific differences in $k_{txn}$ for an input or a gate or $k_{rz}$ for a gate. So, it is possible that slight variations in these rate constants across sequences contribute to the differences in measured reporter kinetics. The majority of ctRSD components in this study use the same 16-base initiation sequence in an effort to maintain the same transcription rate across sequences[6], so we do not expect $k_{txn}$ to be strongly sequence dependent. However, slight differences in cleavage efficiency, *i.e.*, the fraction of transcripts that cleave, across sequences would likely result in lower effective $k_{txn}$ values. It is worth noting that changes in $k_{rsd}$ and $k_{txn}$ both produce shaded regions with similar shapes (Figure S23), making it hard to distinguish these two rate constants based on our measurements. Considering the ribozyme cleavage rate, our simulations suggest that higher $k_{rz}$ values cannot explain reporter kinetics that are faster than with the expected $k_{rsd}$ (Figure S23). Further, gates with a 10-fold lower $k_{rz}$ value would show poor cleavage in our denaturing gel electrophoresis assay and would be characterized as bad performers based on that metric. For gates that cleave well, we do not expect small differences in $k_{rz}$ across sequences to dramatically influence reporter kinetics. For a set of gates with the same output domain we do not expect $k_{rep}$ to differ substantially across input sequences. Lastly, under our effectively irreversible reporting conditions, $k_{rev}$ would need to be increased >10-fold compared to the expected value to change reporter kinetics[9].

There are also mechanisms that are not captured in our simple model that could influence reporter kinetics. For example, gate transcripts may initially misfold during transcription and then, after the full transcript is produced or cleavage has occurred, rearrange slowly into the desired structure for strand displacement[1,2,11]. This mechanism could effectively reduce the $k_{rsd}$ value or be modeled as reducing $k_{txn}$ because less reactive transcript is produced per unit time.

| Schematic | Reaction |
|---|---|
| **Transcription** 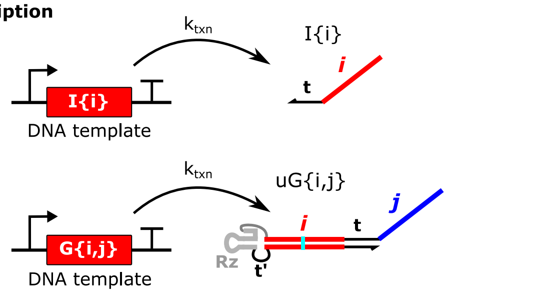  | $\mathrm{I\{i\}_{temp}} \xrightarrow{k_{txn}} \mathrm{I\{i\}}$<br><br>$\mathrm{G\{i,j\}_{temp}} \xrightarrow{k_{txn}} \mathrm{uG\{i,j\}}$ |
| **Ribozyme cleavage** 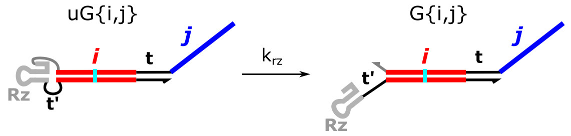  | $\mathrm{uG\{i,j\}} \xrightarrow{k_{rz}} \mathrm{G\{i,j\}}$ |
| **RNA strand displacement with inputs** 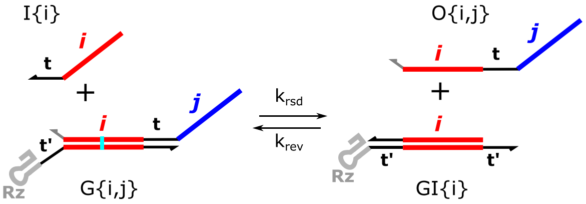  | $\mathrm{I\{i\}+G\{i,j\}} \xrightarrow{k_{rsd}} \mathrm{O\{i,j\}+GI\{i\}}$<br><br>$\mathrm{O\{i,j\}+GI\{i\}} \xrightarrow{k_{rev}} \mathrm{I\{i\}+G\{i,j\}}$ |
| **RNA strand displacement with outputs** 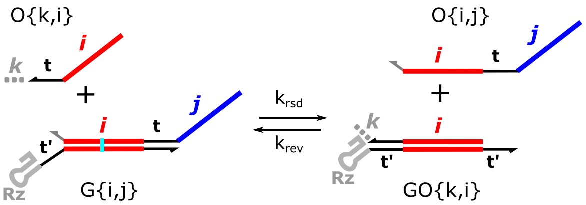  | $\mathrm{O\{k,i\}+G\{i,j\}} \xrightarrow{k_{rsd}} \mathrm{O\{i,j\}+GO\{k,i\}}$<br><br>$\mathrm{O\{i,j\}+GO\{k,i\}} \xrightarrow{k_{rev}} \mathrm{O\{k,i\}+G\{i,j\}}$ |
| **Output reporting** 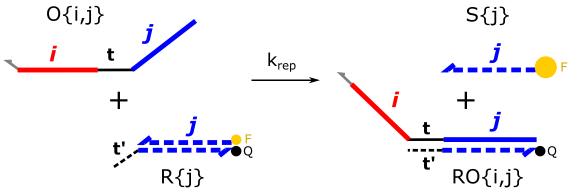  | $\mathrm{O\{i,j\}+R\{j\}} \xrightarrow{k_{rep}} \mathrm{S\{j\}+RO\{i,j\}}$ |
| **Leak transcription** 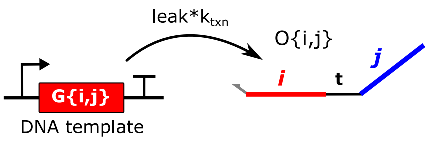  | $\mathrm{G\{i,j\}_{temp}} \xrightarrow{leak*k_{txn}} \mathrm{O\{i,j\}}$ |

**Figure S22:** Chemical reactions in the ctRSD kinetic model. *i*,*j*,*k* represent input and output domains. For simplicity, the model is implemented with a universal toehold that ignores the toehold connectivity of components. Consequently, the toehold names are dropped from the nomenclature. The expected rate constant values are presented in Table S1. The rate constants can be customized for individual components but for simplicity are shown as single constants above.

**Table S1:** The expected rate constants used in most simulations. Changes to these values in for specific components in simulations is noted in figure captions. See Supplementary Section 5 of Ref[9] for details on how these values were obtained. Note the *u* and *w* toeholds correspond to the *a* and *b* toeholds, respectively, from the previous description. Gates with the *w* output toehold, for example, G{-,w2r} were given a lower $k_{rev}$ value than gates with a *u* or *v* output toehold, *i.e.*, G{-,u2r}, G{-,v1}, etc.). This was done because the *w* toehold is weaker (lower GC content) than the *u* and *v* toeholds.

| Rate constant | Value |
|---|---|
| $k_{txn}$ | Calibrated in each experiment (0.009-0.024) $s^{-1}$ |
| $k_{rz}$ | 0.00417 $s^{-1}$ |
| $k_{rsd}$ | $1x10^3$ $L \cdot mol^{-1}s^{-1}$ |
| $k_{rev}$ (*u*,*v* toeholds) | $0.27x10^3$ $L \cdot mol^{-1}s^{-1}$ |
| $k_{rev}$ (*w* toehold) | 5 $L \cdot mol^{-1}s^{-1}$ |
| $k_{sd}$ | $1x10^4$ $L \cdot mol^{-1}s^{-1}$ |
| leak | 0.03 |

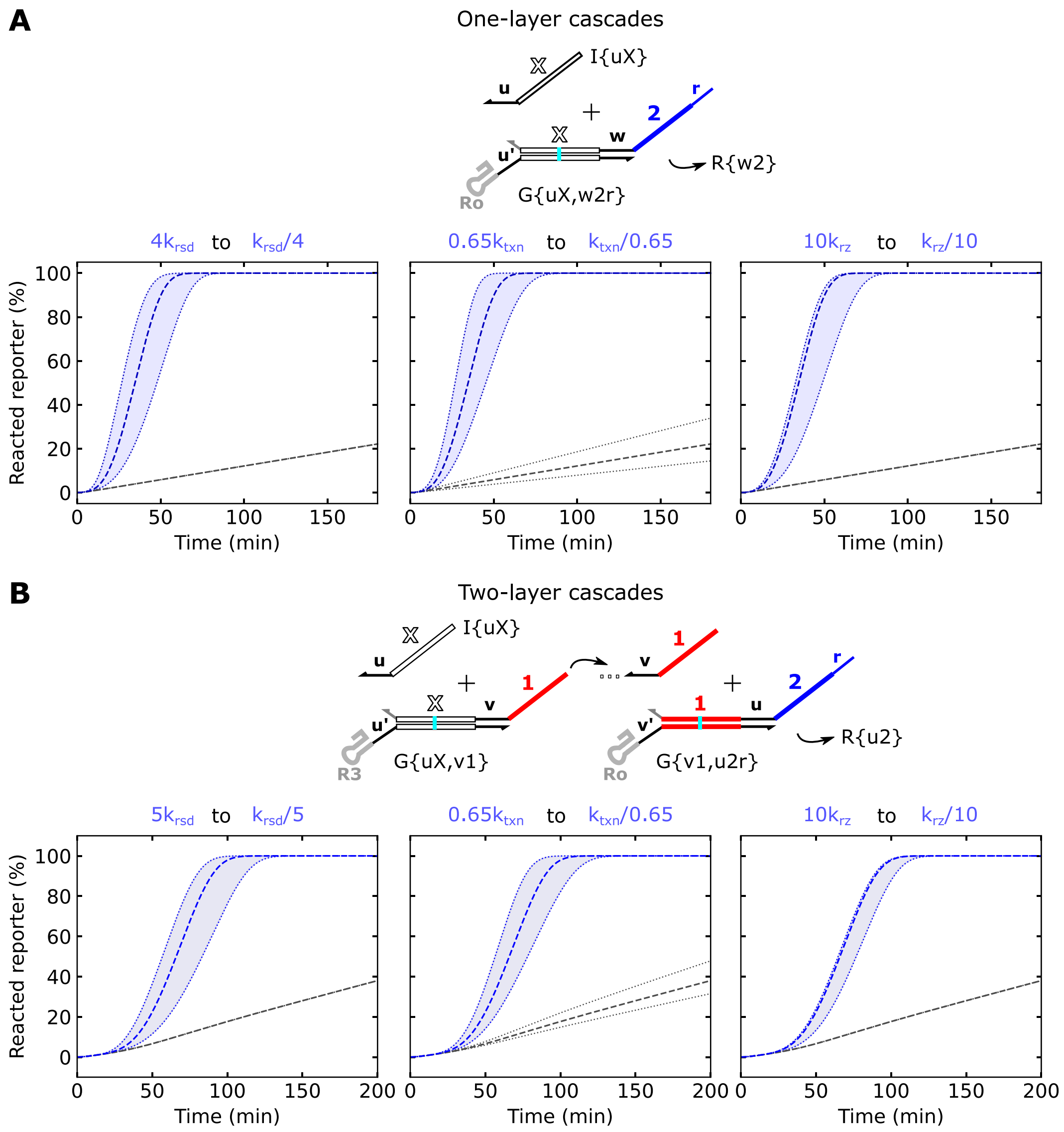

**Figure S23:** Simulation results for one- and two-layer ctRSD cascades with ranges of $k_{rsd}$, $k_{txn}$, or $k_{rz}$ values. Schematics above the plots show the simulated systems. The dashed blue lines represent simulation results with the expected rate constant values. The blue shaded region between dotted lines denotes simulations spanning the rate constant range indicated above the plots. The gray lines represent simulations results without inputs. For simulations with $k_{txn}$ varied, the dotted lines represent the high and low $k_{txn}$ values because leak is proportional to transcription rate in the model. The white *X* domains represent different sequences and the rate constants for those components with *X* domains were varied in the simulations as follows: For one-layer cascades (**A**), $k_{rsd}$ and $k_{rz}$ were changed for G{uX,w2r} and $k_{txn}$ was changed for both G{uX,w2r} and I{uX}. For two-layer cascades (**B**), $k_{rsd}$ and $k_{rz}$ were changed for G{uX,v1} only, and $k_{txn}$ was changed for both G{uX,v1} and I{uX}. The expected rate constant values in Table S1 were used for G{v1,u2r} in all simulations.

### Supporting Note 3: Nomenclature and sequence schematics

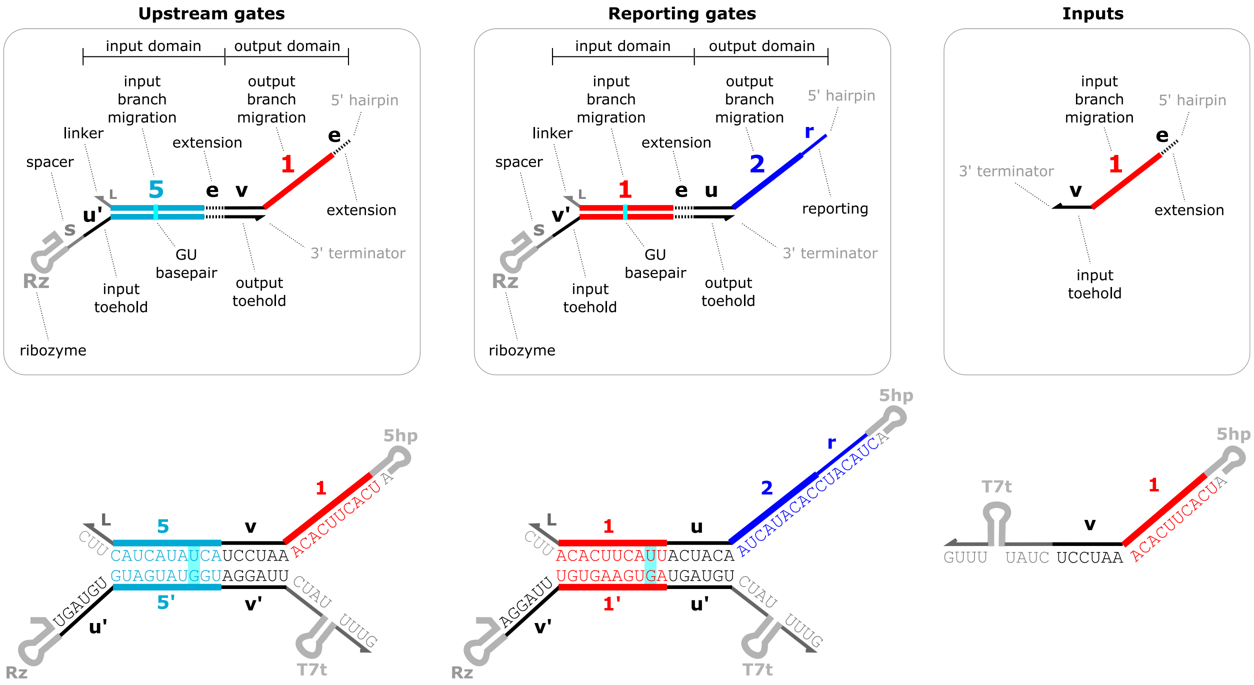

**Figure S24:** Overview of full gate and input anatomy. Most gates tested in this study do not have the *s* or *e* domains so those domains are omitted in the sequence schematics. Cyan shaded base pairs represent G-U wobble base pairs.

| Character | Definition |
|---|---|
| 1, 2, ... 16 | Branch migration domains |
| u, v, w | Toehold domains |
| Rz | ribozyme (Rz: Ro, R3, Rh, etc.) |
| r | reporting domain |
| L | linker |
| s | spacer |
| e | extension domain |

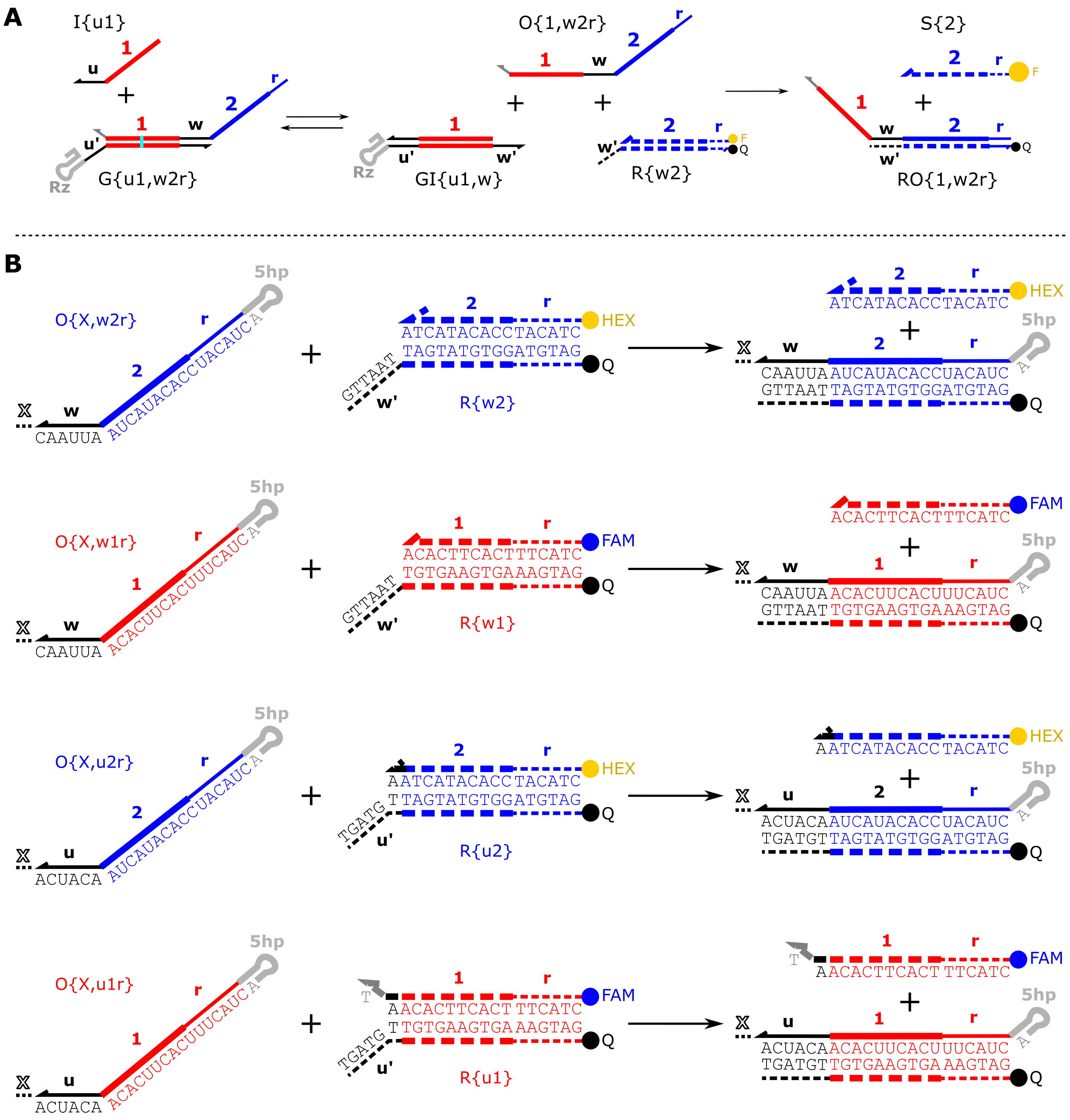

**Figure S25:** Sequence schematics of DNA reporter complexes. RNA and DNA are represented with solid and dashed lines, respectively. The *r* domain is present on the reporters so that they remain hybridized at 37 ºC. The outputs that react with the reporter possess the *r* domain so that the reporting reaction will be effectively irreversible. The reporters with the *w* toehold are 6 bases and reporters with the *u* toehold were shortened to 5 bases to maintain a similar strand displacement rate despite the higher GC content. The fluorophore modified strand of R{u1} has a single unpaired T at the 3’ end. We found the same reporter without this overhang produced artifacts in our fluorescence measurements, *e.g.*, fluorescence would reach a maximum and then slowly decrease to plateau at ~85% the maximum value.

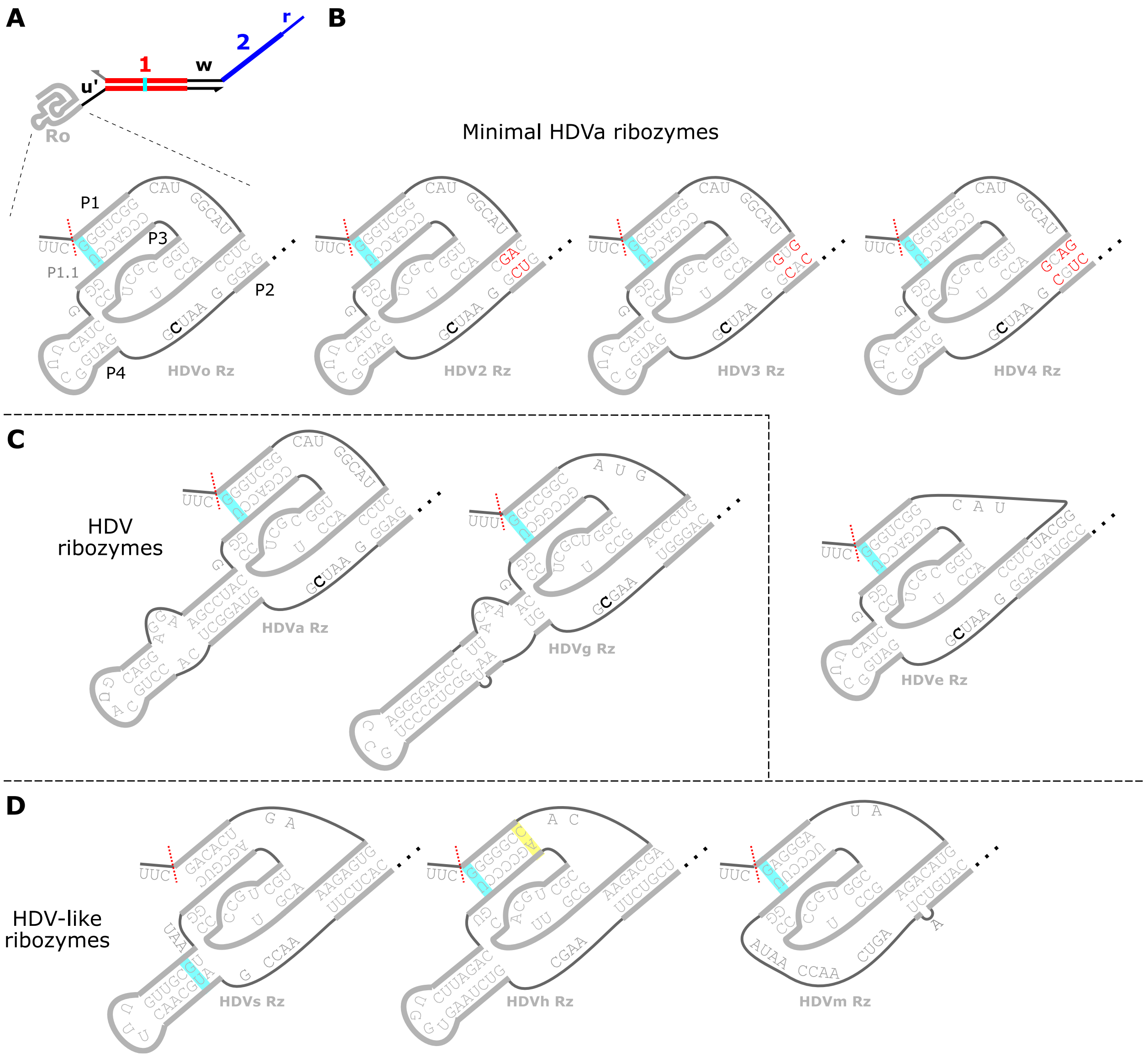

**Figure S26:** Sequence schematics of ribozymes. (**A**) The minimal HDV ribozyme (Ro) primarily used in this study. The P1, P1.1, P2, P3, and P4 helical domains are labeled. The red dashed line indicates the cleavage site. The bold black C is required for catalysis and was changed to a U in the uncleaved gate size markers for gel electrophoresis[12]. Cyan and yellow shaded base pairs represent G-U wobble base pairs and G-A noncanonical base pairs, respectively. This ribozyme is based on the antigenomic HDV ribozyme with a truncated P4 stem[13] (panel C, left). (**B**) Ribozymes derived from the minimal HDV ribozyme (Ro) that were designed to reduce misfolding in different gate sequence contexts. HDV2 to HDV4 have mutations in the P2 helix (red bases) and HDVe has an extended P2 helix. (**C**) The genomic (left) and antigenomic (right) HDV ribozymes as identified in the hepatitis delta virus genome[14,15]. (**D**) Ribozymes that possess the same pseudoknotted fold as the HDV ribozymes but were identified in different species. HDVs was identified in sea urchins (drz-Spur-3)[16], HDVh was identified in humans (CPEB3)[17], and HDVm was identified in human gut microbes (drz-Mtgn-3)[16,18]. The HDVm ribozyme does not possess the P4 helix.

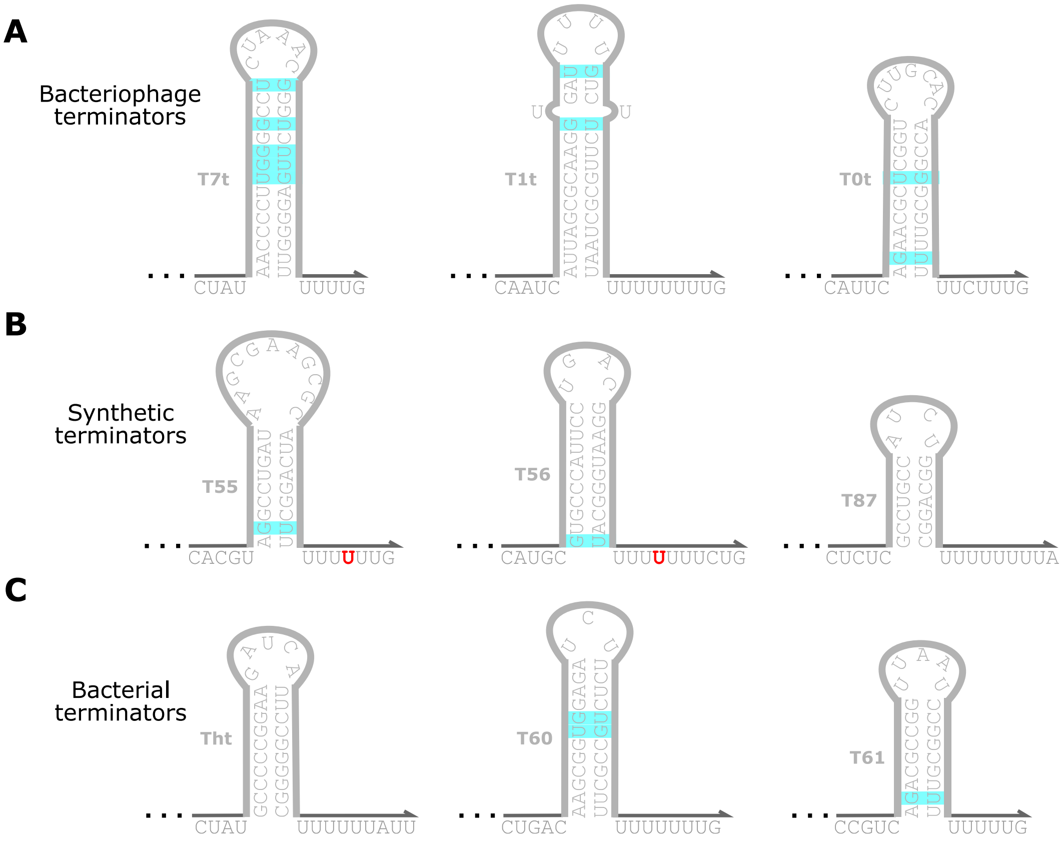

**Figure S27:** Sequence schematics of terminators. Cyan shaded base pairs represent G-U wobble base pairs. T7t was modified from the terminator in the T7 bacteriophage genome[19]. T1t and T0t were modified from the T1 and T0 terminators from lambda phage[20,21]. Tht was modified from the terminator that regulates histidine operon expression in bacteria[22]. T55, T56, T87, T60, T61 were modified from Ref[23]. The sequences to the 5' and 3' of hairpin stems were truncated and/or altered to reduce the possibility of undesired secondary structure with the G{u1,w2r} sequence. Red letters indicate the predominant base for termination[23].

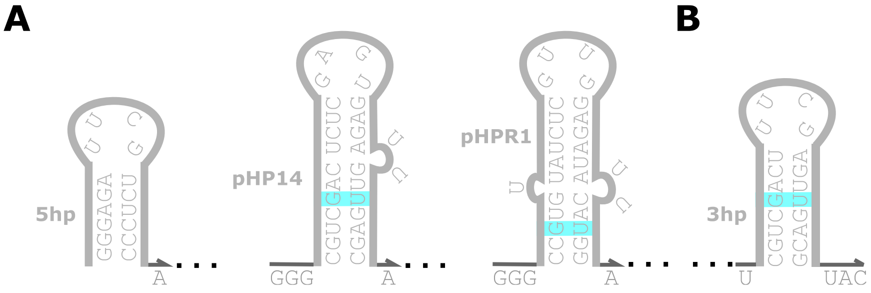

**Figure S28:** Sequence schematics of hairpins used at the 5' (**A**) or 3' (**B**) ends of output strands. pHP14 is from Ref[4] and pHPR1 is from Ref[5]. 3hp was derived from pHP14 with a shortened stem. Cyan shaded base pairs represent G-U wobble base pairs.

**Supporting Note 4: Analysis of reproducibility in DNA reporter assays**

Our experimental results indicate that gates with different input domains can exhibit different reporter kinetics. To confirm that observed differences in reporter kinetics were not due solely to experimental variation, we conducted a study of experimental reproducibility in the DNA reporter assay used to measure ctRSD circuit kinetics. Figure S29A shows experimental workflow for preparing DNA templates for the assay. To assess reproducibility, we conducted two types of replicates that we refer to as technical and biological replicates (Figure S29B). Technical replicates were composed of three repeats of the same experiment on the same day. Each technical replicate used DNA templates from the same PCR preparation and each replicate gate and input were independently pipetted into each reaction well. Biological replicates were conducted from PCRs prepared on different days. Each biological replicate was performed in technical triplicate.

We conducted these replicate experiments for both one-layer cascades (Figure S29C) and two-layer cascades (Figure S29D). For the one-layer cascade we tested G{u1,w2r} and G{u12,w2r}, which had markedly slower reporter kinetics than G{u1,w2r} (Figure 2 of the main text). For the two-layer cascade we tested three gates: one with that had exhibited the expected DNA reporter kinetics (G{u12,v1}), one that had exhibited faster DNA reporter kinetics (G{u3,v1}), and one that had exhibited slower DNA reporter kinetics (G{u16,v1}) (Figure 5 of main text). These trends in reporter kinetics were observed across biological and technical replicates (Figure S29C,D). Further, the technical replicates showed low variability, with a standard deviation of < 6 % from the mean. Together these results reinforce that experimental variation is not likely to account for the substantial differences observed in reporter kinetics across gates with different input sequences.

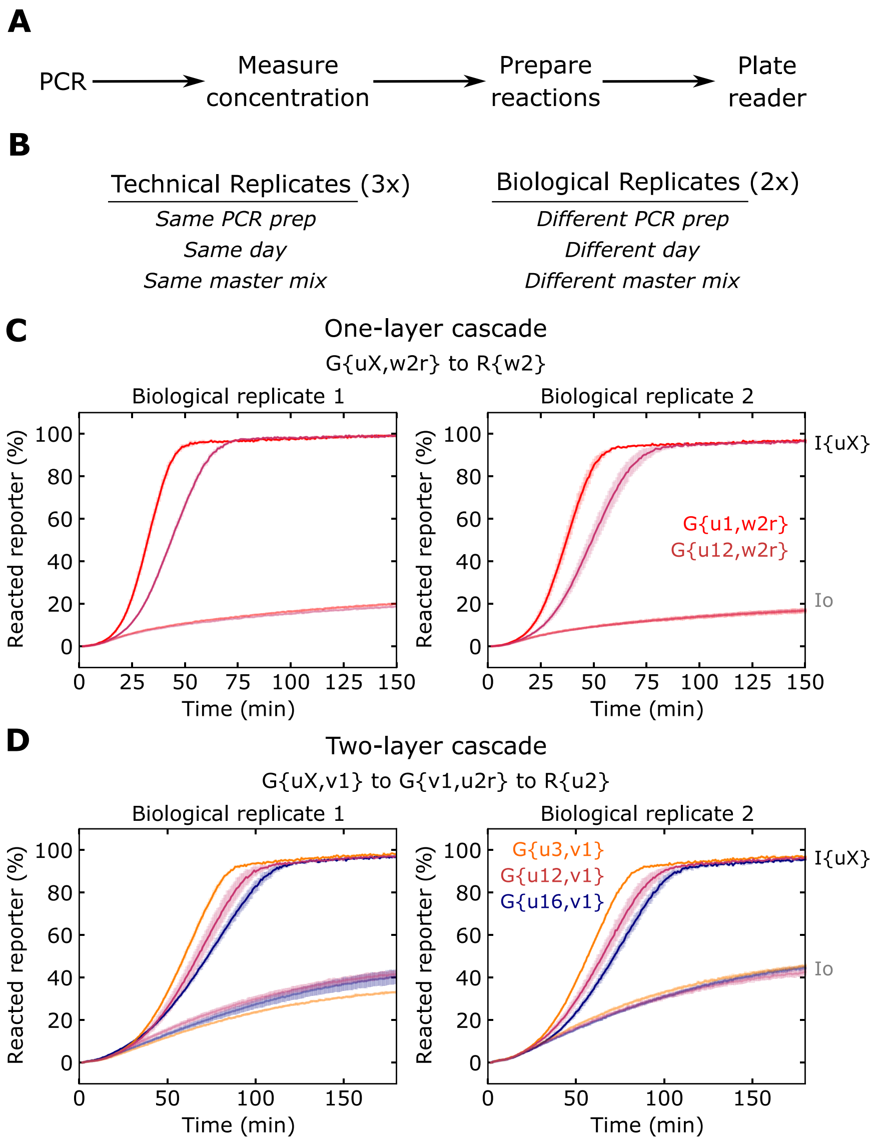

**Figure S29:** Assessing reproducibility of ctRSD circuit experiments. (**A**) Experimental workflow for preparing DNA templates for the reporter assay. (**B**) Description of technical and biological replicates. (**C**) Reporter kinetics for replicates of select one-layer cascades. Shaded regions represent one standard deviation. Biological replicate 1 standard deviation: G{u1,w2r}: < 3 %, G{u12,w2r}: < 2 %. Biological replicate 2 standard deviation: G{u1,w2r}: < 5 %, G{u12,w2r}: < 6 %. The relative trends of reporter kinetics align with the results in Figure 2A of the main text. Gate and input templates were present at 25 nmol/L and 50 nmol/L, respectively. R{w2} was present at 500 nmol/L. T7 RNAP was present at 3 U/µL. (**D**) Reporter kinetics for replicates of select two-layer cascades. Shaded regions represent one standard deviation. Biological replicate 1 standard deviation: G{u3,v1}: < 2 %, G{u12,v1}: < 6 %, G{u16,v1}: < 4 %. Biological replicate 2 standard deviation: G{u3,v1}: < 2 %, G{u12,v1}: < 6 %, G{u16,v1}: < 4 %. The relative trends of reporter kinetics align with the results in Figure 5C of the main text. Gate and input templates were present at 25 nmol/L, respectively. R{u2} was present at 500 nmol/L. T7 RNAP was present at 3 U/µL. In the plots, the solid lines show the mean of the three replicates; semi-transparent error bars represent one standard deviation.

**Supporting Note 5: Sequence schematics of possible misfolded gate structures**

Below are sequence schematics showing possible misfolded secondary structures for some of the poor performing gates identified in this study. Because these misfolds likely arise during cotranscriptional folding, it is difficult to use thermodynamics-based secondary structure prediction software (such as NUPACK) to analyze these structures. These software packages typically predict the desired structure for all gates because the desired structure is designed to be the minimum free energy structure. Further, most thermodynamic software packages cannot predict the pseudoknotted structure of the ribozymes used in this study. This makes it difficult to use these packages to accurately predict disruption of the ribozyme fold. Therefore, the secondary structures presented are not the result of any strict thermodynamic predictions, but rather, the result of simply analyzing sequence complementarity. These secondary structures are consistent with the performance measurements for the gates, *i.e.*, reporter kinetics and cleavage efficiency. When analyzing gates with poor cleavage, we focused on the disruption of the P2 helix of the ribozyme because we deemed the neighboring single-stranded input toehold of the gates as a likely region to nucleate misfolds. It is possible that other helices of the ribozyme could be disrupted. It is also important to note that for any ctRSD gate with poor performance, an ensemble of different structures likely exist, some of which may be the desired structure.

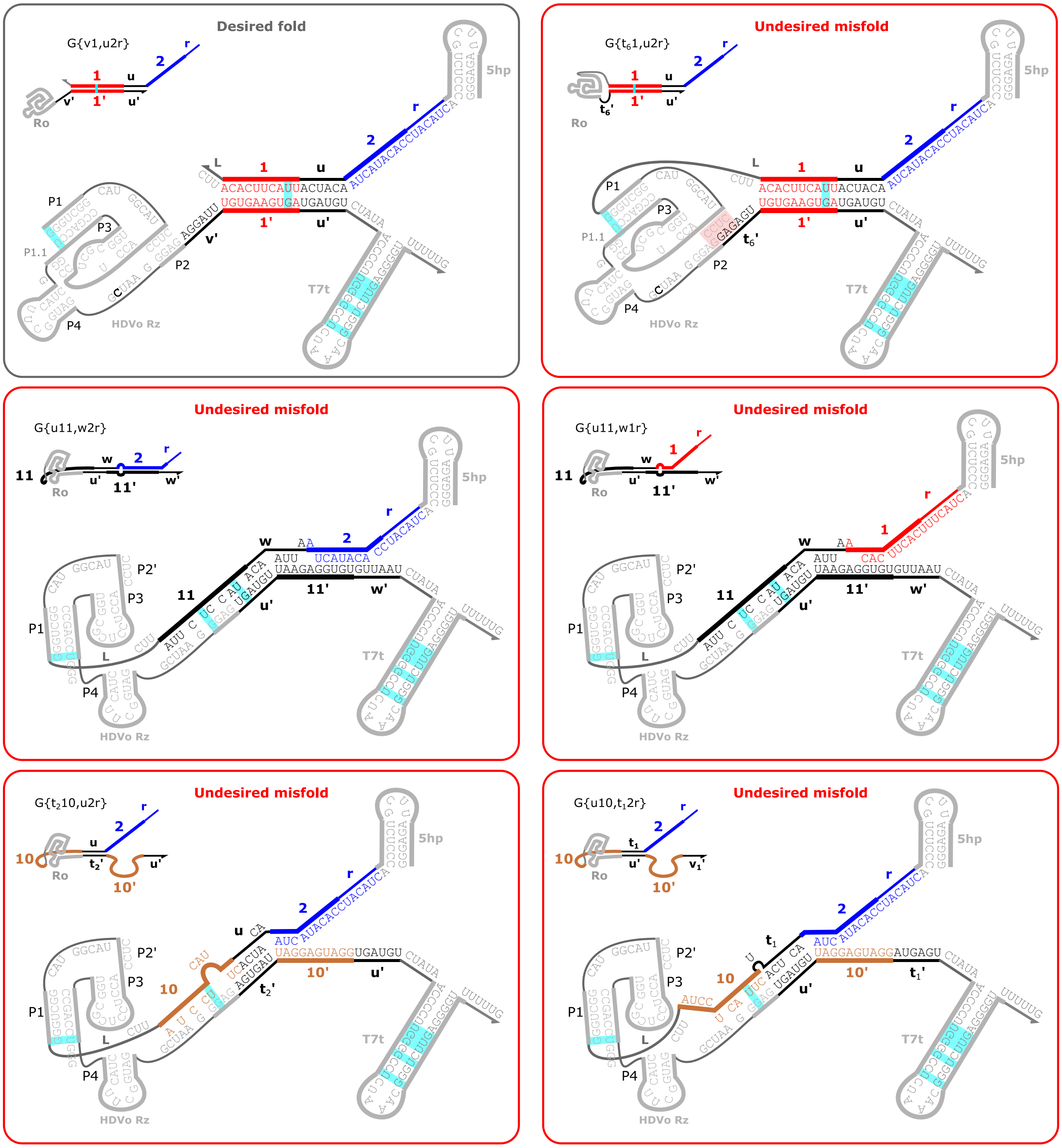

**Figure S30:** Possible misfolded structures of gates with poor performance that did not have repeated input-output toehold sequences. G{v1,u2r} and G{$t_6$1,u2r} were characterized in Figure 3 of the main text. The cleavage activity of G{$t_6$1,u2r} can be recovered with the R3 and Rh ribozymes (Figure S10). G{u11,u2r} and G{u11,u1r} were characterized in Figure 2 of the main text. The cleavage activity of G{u11,u2r} can be recovered with the R3 and Rh ribozymes (Figure 4 of the main text). G{$t_2$10,u2r} and G{u10,$t_6$2r} were characterized in Figure S3. Cyan shaded base pairs represent G-U wobble base pairs.

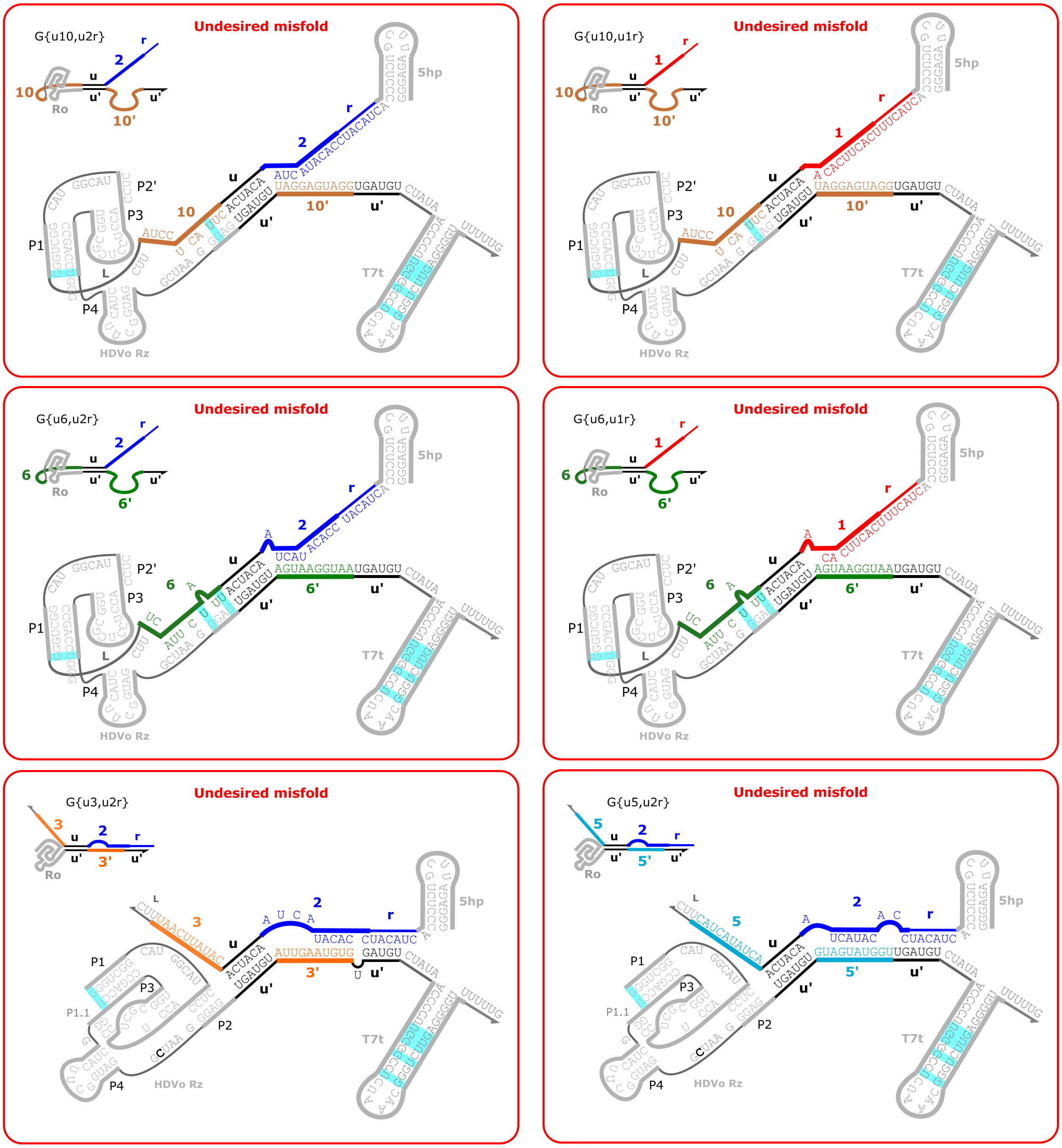

**Figure S31:** Possible misfolded structures of gates with poor performance that had repeated input-output toehold sequences. These gates were all characterized in Figure 2 of the main text. Cyan shaded base pairs represent G-U wobble base pairs.

### Supporting Note 6: Individual plots of reporter kinetics

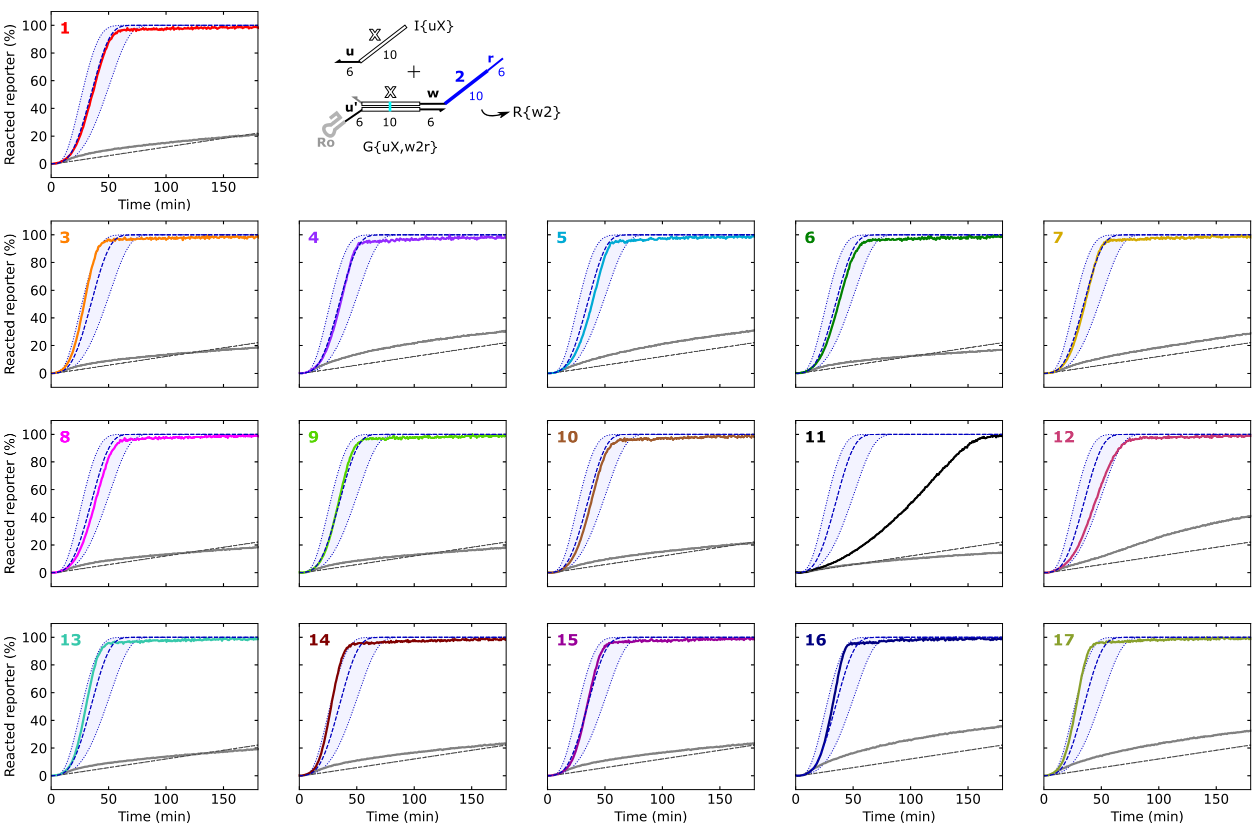

**Figure S32:** Individual plots of reporter kinetics from Figure 2A. In the plots, numbers in the upper righthand corner indicate the identity of the *X* domain. Colored lines indicate gates transcribed with their designed input (I{uX}), and gray lines indicate gates transcribed with a scrambled input (Io). The G{u1,w2r} sample was used to calibrate the transcription rate constant ($k_{txn}$) for these experiments.

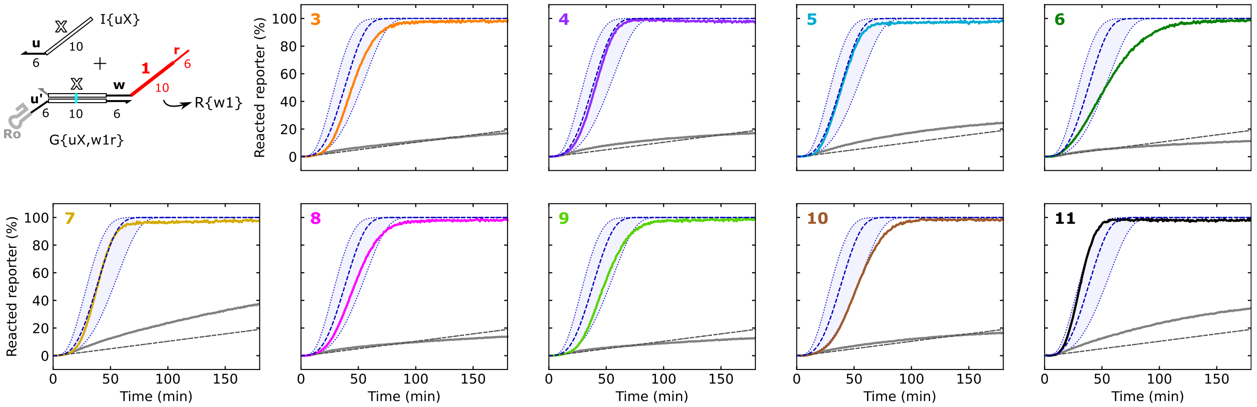

**Figure S33:** Individual plots of reporter kinetics from Figure 2C. In the plots, numbers in the upper righthand corner indicate the identity of the *X* domain. Colored lines indicate gates transcribed with their designed input (I{uX}), and gray lines indicate gates transcribed with a scrambled input (Io).

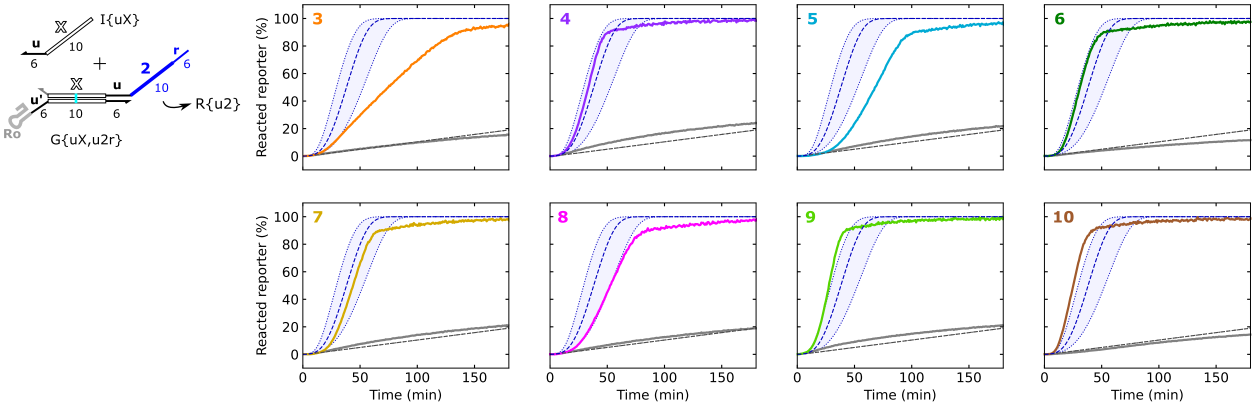

**Figure S34:** Individual plots of reporter kinetics from Figure 2D, left. In the plots, numbers in the upper righthand corner indicate the identity of the *X* domain. Colored lines indicate gates transcribed with their designed input (I{uX}), and gray lines indicate gates transcribed with a scrambled input (Io).

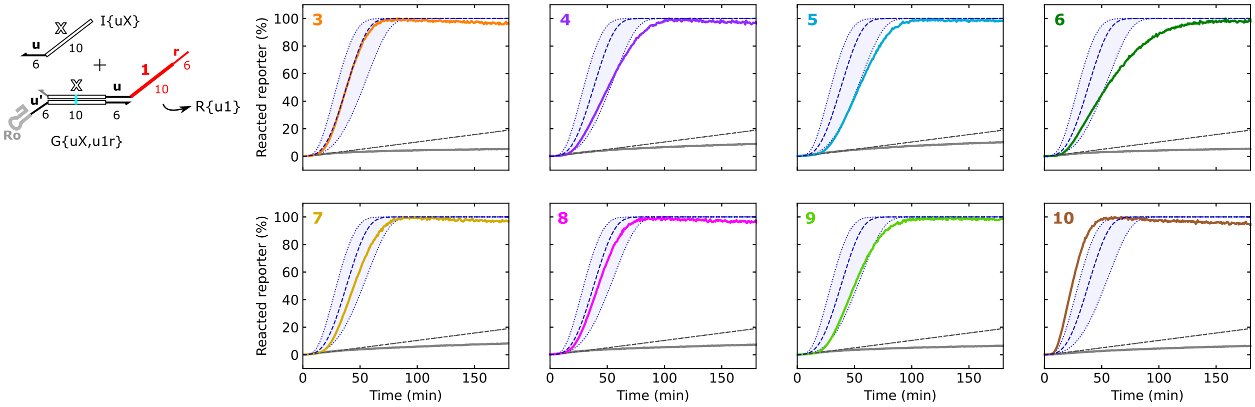

**Figure S35:** Individual plots of reporter kinetics from Figure 2D, right. In the plots, numbers in the upper righthand corner indicate the identity of the *X* domain. Colored lines indicate gates transcribed with their designed input (I{uX}), and gray lines indicate gates transcribed with a scrambled input (Io).

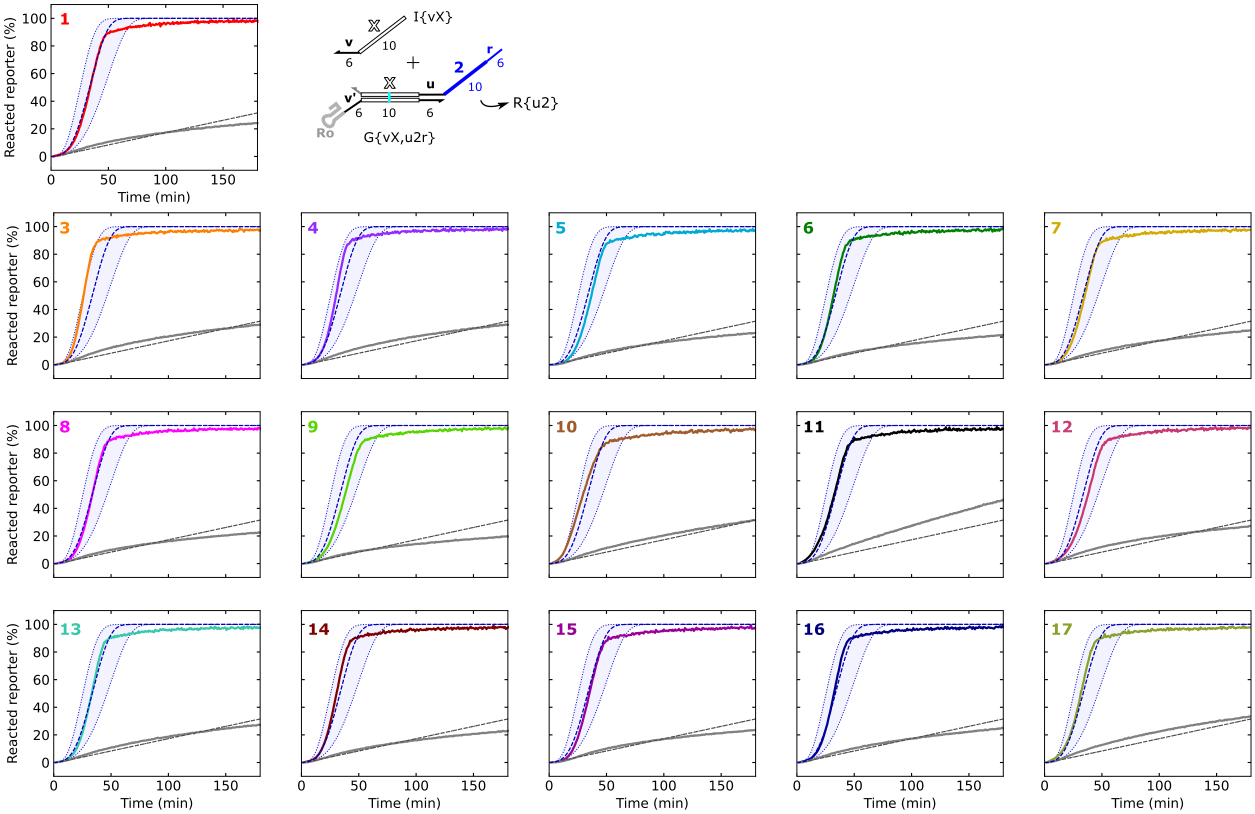

**Figure S36:** Individual plots of reporter kinetics from Figure 3E. In the plots, numbers in the upper righthand corner indicate the identity of the *X* domain. Colored lines indicate gates transcribed with their designed input (I{uX}), and gray lines indicate gates transcribed with a scrambled input (Io). The G{v1,u2r} sample was used to calibrate the transcription rate constant ($k_{txn}$) for these experiments.

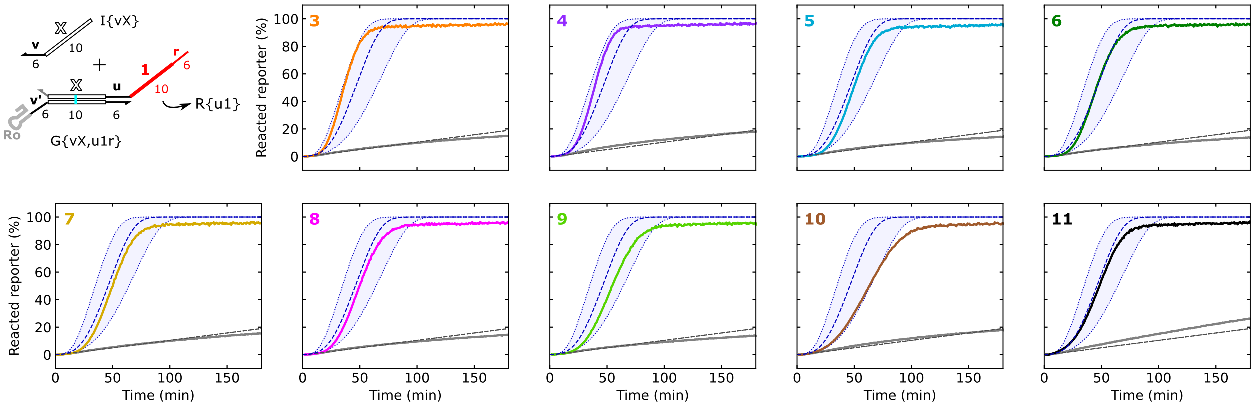

**Figure S37:** Individual plots of reporter kinetics from Figure S5. In the plots, numbers in the upper righthand corner indicate the identity of the *X* domain. Colored lines indicate gates transcribed with their designed input (I{uX}), and gray lines indicate gates transcribed with a scrambled input (Io).

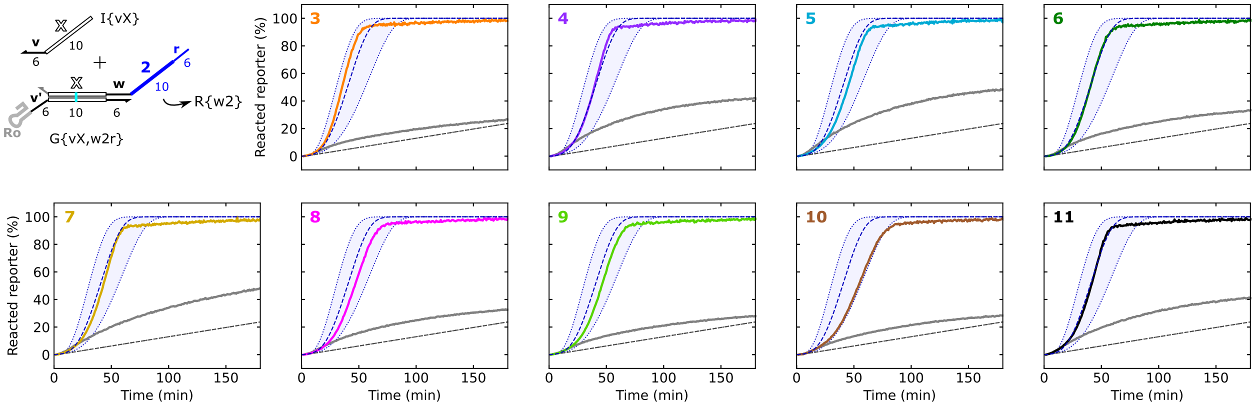

**Figure S38:** Individual plots of reporter kinetics from Figure S6A. In the plots, numbers in the upper righthand corner indicate the identity of the *X* domain. Colored lines indicate gates transcribed with their designed input (I{uX}), and gray lines indicate gates transcribed with a scrambled input (Io).

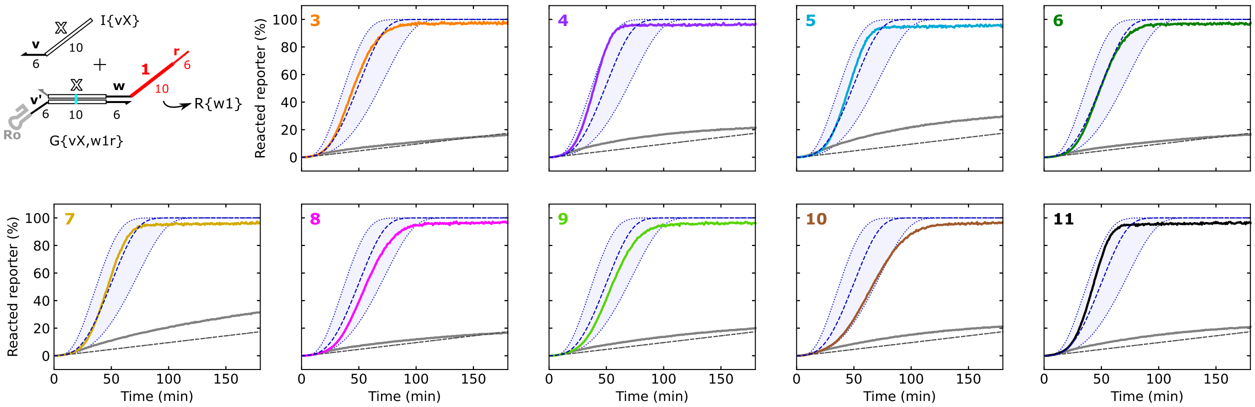

**Figure S39:** Individual plots of reporter kinetics from Figure S6B. In the plots, numbers in the upper righthand corner indicate the identity of the *X* domain. Colored lines indicate gates transcribed with their designed input (I{uX}), and gray lines indicate gates transcribed with a scrambled input (Io).

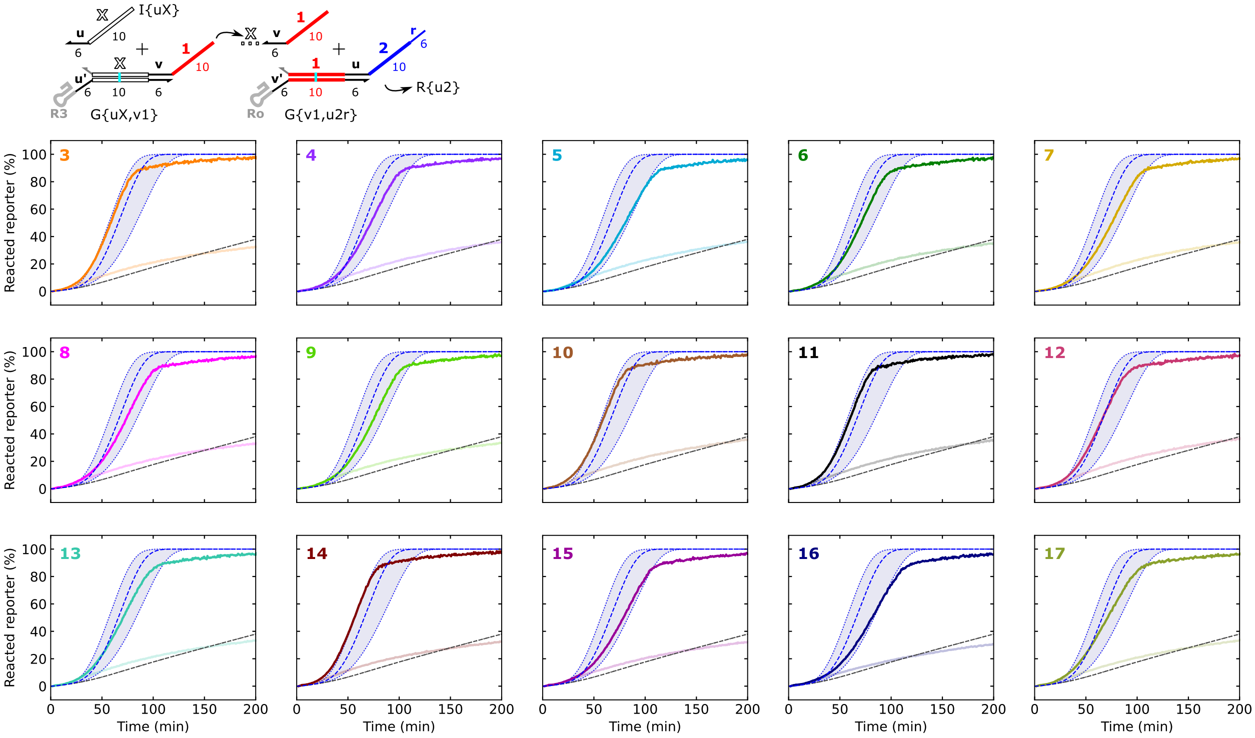

**Figure S40** Individual plots of reporter kinetics from Figure 5C. In the plots, numbers in the upper righthand corner indicate the identity of the *X* domain. Colored lines indicate gates transcribed with their designed input (I{uX}), and gray lines indicate gates transcribed with a scrambled input (Io).

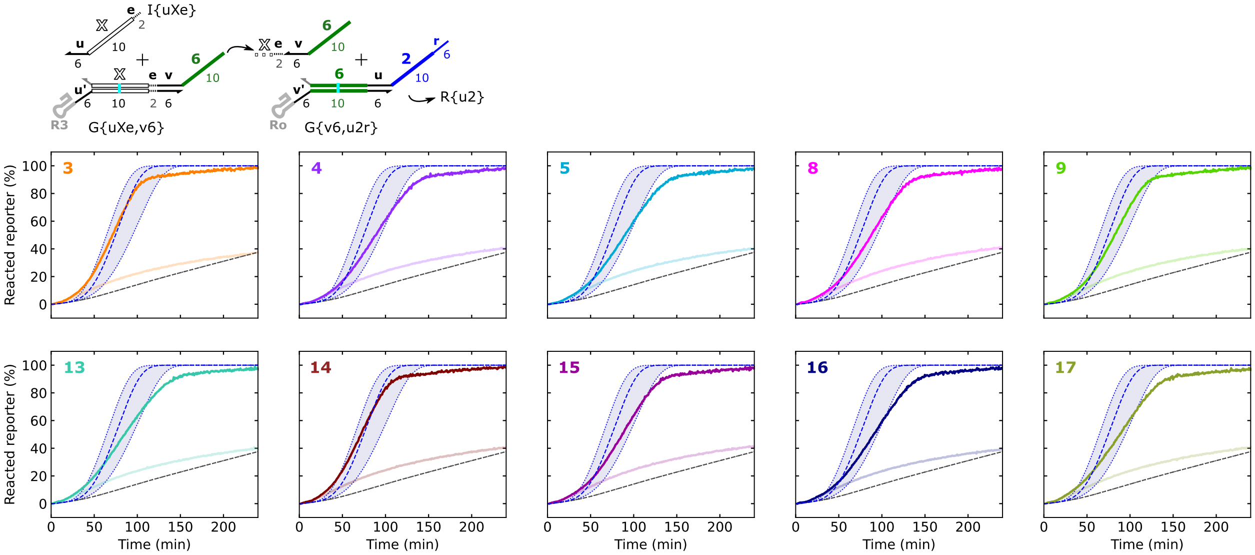

**Figure S41:** Individual plots of reporter kinetics from Figure 5F, top. In the plots, numbers in the upper righthand corner indicate the identity of the *X* domain. Colored lines indicate gates transcribed with their designed input (I{uX}), and gray lines indicate gates transcribed with a scrambled input (Io).

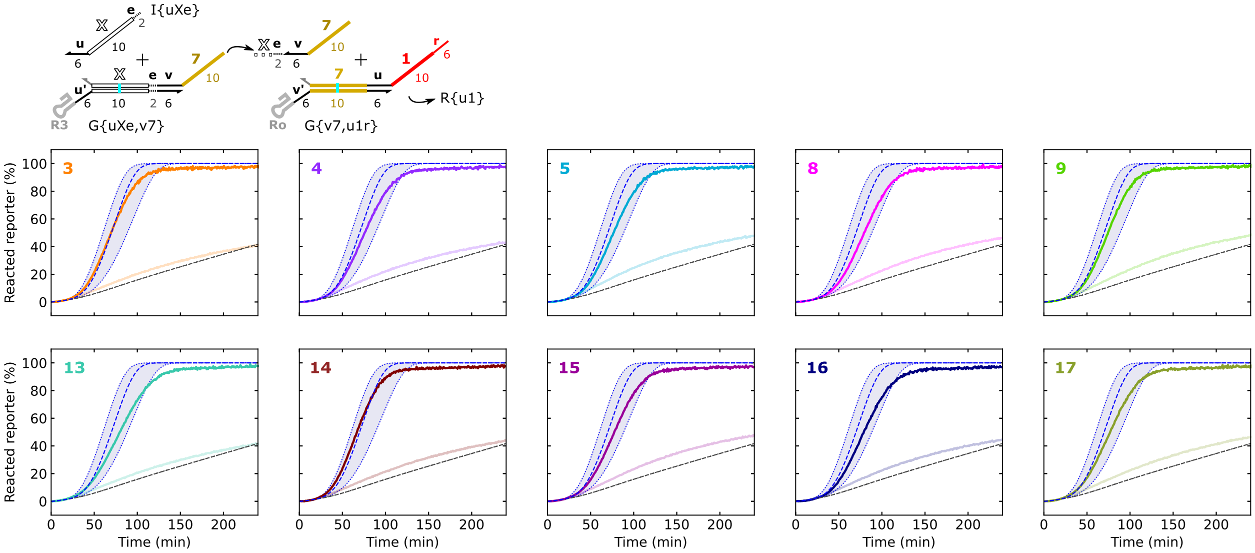

**Figure S42:** Individual plots of reporter kinetics from Figure 5F, bottom. In the plots, numbers in the upper righthand corner indicate the identity of the *X* domain. Colored lines indicate gates transcribed with their designed input (I{uX}), and gray lines indicate gates transcribed with a scrambled input (Io).
