## Supplementary material for "Design approaches to expand the toolkit for building cotranscriptionally encoded RNA strand displacement circuits": Simulation Scripts: Figure2A_ctRSD_toolkit.py

```
# -*- coding: utf-8 -*-
"""
Created on Wed Jul 27 14:05:50 2022

@author: tnm12
"""

import numpy as np
import matplotlib.pyplot as plt
from matplotlib.ticker import MaxNLocator
import openpyxl as opxl
import xlrd
xlrd.xlsx.ensure_elementtree_imported(False, None)
xlrd.xlsx.Element_has_iter = True
import scipy.integrate as spi

import sys
sys.path.insert(1,'C:\\Users\\sws5\\OneDrive - National Institute of Standards and Technology (NIST)\\ctRSD simulator and documentation\\ctRSD-simulator\\ctRSD-simulator-2.0')
import ctRSD_simulator_200 as RSDs

'''
##############################################################################
Loading data
##############################################################################
'''

save_loc = 'C:\\Users\\sws5\\OneDrive - National Institute of Standards and Technology (NIST)\\Presentations and write ups\\Papers\\RSD technical note\\Data upload files\\'
save_name = 'Supporting File S3'

wb = xlrd.open_workbook(save_loc+save_name+'.xlsx')

ws = wb.sheet_by_name('Figure 2A (X=1-12)')

t = np.zeros(ws.nrows-3)
for r in range(3,ws.nrows):
    t[r-3] = ws.cell_value(r,0)
    
DDn = np.zeros([ws.nrows-3,int(ws.ncols/2)-1])

for c in range(1,int(ws.ncols/2)):
    for r in range(3,ws.nrows):
        DDn[r-3,c-1] = ws.cell_value(r,c)
        
ws2 = wb.sheet_by_name('Figure 2A (X=13-17)')

t2 = np.zeros(ws2.nrows-3)
for r in range(3,ws2.nrows):
    t2[r-3] = ws2.cell_value(r,0)
    
DDn2 = np.zeros([ws2.nrows-3,int(ws2.ncols/2)-1])

for c in range(1,int(ws2.ncols/2)):
    for r in range(3,ws2.nrows):
        DDn2[r-3,c-1] = ws2.cell_value(r,c)


'''
###############################################################################
PLOTTING DATA
###############################################################################
'''

clp = [0.5,0.5,0.5]

clp2 = [[1,0,0],
       [1,0.5,0],
       [147/255,42/255,255/255],
       [0/255,170/255,212/255],
       [0,0.5,0],
       [212/255,170/255,0],
       [1,0,1],
       [85/255,212/255,0],
       [160/255,90/255,44/255],
       [0,0,0],
       [200/255,55/255,113/255]]

clp3 = [[55/255,200/255,171/255],
       [0.5,0,0],
       [0.6,0,0.6],
       [0,0,0.5],
       [137/255,160/255,44/255]]


fs = 12 # font size
ts = 26 # time before T7 RNAP was added to reactions (X=1-12)

# individual fluorescence plots (X=1-12)
plt.figure(1)

p = [1,6,7,8,9,10,11,12,13,14,15]
for n in range(len(p)):
    plt.subplot(3,5,p[n])  
    plt.plot(t-ts,DDn[:,n]*100,color=clp,linewidth=2)
    plt.plot(t-ts,DDn[:,n+11]*100,color=clp2[n],linewidth=2)
    plt.xticks(fontsize=fs)
    plt.yticks(fontsize=fs)
    plt.ylim(-10,110)
    plt.xlim(0,180)
    ax1 = plt.gca()
    ax1.xaxis.set_tick_params(which='both',size=3,width=1,direction='in',top='on')
    #ax1.xaxis.set_major_locator(MaxNLocator(integer=True))
    ax1.yaxis.set_tick_params(which='both',size=3,width=1,direction='in',right='on')
    if n == 0:
        plt.xlabel('Time (min)',fontsize=fs)
    else:
        ax1.xaxis.set_ticklabels([])
    if n == 0 or n == 1 or n == 6:
        plt.ylabel('Reacted reporter (%)',fontsize=fs)
    else:
        ax1.yaxis.set_ticklabels([])


# individual fluorescence plots (X=13-17)
plt.figure(2)

ts2 = 14.7 # time before T7 RNAP was added to reactions (X=13-17)

p2 = [6,7,8,9,10]
for n in range(len(p2)):
    plt.subplot(3,5,p2[n])
    plt.plot(t2-ts2,DDn2[:,n]*100,color=clp,linewidth=2)
    plt.plot(t2-ts2,DDn2[:,n+6]*100,color=clp3[n],linewidth=2)
    plt.xticks(fontsize=fs)
    plt.yticks(fontsize=fs)
    plt.ylim(-10,110)
    plt.xlim(0,180)
    ax1 = plt.gca()
    ax1.xaxis.set_tick_params(which='both',size=3,width=1,direction='in',top='on')
    #ax1.xaxis.set_major_locator(MaxNLocator(integer=True))
    ax1.yaxis.set_tick_params(which='both',size=3,width=1,direction='in',right='on')
    plt.xlabel('Time (min)',fontsize=fs)
    if n == 0:
        plt.ylabel('Reacted reporter (%)',fontsize=fs)
    else:
        ax1.yaxis.set_ticklabels([])
        
        
# Overlaid fluorescence plots 
plt.figure(3)

plt.subplot(3,5,1)
for n in range(len(p)):
    plt.plot(t-ts,DDn[:,n]*100,color=clp2[n],linewidth=2,alpha=0.25)
    plt.plot(t-ts,DDn[:,n+11]*100,color=clp2[n],linewidth=2)
    plt.xticks(fontsize=fs)
    plt.yticks(fontsize=fs)
    plt.ylim(-10,110)
    plt.xlim(0,180)

plt.subplot(3,5,1)
for n in range(len(p2[1:])):
    plt.plot(t2-ts2,DDn2[:,n+1]*100,color=clp3[n+1],linewidth=2,alpha=0.25)
    plt.plot(t2-ts2,DDn2[:,n+6+1]*100,color=clp3[n+1],linewidth=2)
    plt.xticks(fontsize=fs)
    plt.yticks(fontsize=fs)
    plt.ylim(-10,110)
    plt.xlim(0,180)
    
ax1 = plt.gca()
ax1.xaxis.set_tick_params(which='both',size=3,width=1,direction='in',top='on')
#ax1.xaxis.set_major_locator(MaxNLocator(integer=True))
ax1.yaxis.set_tick_params(which='both',size=3,width=1,direction='in',right='on')
plt.xlabel('Time (min)',fontsize=fs)
plt.ylabel('Reacted reporter (%)',fontsize=fs)
plt.show()
        
'''
##############################################################################
SIMULATIONS 
##############################################################################
'''

t_sim = np.linspace(0,6,1001)*3600 # seconds

k_txn = 0.014

IN = [0,50]

REP_con = 500

krsd = [0.25e3/1e9,1e3/1e9,4e3/1e9] # 1/nM-s


REP2 = np.array([])
REP2_l = np.array([])
REP2_h = np.array([])

for x in range(len(krsd)):
        
    for n in range(len(IN)):
        

        model = RSDs.RSD_sim() # define the model instance
        
        model.global_rate_constants(krsd=krsd[x],ktxn=k_txn)
        
        # specify DNA txn templates and reporters and concentrations
        model.molecular_species('G{1,2}',DNA_con=25,krev=5/1e9)
        model.molecular_species('R{2}',DNA_con=REP_con)
       
        model.molecular_species('I{1}',DNA_con=IN[n])
        
        # simulate the model
        model.simulate(t_sim)
        
        if x == 0:
            # pull out the species from the model solution to plot
            if len(REP2_l) == 0:
                REP2_l = model.output_concentration('S{2}')
            else:
                REP2_l = np.append([REP2_l],[model.output_concentration('S{2}')],axis=0)
        elif x == 1:
            if len(REP2) == 0:
                REP2 = model.output_concentration('S{2}')
            else:
                REP2 = np.append([REP2],[model.output_concentration('S{2}')],axis=0)
        elif x == 2:
            if len(REP2_h) == 0:
                REP2_h = model.output_concentration('S{2}')
            else:
                REP2_h = np.append([REP2_h],[model.output_concentration('S{2}')],axis=0)
     
plt.figure(1)
i = 0     
for m in range(len(p)):
    plt.subplot(3,5,p[m])
    plt.plot(model.t/60,(REP2[i]/REP_con)*100,color=[0.3,0.3,0.3],linewidth=1,linestyle='--')
    plt.plot(model.t/60,(REP2[i+1]/REP_con)*100,color=[0,0,0.75],linewidth=1,linestyle='--')
    plt.plot(model.t/60,(REP2_l[i]/REP_con)*100,color=[0.3,0.3,0.3],linewidth=0.75,linestyle=':',zorder=0)
    plt.plot(model.t/60,(REP2_l[i+1]/REP_con)*100,color=[0,0,0.75],linewidth=.75,linestyle=':',zorder=0)
    plt.plot(model.t/60,(REP2_h[i]/REP_con)*100,color=[0.3,0.3,0.3],linewidth=0.75,linestyle=':',zorder=0)
    plt.plot(model.t/60,(REP2_h[i+1]/REP_con)*100,color=[0,0,0.75],linewidth=.75,linestyle=':',zorder=0)
    plt.fill_between(model.t/60,(REP2_h[i+1]/REP_con)*100,(REP2_l[i+1]/REP_con)*100,color=[0,0,1],alpha=0.05,zorder=0)

plt.figure(2)
i = 0     
for m in range(len(p2)):
    plt.subplot(3,5,p2[m])
    plt.plot(model.t/60,(REP2[i]/REP_con)*100,color=[0.3,0.3,0.3],linewidth=1,linestyle='--')
    plt.plot(model.t/60,(REP2[i+1]/REP_con)*100,color=[0,0,0.75],linewidth=1,linestyle='--')
    plt.plot(model.t/60,(REP2_l[i]/REP_con)*100,color=[0.3,0.3,0.3],linewidth=0.75,linestyle=':',zorder=0)
    plt.plot(model.t/60,(REP2_l[i+1]/REP_con)*100,color=[0,0,0.75],linewidth=.75,linestyle=':',zorder=0)
    plt.plot(model.t/60,(REP2_h[i]/REP_con)*100,color=[0.3,0.3,0.3],linewidth=0.75,linestyle=':',zorder=0)
    plt.plot(model.t/60,(REP2_h[i+1]/REP_con)*100,color=[0,0,0.75],linewidth=.75,linestyle=':',zorder=0)
    plt.fill_between(model.t/60,(REP2_h[i+1]/REP_con)*100,(REP2_l[i+1]/REP_con)*100,color=[0,0,1],alpha=0.05,zorder=0)


plt.figure(3)
i = 0     
plt.subplot(3,5,1)
plt.plot(model.t/60,(REP2[i]/REP_con)*100,color=[0.3,0.3,0.3],linewidth=1,linestyle='--')
plt.plot(model.t/60,(REP2[i+1]/REP_con)*100,color=[0,0,0.75],linewidth=1,linestyle='--')
plt.plot(model.t/60,(REP2_l[i]/REP_con)*100,color=[0.3,0.3,0.3],linewidth=0.75,linestyle=':',zorder=0)
plt.plot(model.t/60,(REP2_l[i+1]/REP_con)*100,color=[0,0,0.75],linewidth=.75,linestyle=':',zorder=0)
plt.plot(model.t/60,(REP2_h[i]/REP_con)*100,color=[0.3,0.3,0.3],linewidth=0.75,linestyle=':',zorder=0)
plt.plot(model.t/60,(REP2_h[i+1]/REP_con)*100,color=[0,0,0.75],linewidth=.75,linestyle=':',zorder=0)
plt.fill_between(model.t/60,(REP2_h[i+1]/REP_con)*100,(REP2_l[i+1]/REP_con)*100,color=[0,0,1],alpha=0.1,zorder=0)
```
