## Supplementary material for "Design approaches to expand the toolkit for building cotranscriptionally encoded RNA strand displacement circuits": Simulation Scripts: Figure4Dbottom_ctRSD_toolkit.py

```
# -*- coding: utf-8 -*-
"""
Created on Wed Jul 27 16:34:07 2022

@author: tnm12
"""

import numpy as np
import matplotlib.pyplot as plt
from matplotlib.ticker import MaxNLocator
import openpyxl as opxl
import xlrd
xlrd.xlsx.ensure_elementtree_imported(False, None)
xlrd.xlsx.Element_has_iter = True
import scipy.integrate as spi

import sys
sys.path.insert(1,'C:\\Users\\sws5\\OneDrive - National Institute of Standards and Technology (NIST)\\ctRSD simulator and documentation\\ctRSD-simulator\\ctRSD-simulator-2.0')
import ctRSD_simulator_200 as RSDs

'''
##############################################################################
Loading data
##############################################################################
'''

save_loc = 'C:\\Users\\sws5\\OneDrive - National Institute of Standards and Technology (NIST)\\Presentations and write ups\\Papers\\RSD technical note\\Data upload files\\'
save_name = 'Supporting File S3'

wb = xlrd.open_workbook(save_loc+save_name+'.xlsx')

ws = wb.sheet_by_name('Figure 4D, bottom')

t = np.zeros(ws.nrows-3)
for r in range(3,ws.nrows):
    t[r-3] = ws.cell_value(r,0)
    
DDn = np.zeros([ws.nrows-3,int(ws.ncols/2)-1])

for c in range(1,int(ws.ncols/2)):
    for r in range(3,ws.nrows):
        DDn[r-3,c-1] = ws.cell_value(r,c)

'''
###############################################################################
PLOTTING DATA
###############################################################################
'''

clp2 = [[0,0,1],
       [0.25,0.25,0.25],
       [0.5,0.5,0.5],
       [158/255,100/255,64/255],
       [85/255,212/255,0],
       [0,0,0.5]]

fs = 12
ts = 0

# fluorescence plots
plt.figure(1)
for n in range(int(len(DDn[1,:])/2)):
    plt.subplot(3,5,1)
    plt.plot(t-ts,DDn[:,n+1]*100,color=clp2[n],alpha=0.25,linewidth=2)
    plt.plot(t-ts,DDn[:,n+4+1]*100,color=clp2[n],linewidth=2)
    plt.xticks(fontsize=fs)
    plt.yticks(fontsize=fs)
    plt.ylim(-10,110)
    plt.xlim(0,240)
    ax1 = plt.gca()
    ax1.xaxis.set_tick_params(which='both',size=3,width=1,direction='in',top='on')
    #ax1.xaxis.set_major_locator(MaxNLocator(integer=True))
    ax1.yaxis.set_tick_params(which='both',size=3,width=1,direction='in',right='on')
    plt.xlabel('Time (min)',fontsize=fs)
    plt.ylabel('Reacted reporter (%)',fontsize=fs)


'''
##############################################################################
Simulations 
##############################################################################
'''

t_sim = np.linspace(0,6,1001)*3600 # seconds

k_txn = 0.010

IN = [0,25]
REP_con=500

# ksd is 1.8e3 and 6e2 to recapitulate the other two curves (less than a factor of 2 difference)
krsd = [0.2e3/1e9,1e3/1e9,5e3/1e9] # 1/nM-s

REP2 = np.array([])
REP2_l = np.array([])
REP2_h = np.array([])

for x in range(len(krsd)):
        
    for n in range(len(IN)):
        

        model = RSDs.RSD_sim() # define the model instance
        
        model.global_rate_constants(ktxn=k_txn)
        
        # specify DNA txn templates and reporters and concentrations
        model.molecular_species('G{1,2}',DNA_con=25)
        model.molecular_species('G{5,1}',DNA_con=25,krsd=krsd[x])
        model.molecular_species('R{2}',DNA_con=500)
       
        model.molecular_species('I{5}',DNA_con=IN[n])
        
        # simulate the model
        model.simulate(t_sim)
        
        if x == 0:
            # pull out the species from the model solution to plot
            if len(REP2_l) == 0:
                REP2_l = model.output_concentration('S{2}')
            else:
                REP2_l = np.append([REP2_l],[model.output_concentration('S{2}')],axis=0)
        elif x == 1:
            if len(REP2) == 0:
                REP2 = model.output_concentration('S{2}')
            else:
                REP2 = np.append([REP2],[model.output_concentration('S{2}')],axis=0)
        elif x == 2:
            if len(REP2_h) == 0:
                REP2_h = model.output_concentration('S{2}')
            else:
                REP2_h = np.append([REP2_h],[model.output_concentration('S{2}')],axis=0)
                
i = 0     
plt.figure(1)
plt.subplot(3,5,1)
plt.plot(model.t/60,(REP2[i]/REP_con)*100,color=[0.3,0.3,0.3],linewidth=1,linestyle='--')
plt.plot(model.t/60,(REP2[i+1]/REP_con)*100,color=[0,0,1],linewidth=1,linestyle='--')
plt.plot(model.t/60,(REP2_l[i]/REP_con)*100,color=[0.3,0.3,0.3],linewidth=0.75,linestyle=':',zorder=0)
plt.plot(model.t/60,(REP2_l[i+1]/REP_con)*100,color=[0,0,1],linewidth=.75,linestyle=':',zorder=0)
plt.plot(model.t/60,(REP2_h[i]/REP_con)*100,color=[0.3,0.3,0.3],linewidth=0.75,linestyle=':',zorder=0)
plt.plot(model.t/60,(REP2_h[i+1]/REP_con)*100,color=[0,0,1],linewidth=.75,linestyle=':',zorder=0)
plt.fill_between(model.t/60,(REP2_h[i+1]/REP_con)*100,(REP2_l[i+1]/REP_con)*100,color=[0,0,0.5],alpha=0.1,zorder=0)
```
