## Supplementary material for "Design approaches to expand the toolkit for building cotranscriptionally encoded RNA strand displacement circuits": Simulation Scripts: Figure6C_ctRSD_toolkit.py

```
import numpy as np
import matplotlib.pyplot as plt
from matplotlib.ticker import MaxNLocator
import openpyxl as opxl
import xlrd
import scipy.integrate as spi

import sys
sys.path.insert(1,'C:\\Users\\sws5\\OneDrive - National Institute of Standards and Technology (NIST)\\ctRSD simulator and documentation\\ctRSD-simulator\\ctRSD-simulator-2.0')
import ctRSD_simulator_200 as RSDs

'''
##############################################################################
Loading data
##############################################################################
'''

save_loc = 'C:\\Users\\sws5\\OneDrive - National Institute of Standards and Technology (NIST)\\Presentations and write ups\\Papers\\RSD technical note\\Data upload files\\'
save_name = 'Supporting File S3'

wb = xlrd.open_workbook(save_loc+save_name+'.xlsx')

ws = wb.sheet_by_name('Figure 6C, 1')

t = np.zeros(ws.nrows-3)
for r in range(3,ws.nrows):
    t[r-3] = ws.cell_value(r,0)
    
DDn = np.zeros([ws.nrows-3,int(ws.ncols/2)-1])

for c in range(1,int(ws.ncols/2)):
    for r in range(3,ws.nrows):
        DDn[r-3,c-1] = ws.cell_value(r,c)
        
ws2 = wb.sheet_by_name('Figure 6C, 3')

t2 = np.zeros(ws2.nrows-3)
for r in range(3,ws2.nrows):
    t2[r-3] = ws2.cell_value(r,0)
    
DDn2 = np.zeros([ws2.nrows-3,int(ws2.ncols/2)-1])

for c in range(1,int(ws2.ncols/2)):
    for r in range(3,ws2.nrows):
        DDn2[r-3,c-1] = ws2.cell_value(r,c)
        
ws3 = wb.sheet_by_name('Figure 6C, 5')

t3 = np.zeros(ws3.nrows-3)
for r in range(3,ws3.nrows):
    t3[r-3] = ws3.cell_value(r,0)
    
DDn3 = np.zeros([ws3.nrows-3,int(ws3.ncols/2)-1])

for c in range(1,int(ws3.ncols/2)):
    for r in range(3,ws3.nrows):
        DDn3[r-3,c-1] = ws3.cell_value(r,c)
        
ws4 = wb.sheet_by_name('Figure 6C, 7')

t4 = np.zeros(ws4.nrows-3)
for r in range(3,ws4.nrows):
    t4[r-3] = ws4.cell_value(r,0)
    
DDn4 = np.zeros([ws4.nrows-3,int(ws4.ncols/2)-1])

for c in range(1,int(ws4.ncols/2)):
    for r in range(3,ws4.nrows):
        DDn4[r-3,c-1] = ws4.cell_value(r,c)


'''
###############################################################################
PLOTTING DATA
###############################################################################
'''

clp1 = [[1,0.5,0.5],
       [1,0,0],
       [0.5,0,0]]

clp2 = [[1,0.8,0],
       [1,0.6,0],
       [1,0.4,0]]

clp3 = [[0,0.8,0.8],
       [0,0.6,0.6],
       [0,0.4,0.4]]

clp4 = [[0.9,0.8,0],
       [0.7,0.6,0],
       [0.5,0.4,0]]

fs = 12
# time shifts before T7 RNAP was added to samples
ts1 = 13
ts2 = 29
ts3 = 37
ts4 = 0

# fluorescence plots
plt.figure(1)

# G{u1,w2r}
plt.subplot(3,5,1)
plt.plot(t-ts1,DDn[:,0]*100,color=clp1[0],alpha=0.25,linewidth=2)
plt.plot(t-ts1,DDn[:,1]*100,color=clp1[1],alpha=0.25,linewidth=2)
plt.plot(t-ts1,DDn[:,2]*100,color=clp1[2],alpha=0.25,linewidth=2)
plt.plot(t-ts1,DDn[:,3]*100,color=clp1[0],linewidth=2)
plt.plot(t-ts1,DDn[:,4]*100,color=clp1[1],linewidth=2)
plt.plot(t-ts1,DDn[:,5]*100,color=clp1[2],linewidth=2)
plt.xticks(fontsize=fs)
plt.yticks(fontsize=fs)
plt.ylim(-10,110)
plt.xlim(0,150)
ax1 = plt.gca()
ax1.xaxis.set_tick_params(which='both',size=3,width=1,direction='in',top='on')
#ax1.xaxis.set_major_locator(MaxNLocator(integer=True))
ax1.yaxis.set_tick_params(which='both',size=3,width=1,direction='in',right='on')
ax1.xaxis.set_ticklabels([])
plt.ylabel('Reacted reporter (%)',fontsize=fs)

# inverted G{u1,w2r}
plt.subplot(3,5,6)
plt.plot(t-ts1,DDn[:,6]*100,color=clp1[0],alpha=0.25,linewidth=2)
plt.plot(t-ts1,DDn[:,7]*100,color=clp1[1],alpha=0.25,linewidth=2)
plt.plot(t-ts1,DDn[:,8]*100,color=clp1[2],alpha=0.25,linewidth=2)
plt.plot(t-ts1,DDn[:,9]*100,color=clp1[0],linewidth=2)
plt.plot(t-ts1,DDn[:,10]*100,color=clp1[1],linewidth=2)
plt.plot(t-ts1,DDn[:,11]*100,color=clp1[2],linewidth=2)
plt.xticks(fontsize=fs)
plt.yticks(fontsize=fs)
plt.ylim(-10,110)
plt.xlim(0,150)
ax1 = plt.gca()
ax1.xaxis.set_tick_params(which='both',size=3,width=1,direction='in',top='on')
#ax1.xaxis.set_major_locator(MaxNLocator(integer=True))
ax1.yaxis.set_tick_params(which='both',size=3,width=1,direction='in',right='on')
plt.ylabel('Reacted reporter (%)',fontsize=fs)
plt.xlabel('Time (min)',fontsize=fs)

# G{u3,w2r}
plt.subplot(3,5,2)
plt.plot(t2-ts2,DDn2[:,0]*100,color=clp2[0],alpha=0.25,linewidth=2)
plt.plot(t2-ts2,DDn2[:,1]*100,color=clp2[1],alpha=0.25,linewidth=2)
plt.plot(t2-ts2,DDn2[:,2]*100,color=clp2[2],alpha=0.25,linewidth=2)
plt.plot(t2-ts2,DDn2[:,3]*100,color=clp2[0],linewidth=2)
plt.plot(t2-ts2,DDn2[:,4]*100,color=clp2[1],linewidth=2)
plt.plot(t2-ts2,DDn2[:,5]*100,color=clp2[2],linewidth=2)
plt.xticks(fontsize=fs)
plt.yticks(fontsize=fs)
plt.ylim(-10,110)
plt.xlim(0,150)
ax1 = plt.gca()
ax1.xaxis.set_tick_params(which='both',size=3,width=1,direction='in',top='on')
#ax1.xaxis.set_major_locator(MaxNLocator(integer=True))
ax1.yaxis.set_tick_params(which='both',size=3,width=1,direction='in',right='on')
ax1.xaxis.set_ticklabels([])
ax1.yaxis.set_ticklabels([])

# inverted G{u3,w2r}
plt.subplot(3,5,7)
plt.plot(t2-ts2,DDn2[:,6]*100,color=clp2[0],alpha=0.25,linewidth=2)
plt.plot(t2-ts2,DDn2[:,7]*100,color=clp2[1],alpha=0.25,linewidth=2)
plt.plot(t2-ts2,DDn2[:,8]*100,color=clp2[2],alpha=0.25,linewidth=2)
plt.plot(t2-ts2,DDn2[:,9]*100,color=clp2[0],linewidth=2)
plt.plot(t2-ts2,DDn2[:,10]*100,color=clp2[1],linewidth=2)
plt.plot(t2-ts2,DDn2[:,11]*100,color=clp2[2],linewidth=2)
plt.xticks(fontsize=fs)
plt.yticks(fontsize=fs)
plt.ylim(-10,110)
plt.xlim(0,150)
ax1 = plt.gca()
ax1.xaxis.set_tick_params(which='both',size=3,width=1,direction='in',top='on')
#ax1.xaxis.set_major_locator(MaxNLocator(integer=True))
ax1.yaxis.set_tick_params(which='both',size=3,width=1,direction='in',right='on')
ax1.yaxis.set_ticklabels([])
plt.xlabel('Time (min)',fontsize=fs)

# G{u5,w2r}
plt.subplot(3,5,3)
plt.plot(t3-ts3,DDn3[:,0]*100,color=clp3[0],alpha=0.25,linewidth=2)
plt.plot(t3-ts3,DDn3[:,1]*100,color=clp3[1],alpha=0.25,linewidth=2)
plt.plot(t3-ts3,DDn3[:,2]*100,color=clp3[2],alpha=0.25,linewidth=2)
plt.plot(t3-ts3,DDn3[:,3]*100,color=clp3[0],linewidth=2)
plt.plot(t3-ts3,DDn3[:,4]*100,color=clp3[1],linewidth=2)
plt.plot(t3-ts3,DDn3[:,5]*100,color=clp3[2],linewidth=2)
plt.xticks(fontsize=fs)
plt.yticks(fontsize=fs)
plt.ylim(-10,110)
plt.xlim(0,150)
ax1 = plt.gca()
ax1.xaxis.set_tick_params(which='both',size=3,width=1,direction='in',top='on')
#ax1.xaxis.set_major_locator(MaxNLocator(integer=True))
ax1.yaxis.set_tick_params(which='both',size=3,width=1,direction='in',right='on')
ax1.xaxis.set_ticklabels([])
ax1.yaxis.set_ticklabels([])

# inverted G{u5,w2r}
plt.subplot(3,5,8)
plt.plot(t3-ts3,DDn3[:,6]*100,color=clp3[0],alpha=0.25,linewidth=2)
plt.plot(t3-ts3,DDn3[:,7]*100,color=clp3[1],alpha=0.25,linewidth=2)
plt.plot(t3-ts3,DDn3[:,8]*100,color=clp3[2],alpha=0.25,linewidth=2)
plt.plot(t3-ts3,DDn3[:,9]*100,color=clp3[0],linewidth=2)
plt.plot(t3-ts3,DDn3[:,10]*100,color=clp3[1],linewidth=2)
plt.plot(t3-ts3,DDn3[:,11]*100,color=clp3[2],linewidth=2)
plt.xticks(fontsize=fs)
plt.yticks(fontsize=fs)
plt.ylim(-10,110)
plt.xlim(0,150)
ax1 = plt.gca()
ax1.xaxis.set_tick_params(which='both',size=3,width=1,direction='in',top='on')
#ax1.xaxis.set_major_locator(MaxNLocator(integer=True))
ax1.yaxis.set_tick_params(which='both',size=3,width=1,direction='in',right='on')
ax1.yaxis.set_ticklabels([])
plt.xlabel('Time (min)',fontsize=fs)

# G{u7,w2r}
plt.subplot(3,5,4)
plt.plot(t4-ts4,DDn4[:,0]*100,color=clp4[0],alpha=0.25,linewidth=2)
plt.plot(t4-ts4,DDn4[:,1]*100,color=clp4[1],alpha=0.25,linewidth=2)
plt.plot(t4-ts4,DDn4[:,2]*100,color=clp4[2],alpha=0.25,linewidth=2)
plt.plot(t4-ts4,DDn4[:,3]*100,color=clp4[0],linewidth=2)
plt.plot(t4-ts4,DDn4[:,4]*100,color=clp4[1],linewidth=2)
plt.plot(t4-ts4,DDn4[:,5]*100,color=clp4[2],linewidth=2)
plt.xticks(fontsize=fs)
plt.yticks(fontsize=fs)
plt.ylim(-10,110)
plt.xlim(0,150)
ax1 = plt.gca()
ax1.xaxis.set_tick_params(which='both',size=3,width=1,direction='in',top='on')
#ax1.xaxis.set_major_locator(MaxNLocator(integer=True))
ax1.yaxis.set_tick_params(which='both',size=3,width=1,direction='in',right='on')
ax1.xaxis.set_ticklabels([])
ax1.yaxis.set_ticklabels([])

# in verted G{u7,w2r}
plt.subplot(3,5,9)
plt.plot(t4-ts4,DDn4[:,6]*100,color=clp4[0],alpha=0.25,linewidth=2)
plt.plot(t4-ts4,DDn4[:,7]*100,color=clp4[1],alpha=0.25,linewidth=2)
plt.plot(t4-ts4,DDn4[:,8]*100,color=clp4[2],alpha=0.25,linewidth=2)
plt.plot(t4-ts4,DDn4[:,9]*100,color=clp4[0],linewidth=2)
plt.plot(t4-ts4,DDn4[:,10]*100,color=clp4[1],linewidth=2)
plt.plot(t4-ts4,DDn4[:,11]*100,color=clp4[2],linewidth=2)
plt.xticks(fontsize=fs)
plt.yticks(fontsize=fs)
plt.ylim(-10,110)
plt.xlim(0,150)
ax1 = plt.gca()
ax1.xaxis.set_tick_params(which='both',size=3,width=1,direction='in',top='on')
#ax1.xaxis.set_major_locator(MaxNLocator(integer=True))
ax1.yaxis.set_tick_params(which='both',size=3,width=1,direction='in',right='on')
ax1.yaxis.set_ticklabels([])
plt.xlabel('Time (min)',fontsize=fs)
```
