## Supplementary material for "Design approaches to expand the toolkit for building cotranscriptionally encoded RNA strand displacement circuits": Simulation Scripts: Figure6G_ctRSD_toolkit.py

```
# -*- coding: utf-8 -*-
"""
Created on Thu Jul 28 10:30:03 2022

@author: tnm12
"""

import numpy as np
import matplotlib.pyplot as plt
from matplotlib.ticker import MaxNLocator
import openpyxl as opxl
import xlrd
xlrd.xlsx.ensure_elementtree_imported(False, None)
xlrd.xlsx.Element_has_iter = True
import scipy.integrate as spi

import sys
sys.path.insert(1,'C:\\Users\\sws5\\OneDrive - National Institute of Standards and Technology (NIST)\\ctRSD simulator and documentation\\ctRSD-simulator\\ctRSD-simulator-2.0')
import ctRSD_simulator_200 as RSDs

'''
##############################################################################
Loading data
##############################################################################
'''

save_loc = 'C:\\Users\\sws5\\OneDrive - National Institute of Standards and Technology (NIST)\\Presentations and write ups\\Papers\\RSD technical note\\Data upload files\\'
save_name = 'Supporting File S3'

wb = xlrd.open_workbook(save_loc+save_name+'.xlsx')

ws = wb.sheet_by_name('Figure 6G')

t = np.zeros(ws.nrows-3)
for r in range(3,ws.nrows):
    t[r-3] = ws.cell_value(r,0)
    
DDn = np.zeros([ws.nrows-3,int(ws.ncols/2)-1])

for c in range(1,int(ws.ncols/2)):
    for r in range(3,ws.nrows):
        DDn[r-3,c-1] = ws.cell_value(r,c)

'''
###############################################################################
PLOTTING DATA
###############################################################################
'''

clp2 = [[0.4,0.4,0.4],
       [0,0,0.5],
       [0,0,1],
       [0,0.5,1],
       [212/255,170/255,0],
       [1,0,1],
       [85/255,212/255,0],
       [160/255,90/255,44/255],
       [0,0,0]]

fs = 12
ts = 0

# fluorescence plots
plt.figure(1)
for n in range(4):
    # without 3hp 
    plt.subplot(3,5,1)
    plt.plot(t-ts,DDn[:,n]*100,color=clp2[n],linewidth=2)
    plt.xticks(fontsize=fs)
    plt.yticks(fontsize=fs)
    plt.ylim(-10,110)
    plt.xlim(0,180)
    ax1 = plt.gca()
    ax1.xaxis.set_tick_params(which='both',size=3,width=1,direction='in',top='on')
    #ax1.xaxis.set_major_locator(MaxNLocator(integer=True))
    ax1.yaxis.set_tick_params(which='both',size=3,width=1,direction='in',right='on')
    plt.xlabel('Time (min)',fontsize=fs)
    plt.ylabel('Reacted reporter (%)',fontsize=fs)
    
    # with 3hp
    plt.subplot(3,5,2)
    plt.plot(t-ts,DDn[:,n+4]*100,color=clp2[n],linewidth=2)
    plt.xticks(fontsize=fs)
    plt.yticks(fontsize=fs)
    plt.ylim(-10,110)
    plt.xlim(0,180)
    ax1 = plt.gca()
    ax1.xaxis.set_tick_params(which='both',size=3,width=1,direction='in',top='on')
    #ax1.xaxis.set_major_locator(MaxNLocator(integer=True))
    ax1.yaxis.set_tick_params(which='both',size=3,width=1,direction='in',right='on')
    plt.xlabel('Time (min)',fontsize=fs)
    plt.ylabel('Reacted reporter (%)',fontsize=fs)
    
    # individual comparisons of w/wo the 3hp (3hp is with alpha = 0.5)
    plt.subplot(3,5,11+n)
    plt.plot(t-ts,DDn[:,n]*100,color=clp2[n],linewidth=2)
    plt.plot(t-ts,DDn[:,n+4]*100,color=clp2[n],linewidth=2,alpha=0.5)
    plt.xticks(fontsize=fs)
    plt.yticks(fontsize=fs)
    plt.ylim(-10,110)
    plt.xlim(0,180)
    ax1 = plt.gca()
    ax1.xaxis.set_tick_params(which='both',size=3,width=1,direction='in',top='on')
    #ax1.xaxis.set_major_locator(MaxNLocator(integer=True))
    ax1.yaxis.set_tick_params(which='both',size=3,width=1,direction='in',right='on')
    plt.xlabel('Time (min)',fontsize=fs)
    plt.ylabel('Reacted reporter (%)',fontsize=fs)
    

    
# Figure 6G plot
plt.figure(2)
plt.subplot(3,5,5)
plt.plot(t-ts,DDn[:,0]*100,color=[0,0,1],linewidth=2,alpha=0.25)
plt.plot(t-ts,DDn[:,0+4]*100,color=[16/255,136/255,255/255],linewidth=2,alpha=0.25)
plt.plot(t-ts,DDn[:,2]*100,color=[0,0,1],linewidth=2)
plt.plot(t-ts,DDn[:,2+4]*100,color=[16/255,136/255,255/255],linewidth=2)
plt.xticks(fontsize=fs)
plt.yticks(fontsize=fs)
plt.ylim(-10,110)
plt.xlim(0,180)
ax1 = plt.gca()
ax1.xaxis.set_tick_params(which='both',size=3,width=1,direction='in',top='on')
#ax1.xaxis.set_major_locator(MaxNLocator(integer=True))
ax1.yaxis.set_tick_params(which='both',size=3,width=1,direction='in',right='on')
plt.xlabel('Time (min)',fontsize=fs)
plt.ylabel('Reacted reporter (%)',fontsize=fs)
```
