## Supplementary material for "Design approaches to expand the toolkit for building cotranscriptionally encoded RNA strand displacement circuits": Simulation Scripts: FigureS2_ctRSD_toolkit.py

```
# -*- coding: utf-8 -*-
"""
Created on Thu Jul 28 08:49:11 2022

@author: tnm12
"""

import numpy as np
import matplotlib.pyplot as plt
from matplotlib.ticker import MaxNLocator
import openpyxl as opxl
import xlrd
xlrd.xlsx.ensure_elementtree_imported(False, None)
xlrd.xlsx.Element_has_iter = True
import scipy.integrate as spi

import sys
sys.path.insert(1,'C:\\Users\\sws5\\OneDrive - National Institute of Standards and Technology (NIST)\\ctRSD simulator and documentation\\ctRSD-simulator\\ctRSD-simulator-2.0')
import ctRSD_simulator_200 as RSDs

'''
##############################################################################
Loading data (Figure S2B)
##############################################################################
'''

save_loc = 'C:\\Users\\sws5\\OneDrive - National Institute of Standards and Technology (NIST)\\Presentations and write ups\\Papers\\RSD technical note\\Data upload files\\'
save_name = 'Supporting File S3'

wb = xlrd.open_workbook(save_loc+save_name+'.xlsx')

ws = wb.sheet_by_name('Figure S2B')

t = np.zeros(ws.nrows-3)
for r in range(3,ws.nrows):
    t[r-3] = ws.cell_value(r,0)
    
DDn = np.zeros([ws.nrows-3,int(ws.ncols/2)-1])

for c in range(1,int(ws.ncols/2)):
    for r in range(3,ws.nrows):
        DDn[r-3,c-1] = ws.cell_value(r,c)

'''
###############################################################################
PLOTTING DATA (Figure S2B)
###############################################################################
'''      

clp2 = [[0,0.75,1],
       [0,0.5,1],
       [0,0,1],
       [0,0,0.5],
       [0.8,0.8,0.8],
       [0.6,0.6,0.6],
       [0.4,0.4,0.4],
       [0.2,0.2,0.2],
       [0,0.5,0.5],
       [1,0.5,1],
       [1,0,1]]

fs = 12
ts = 9

# fluorescence
plt.figure(1)

plt.subplot(3,5,1)
plt.plot(t-ts,DDn[:,0]*100,color=clp2[0],linewidth=2,alpha=0.25)
plt.plot(t-ts,DDn[:,1]*100,color=clp2[0],linewidth=2)
plt.plot(t-ts,DDn[:,2]*100,color=clp2[1],linewidth=2,alpha=0.25)
plt.plot(t-ts,DDn[:,3]*100,color=clp2[1],linewidth=2)
plt.plot(t-ts,DDn[:,4]*100,color=clp2[2],linewidth=2,alpha=0.25)
plt.plot(t-ts,DDn[:,5]*100,color=clp2[2],linewidth=2)
plt.plot(t-ts,DDn[:,6]*100,color=clp2[3],linewidth=2,alpha=0.25)
plt.plot(t-ts,DDn[:,7]*100,color=clp2[3],linewidth=2)
plt.xticks(fontsize=fs)
plt.yticks(fontsize=fs)
plt.ylim(-10,110)
plt.xlim(0,120)
ax1 = plt.gca()
ax1.xaxis.set_tick_params(which='both',size=3,width=1,direction='in',top='on')
#ax1.xaxis.set_major_locator(MaxNLocator(integer=True))
ax1.yaxis.set_tick_params(which='both',size=3,width=1,direction='in',right='on')
plt.xlabel('Time (min)',fontsize=fs)
plt.ylabel('Reacted reporter (%)',fontsize=fs)

'''
###############################################################################
Loading data (Figure S2C)
###############################################################################
'''

ws = wb.sheet_by_name('Figure S2C')

t = np.zeros(ws.nrows-3)
for r in range(3,ws.nrows):
    t[r-3] = ws.cell_value(r,0)
    
DDn = np.zeros([ws.nrows-3,int(ws.ncols/2)-1])

for c in range(1,int(ws.ncols/2)):
    for r in range(3,ws.nrows):
        DDn[r-3,c-1] = ws.cell_value(r,c)

'''
###############################################################################
PLOTTING DATA (Figure S2C)
###############################################################################
'''

clp = [[0.7,0.7,0.7],
       [0.5,0.5,0.5],
       [0.3,0.3,0.3],
       [0.8,0.8,0.8],
       [0.2,0.2,0.2],
       [0.4,0.4,0.4],
       [0.6,0.6,0.6],
       [0.8,0.8,0.8]]

clp2 = [[0,0.7,0.7],
       [0,0.5,0.5],
       [0,0.3,0.3],
       [0,0.5,1],
       [0,0,0.33],
       [0,0,0.66],
       [0,0,1],
       [0,0.5,1]]

clp3 = [[0.5,0.5,0.5],
       [0,0,0.5]]

fs = 12
ts = 18

# fluorescence
plt.figure(1)

x = [0,2,4]
for n in range(len(x)):    
    plt.subplot(3,5,3)
    plt.plot(t-ts,DDn[:,x[n]]*100,color=clp2[n],linewidth=2,alpha=0.25)
    plt.plot(t-ts,DDn[:,x[n]+1]*100,color=clp2[n],linewidth=2)
    plt.xticks(fontsize=fs)
    plt.yticks(fontsize=fs)
    plt.ylim(-10,110)
    plt.xlim(0,180)
    ax1 = plt.gca()
    ax1.xaxis.set_tick_params(which='both',size=3,width=1,direction='in',top='on')
    #ax1.xaxis.set_major_locator(MaxNLocator(integer=True))
    ax1.yaxis.set_tick_params(which='both',size=3,width=1,direction='in',right='on')
    plt.xlabel('Time (min)',fontsize=fs)
    plt.ylabel('Reacted reporter (%)',fontsize=fs)
    
'''
##############################################################################
Simulations 
##############################################################################
'''

'''
Figure S2B (Single layer)
'''

t_sim = np.linspace(0,6,1001)*3600 # seconds

k_txn = 0.02

IN = [0,25]
REP_con = 500

# ksd is 1.8e3 and 6e2 to recapitulate the other two curves (less than a factor of 2 difference)
krsd = [0.5e3/1e9,1e3/1e9,2e3/1e9] # 1/nM-s

REP2 = np.array([])
REP2_l = np.array([])
REP2_h = np.array([])

for x in range(len(krsd)):
        
    for n in range(len(IN)):
        

        model = RSDs.RSD_sim() # define the model instance
        
        model.global_rate_constants(krsd=krsd[x],ktxn=k_txn)
        
        # specify DNA txn templates and reporters and concentrations
        model.molecular_species('G{1,2}',DNA_con=25,krev=5/1e9)
        model.molecular_species('R{2}',DNA_con=REP_con)
       
        model.molecular_species('I{1}',DNA_con=IN[n])
        
        # simulate the model
        model.simulate(t_sim)
        
        if x == 0:
            # pull out the species from the model solution to plot
            if len(REP2_l) == 0:
                REP2_l = model.output_concentration('S{2}')
            else:
                REP2_l = np.append([REP2_l],[model.output_concentration('S{2}')],axis=0)
        elif x == 1:
            if len(REP2) == 0:
                REP2 = model.output_concentration('S{2}')
            else:
                REP2 = np.append([REP2],[model.output_concentration('S{2}')],axis=0)
        elif x == 2:
            if len(REP2_h) == 0:
                REP2_h = model.output_concentration('S{2}')
            else:
                REP2_h = np.append([REP2_h],[model.output_concentration('S{2}')],axis=0)
             
i = 0     
plt.subplot(3,5,1)
plt.plot(model.t/60,(REP2[i]/REP_con)*100,color=[0.3,0.3,0.3],linewidth=1,linestyle='--')
plt.plot(model.t/60,(REP2[i+1]/REP_con)*100,color=[0,0,0.75],linewidth=1,linestyle='--')
plt.plot(model.t/60,(REP2_l[i]/REP_con)*100,color=[0.3,0.3,0.3],linewidth=0.75,linestyle=':',zorder=0)
plt.plot(model.t/60,(REP2_l[i+1]/REP_con)*100,color=[0,0,0.75],linewidth=.75,linestyle=':',zorder=0)
plt.plot(model.t/60,(REP2_h[i]/REP_con)*100,color=[0.3,0.3,0.3],linewidth=0.75,linestyle=':',zorder=0)
plt.plot(model.t/60,(REP2_h[i+1]/REP_con)*100,color=[0,0,0.75],linewidth=.75,linestyle=':',zorder=0)
plt.fill_between(model.t/60,(REP2_h[i+1]/REP_con)*100,(REP2_l[i+1]/REP_con)*100,color=[0,0,1],alpha=0.1,zorder=0)

'''
Figure S2C (Multilayer)
'''

t_sim = np.linspace(0,6,1001)*3600 # seconds

k_txn = 0.011

IN = [0,50]

# ksd is 1.8e3 and 6e2 to recapitulate the other two curves (less than a factor of 2 difference)
krsd = [0.5e3/1e9,1e3/1e9,2e3/1e9] # 1/nM-s


REP2 = np.array([])
REP2_l = np.array([])
REP2_h = np.array([])

for x in range(len(krsd)):
        
    for n in range(len(IN)):
        

        model = RSDs.RSD_sim() # define the model instance
        
        model.global_rate_constants(ktxn=k_txn)
        
        # specify DNA txn templates and reporters and concentrations
        model.molecular_species('G{1,2}',DNA_con=25,krsd=krsd[x])
        model.molecular_species('G{5,1}',DNA_con=25,krsd=krsd[x])
        model.molecular_species('R{2}',DNA_con=REP_con)
       
        model.molecular_species('I{5}',DNA_con=IN[n])
        
        # simulate the model
        model.simulate(t_sim)
        
        if x == 0:
            # pull out the species from the model solution to plot
            if len(REP2_l) == 0:
                REP2_l = model.output_concentration('S{2}')
            else:
                REP2_l = np.append([REP2_l],[model.output_concentration('S{2}')],axis=0)
        elif x == 1:
            if len(REP2) == 0:
                REP2 = model.output_concentration('S{2}')
            else:
                REP2 = np.append([REP2],[model.output_concentration('S{2}')],axis=0)
        elif x == 2:
            if len(REP2_h) == 0:
                REP2_h = model.output_concentration('S{2}')
            else:
                REP2_h = np.append([REP2_h],[model.output_concentration('S{2}')],axis=0)
                
i = 0     
plt.figure(1)
plt.subplot(3,5,3)
plt.plot(model.t/60,(REP2[i]/REP_con)*100,color=[0.3,0.3,0.3],linewidth=1,linestyle='--')
plt.plot(model.t/60,(REP2[i+1]/REP_con)*100,color=[0,0,1],linewidth=1,linestyle='--')
plt.plot(model.t/60,(REP2_l[i]/REP_con)*100,color=[0.3,0.3,0.3],linewidth=0.75,linestyle=':',zorder=0)
plt.plot(model.t/60,(REP2_l[i+1]/REP_con)*100,color=[0,0,1],linewidth=.75,linestyle=':',zorder=0)
plt.plot(model.t/60,(REP2_h[i]/REP_con)*100,color=[0.3,0.3,0.3],linewidth=0.75,linestyle=':',zorder=0)
plt.plot(model.t/60,(REP2_h[i+1]/REP_con)*100,color=[0,0,1],linewidth=.75,linestyle=':',zorder=0)
plt.fill_between(model.t/60,(REP2_h[i+1]/REP_con)*100,(REP2_l[i+1]/REP_con)*100,color=[0,0,0.5],alpha=0.1,zorder=0)

    
```
