## Supplementary material for "Design approaches to expand the toolkit for building cotranscriptionally encoded RNA strand displacement circuits": Simulation Scripts: FigureS13C_ctRSD_toolkit.py

```
# -*- coding: utf-8 -*-
"""
Created on Thu Jul 28 10:12:05 2022

@author: tnm12
"""

import numpy as np
import matplotlib.pyplot as plt
from matplotlib.ticker import MaxNLocator
import openpyxl as opxl
import xlrd
xlrd.xlsx.ensure_elementtree_imported(False, None)
xlrd.xlsx.Element_has_iter = True
import scipy.integrate as spi

import sys
sys.path.insert(1,'C:\\Users\\sws5\\OneDrive - National Institute of Standards and Technology (NIST)\\ctRSD simulator and documentation\\ctRSD-simulator\\ctRSD-simulator-2.0')
import ctRSD_simulator_200 as RSDs

'''
##############################################################################
Loading data 
##############################################################################
'''

save_loc = 'C:\\Users\\sws5\\OneDrive - National Institute of Standards and Technology (NIST)\\Presentations and write ups\\Papers\\RSD technical note\\Data upload files\\'
save_name = 'Supporting File S3'

wb = xlrd.open_workbook(save_loc+save_name+'.xlsx')

ws = wb.sheet_by_name('Figure S13C')

t = np.zeros(ws.nrows-3)
for r in range(3,ws.nrows):
    t[r-3] = ws.cell_value(r,0)
    
DDn = np.zeros([ws.nrows-3,int(ws.ncols/2)-1])

for c in range(1,int(ws.ncols/2)):
    for r in range(3,ws.nrows):
        DDn[r-3,c-1] = ws.cell_value(r,c)


'''
##############################################################################
PLOTTING DATA
############################################################################
'''

clp2 = [[1,0,0],
       [0,0.5,0.9],
       [0.6,0,0.8],
       [1,0.5,0],
       [212/255,170/255,0],
       [1,0,1],
       [85/255,212/255,0],
       [160/255,90/255,44/255],
       [0,0,0]]

fs = 12
ts = 0

# Overlaid fluorescence plots
plt.figure(2)
  
plt.subplot(3,5,2)
plt.plot(t-ts,DDn[:,4]*100,color=clp2[0],linewidth=2)
plt.plot(t-ts,DDn[:,0]*100,color=clp2[0],alpha=0.25,linewidth=2)
plt.plot(t-ts,DDn[:,5]*100,color=clp2[1],linewidth=2)
plt.plot(t-ts,DDn[:,1]*100,color=clp2[1],alpha=0.25,linewidth=2)
plt.plot(t-ts,DDn[:,6]*100,color=clp2[2],linewidth=2)
plt.plot(t-ts,DDn[:,2]*100,color=clp2[2],alpha=0.25,linewidth=2)
plt.plot(t-ts,DDn[:,7]*100,color=clp2[3],linewidth=2)
plt.plot(t-ts,DDn[:,3]*100,color=clp2[3],alpha=0.25,linewidth=2)
plt.xticks(fontsize=fs)
plt.yticks(fontsize=fs)
plt.ylim(-10,110)
plt.xlim(0,200)
ax1 = plt.gca()
ax1.xaxis.set_tick_params(which='both',size=3,width=1,direction='in',top='on')
#ax1.xaxis.set_major_locator(MaxNLocator(integer=True))
ax1.yaxis.set_tick_params(which='both',size=3,width=1,direction='in',right='on')
plt.xlabel('Time (min)',fontsize=fs)
plt.ylabel('Reacted reporter (%)',fontsize=fs)

# Individual fluorescence plots
for n in range(4):
    plt.subplot(3,5,11+n)
    plt.plot(t-ts,DDn[:,n+4]*100,color=clp2[n],linewidth=2)
    plt.plot(t-ts,DDn[:,n]*100,color=clp2[n],alpha=0.25,linewidth=2)
    plt.xticks(fontsize=fs)
    plt.yticks(fontsize=fs)
    plt.ylim(-10,110)
    plt.xlim(0,200)
    ax1 = plt.gca()
    ax1.xaxis.set_tick_params(which='both',size=3,width=1,direction='in',top='on')
    #ax1.xaxis.set_major_locator(MaxNLocator(integer=True))
    ax1.yaxis.set_tick_params(which='both',size=3,width=1,direction='in',right='on')
    plt.xlabel('Time (min)',fontsize=fs)
    # if n == 0:
    plt.ylabel('Reacted reporter (%)',fontsize=fs)
    # else:
    #     ax1.yaxis.set_ticklabels([])


'''
##############################################################################
Simulations 
##############################################################################
'''


k_txn = 0.0095

REP_con = 500

ksd = 1e3/1e9 
ksd1 = 1e3/1e9 # 1/nM-s
ksd2 = 1e3/1e9 # 1/nM-s
ksd3 = 5e3/1e9 #
ksd4 = 1e3/1e9 
ksd5 = 3e2/1e9 # 1/nM-s


'''
##############################################################################
Simulations
##############################################################################
'''

t_sim = np.linspace(0,6,1001)*3600 # seconds


model = RSDs.RSD_sim() # define the model instance

model.global_rate_constants(ktxn=k_txn)

# specify molecular txn DNAlates and reporters and concentrations
model.molecular_species('G{1,2}',DNA_con=25) # thresholding gate

model.molecular_species('R{2}',DNA_con=REP_con)

IN1 = [0,50,0,0,0,0,0,0]
IN4 = [0,0,0,0,0,50,0,0]
IN5 = [0,0,0,50,0,0,0,0]
IN3 = [0,0,0,0,0,0,0,50]

i = 0

pl = [11,11,12,12,13,13,14,14]
for n in range(len(IN1)):
    
    model.molecular_species('I{1}',DNA_con=IN1[n])
    model.molecular_species('I{4}',DNA_con=IN4[n])
    model.molecular_species('I{5}',DNA_con=IN5[n])
    model.molecular_species('I{3}',DNA_con=IN3[n])
    if n > 1:
        model.molecular_species('G{5,1}',DNA_con=25,krsd=ksd5) # output gate
    if n > 3:
        model.molecular_species('G{4,5}',DNA_con=25) # output gate
    if n > 5: 
        model.molecular_species('G{3,4}',DNA_con=25,krsd=ksd3) # output gate
    
    
    # simulate the model
    model.simulate(t_sim)
    
    
    # pull out the species from the model solution to plot
    REP = model.output_concentration('S{2}')
    
    fs = 12
    
    if n == 1 or n == 3 or n == 5 or n == 7:
        plt.subplot(3,5,2)
        plt.plot(model.t/60,100*(REP/REP_con),color=clp2[i-1],linewidth=1,linestyle='--')
        plt.subplot(3,5,pl[n])
        plt.plot(model.t/60,100*(REP/REP_con),color=clp2[i-1],linewidth=1,linestyle='--')
    else: 
        plt.subplot(3,5,2)
        plt.plot(model.t/60,100*(REP/REP_con),color=clp2[i],alpha=0.25,linewidth=1,linestyle='--')
        plt.subplot(3,5,pl[n])
        plt.plot(model.t/60,100*(REP/REP_con),color=clp2[i],alpha=0.25,linewidth=1,linestyle='--')
    
        i+=1
```
