## Supplementary material for "Design approaches to expand the toolkit for building cotranscriptionally encoded RNA strand displacement circuits": Simulation Scripts: FigureS14D_ctRSD_toolkit.py

```
# -*- coding: utf-8 -*-
"""
Created on Thu Jul 28 10:25:45 2022

@author: tnm12
"""

import numpy as np
import matplotlib.pyplot as plt
from matplotlib.ticker import MaxNLocator
import openpyxl as opxl
import xlrd
xlrd.xlsx.ensure_elementtree_imported(False, None)
xlrd.xlsx.Element_has_iter = True
import scipy.integrate as spi

import sys
sys.path.insert(1,'C:\\Users\\sws5\\OneDrive - National Institute of Standards and Technology (NIST)\\ctRSD simulator and documentation\\ctRSD-simulator\\ctRSD-simulator-2.0')
import ctRSD_simulator_200 as RSDs

'''
##############################################################################
Loading data 
##############################################################################
'''

save_loc = 'C:\\Users\\sws5\\OneDrive - National Institute of Standards and Technology (NIST)\\Presentations and write ups\\Papers\\RSD technical note\\Data upload files\\'
save_name = 'Supporting File S3'

wb = xlrd.open_workbook(save_loc+save_name+'.xlsx')

ws = wb.sheet_by_name('Figure S14D')

t = np.zeros(ws.nrows-3)
for r in range(3,ws.nrows):
    t[r-3] = ws.cell_value(r,0)
    
DDn = np.zeros([ws.nrows-3,int(ws.ncols/2)-1])

for c in range(1,int(ws.ncols/2)):
    for r in range(3,ws.nrows):
        DDn[r-3,c-1] = ws.cell_value(r,c)

'''
##############################################################################
PLOTTING DATA
##############################################################################
'''

clp = [[0,0,1],
       [0,0,0.5],
       [0,0.25,0.75],
       [0,0.5,1]]


fs = 12
ts = 0

# fluorescence
plt.figure(1)

plt.subplot(3,5,1)
plt.plot(t-ts,DDn[:,0]*100,color=clp[0],alpha=0.25,linewidth=2)
plt.plot(t-ts,DDn[:,1]*100,color=clp[1],alpha=0.25,linewidth=2)
plt.plot(t-ts,DDn[:,2]*100,color=clp[2],alpha=0.25,linewidth=2)
plt.plot(t-ts,DDn[:,3]*100,color=clp[3],alpha=0.25,linewidth=2)
plt.plot(t-ts,DDn[:,4]*100,color=clp[0],linewidth=2)
plt.plot(t-ts,DDn[:,5]*100,color=clp[1],linewidth=2)
plt.plot(t-ts,DDn[:,6]*100,color=clp[2],linewidth=2)
plt.plot(t-ts,DDn[:,7]*100,color=clp[3],linewidth=2)
plt.xticks(fontsize=fs)
plt.yticks(fontsize=fs)
plt.ylim(-10,110)
plt.xlim(0,220)
ax1 = plt.gca()
ax1.xaxis.set_tick_params(which='both',size=3,width=1,direction='in',top='on')
#ax1.xaxis.set_major_locator(MaxNLocator(integer=True))
ax1.yaxis.set_tick_params(which='both',size=3,width=1,direction='in',right='on')
plt.xlabel('Time (min)',fontsize=fs)
plt.ylabel('Reacted reporter (%)',fontsize=fs)


'''
##############################################################################
Simulations 
##############################################################################
'''

t_sim = np.linspace(0,6,1001)*3600 # seconds

k_txn = 0.0075

IN = [0,25]
REP_con = 500

krsd1 = [1e3/1e9,1e5/1e9,1e3/1e9,1e5/1e9] # 1/nM-s
krsd2 = [1e3/1e9,1e3/1e9,1e5/1e9,1e5/1e9] # 1/nM-s


for x in range(4):
        
    for n in range(len(IN)):
        

        model = RSDs.RSD_sim() # define the model instance
        
        model.global_rate_constants(ktxn=k_txn)
        
        # specify molecular txn DNAlates and reporters and concentrations
        model.molecular_species('G{5,1}',DNA_con=25,krsd=krsd2[x])
        model.molecular_species('G{1,2}',DNA_con=25,krsd=krsd1[x])
        model.molecular_species('R{2}',DNA_con=REP_con)
       
        model.molecular_species('I{5}',DNA_con=IN[n])
        
        # simulate the model
        model.simulate(t_sim)
        
        REP2 = model.output_concentration('S{2}')
        
        plt.figure(1)  
        plt.subplot(3,5,1)
        if  n == 0:
            plt.plot(model.t/60,(REP2/REP_con)*100,color=[0.3,0.3,0.3],linewidth=1,linestyle='--')
        else:
            plt.plot(model.t/60,(REP2/REP_con)*100,color=clp[x],linewidth=1,linestyle='--')

    
```
