## Supplementary material for "Design approaches to expand the toolkit for building cotranscriptionally encoded RNA strand displacement circuits": Simulation Scripts: FigureS29C_ctRSD_toolkit.py

```
import numpy as np
import matplotlib.pyplot as plt
import xlrd

import sys
sys.path.insert(1,'C:\\Users\\sws5\\OneDrive - National Institute of Standards and Technology (NIST)\\ctRSD simulator and documentation\\ctRSD-simulator\\ctRSD-simulator-2.0')
import ctRSD_simulator_200 as RSDs

'''
##############################################################################
Loading data 
##############################################################################
'''

save_loc = 'C:\\Users\\sws5\\OneDrive - National Institute of Standards and Technology (NIST)\\Presentations and write ups\\Papers\\RSD technical note\\Data upload files\\'
save_name = 'Supporting File S3'

wb = xlrd.open_workbook(save_loc+save_name+'.xlsx')

ws = wb.sheet_by_name('Figure S29C, bio rep1')

t = np.zeros(ws.nrows-3)
for r in range(3,ws.nrows):
    t[r-3] = ws.cell_value(r,0)
    
DDn = np.zeros([ws.nrows-3,int(ws.ncols/2)-1])

for c in range(1,int(ws.ncols/2)):
    for r in range(3,ws.nrows):
        DDn[r-3,c-1] = ws.cell_value(r,c)
        
        
ws2 = wb.sheet_by_name('Figure S29C, bio rep2')

t2 = np.zeros(ws2.nrows-3)
for r in range(3,ws2.nrows):
    t2[r-3] = ws2.cell_value(r,0)
    
DDn2 = np.zeros([ws2.nrows-3,int(ws2.ncols/2)-1])

for c in range(1,int(ws2.ncols/2)):
    for r in range(3,ws2.nrows):
        DDn2[r-3,c-1] = ws2.cell_value(r,c)

'''
###############################################################################
BIO REP 1
###############################################################################
'''

fs = 12
ts = 0
tpt = 150

# computing means
G1o = [DDn[:,0]*100,DDn[:,1]*100,DDn[:,2]*100]
G1 = [DDn[:,6]*100,DDn[:,7]*100,DDn[:,8]*100]
G12o = [DDn[:,3]*100,DDn[:,4]*100,DDn[:,5]*100]
G12 = [DDn[:,9]*100,DDn[:,10]*100,DDn[:,11]*100]

G1o_mean = np.mean(G1o,axis=0)
G1_mean = np.mean(G1,axis=0)
G12o_mean = np.mean(G12o,axis=0)
G12_mean = np.mean(G12,axis=0)

# computing standard deviations
G1o_std = np.std(G1o,axis=0)
G1_std = np.std(G1,axis=0)
G12o_std = np.std(G12o,axis=0)
G12_std = np.std(G12,axis=0)

# plotting
plt.figure(1)
plt.subplot(3,5,1)
plt.errorbar(t-ts,G1o_mean,G1o_std,color=[1,0,0],alpha=0.15)
plt.errorbar(t-ts,G1_mean,G1_std,color=[1,0,0],alpha=0.15)
plt.plot(t-ts,G1o_mean,color=[1,0,0],alpha=0.25,linewidth=1)
plt.plot(t-ts,G1_mean,color=[1,0,0],linewidth=1)

plt.errorbar(t-ts,G12o_mean,G12o_std,color=[200/255,55/255,113/255],alpha=0.15)
plt.errorbar(t-ts,G12_mean,G12_std,color=[200/255,55/255,113/255],alpha=0.15)
plt.plot(t-ts,G12o_mean,color=[200/255,55/255,113/255],alpha=0.25,linewidth=1)
plt.plot(t-ts,G12_mean,color=[200/255,55/255,113/255],linewidth=1)

plt.xticks(fontsize=fs)
plt.yticks(fontsize=fs)
plt.ylim(-10,110)
plt.xlim(0,tpt)
ax1 = plt.gca()
ax1.xaxis.set_tick_params(which='both',size=3,width=1,direction='in',top='on')
#ax1.xaxis.set_major_locator(MaxNLocator(integer=True))
ax1.yaxis.set_tick_params(which='both',size=3,width=1,direction='in',right='on')
plt.xlabel('Time (min)',fontsize=fs)
plt.ylabel('Reacted reporter (%)',fontsize=fs)

'''
###############################################################################
BIO REP 2
###############################################################################
'''

DDn_one = DDn
t_one = t
DDn_two = DDn2 
t_two = t2

# rewriting so the same code can be reused below
DDn = DDn2
t = t2

fs = 12
ts = 0
tpt = 150

# computing means
G1o = [DDn[:,0]*100,DDn[:,1]*100,DDn[:,2]*100]
G1 = [DDn[:,6]*100,DDn[:,7]*100,DDn[:,8]*100]
G12o = [DDn[:,3]*100,DDn[:,4]*100,DDn[:,5]*100]
G12 = [DDn[:,9]*100,DDn[:,10]*100,DDn[:,11]*100]

G1o_mean = np.mean(G1o,axis=0)
G1_mean = np.mean(G1,axis=0)
G12o_mean = np.mean(G12o,axis=0)
G12_mean = np.mean(G12,axis=0)

# computing standard deviations
G1o_std = np.std(G1o,axis=0)
G1_std = np.std(G1,axis=0)
G12o_std = np.std(G12o,axis=0)
G12_std = np.std(G12,axis=0)

plt.subplot(3,5,3)
plt.errorbar(t-ts,G1o_mean,G1o_std,color=[1,0,0],alpha=0.15)
plt.errorbar(t-ts,G1_mean,G1_std,color=[1,0,0],alpha=0.15)
plt.plot(t-ts,G1o_mean,color=[1,0,0],alpha=0.25,linewidth=1)
plt.plot(t-ts,G1_mean,color=[1,0,0],linewidth=1)

plt.errorbar(t-ts,G12o_mean,G12o_std,color=[200/255,55/255,113/255],alpha=0.15)
plt.errorbar(t-ts,G12_mean,G12_std,color=[200/255,55/255,113/255],alpha=0.15)
plt.plot(t-ts,G12o_mean,color=[200/255,55/255,113/255],alpha=0.25,linewidth=1)
plt.plot(t-ts,G12_mean,color=[200/255,55/255,113/255],linewidth=1)

plt.xticks(fontsize=fs)
plt.yticks(fontsize=fs)
plt.ylim(-10,110)
plt.xlim(0,tpt)
ax1 = plt.gca()
ax1.xaxis.set_tick_params(which='both',size=3,width=1,direction='in',top='on')
#ax1.xaxis.set_major_locator(MaxNLocator(integer=True))
ax1.yaxis.set_tick_params(which='both',size=3,width=1,direction='in',right='on')
plt.xlabel('Time (min)',fontsize=fs)
plt.ylabel('Reacted reporter (%)',fontsize=fs)
```
