## Supplementary material for "Design approaches to expand the toolkit for building cotranscriptionally encoded RNA strand displacement circuits": Simulation Scripts: README.txt

####################
# Python scripts
####################

Plotting and simulations were conducted using Python 3.8.3 in the Spyder IDE (version 4.1.4)

Note all the filepaths in the uploaded scripts need to be updated to the paths on the user's local computer

Others have run into issues importing *.xlsx files across operating systems / Python versions
	This can usually be overcome by saving the files with a *.xls extension 
	
